# Hepatic Stellate Cell Exosomes Resolve Fibrosis in Mice Livers via Enriched Metabolic and Regenerative Signaling Molecules

**DOI:** 10.64898/2026.04.30.721862

**Authors:** Vyshnavi Bharat, Khushboo Singh, P V Anusha, Mohammed M Idris, Chaturvedula Tripura

## Abstract

**Background:** Hepatic stellate cells (HSC) are Vitamin A storing non-parenchymal cells of the liver. During injury and inflammation, HSCs are the major contributors of excessive extracellular matrix (ECM) leading to Liver Fibrosis (LF). Emerging evidence suggests a fibrosis-independent role of these cells as key regulators of liver homeostasis and liver regeneration, emphasising on the dual role of HSCs in liver. HSCs are known to secrete several growth factors through which they largely execute their functions. However, the role of secretome (exosomes) from early activated or undifferentiated HSCs in a fibrotic milieu nor its composition are completely understood.

**Methods:** LX-2 cells were cultured in low to no serum conditions and their isolated exosomes were transplanted into fibrotic severe combined immune deficient (SCID) mice livers, followed by post-transplantation analysis of the liver tissue and compared to the untreated controls. Total proteomic profiling of cell and exosomal cargo was performed using mass spectrometry and the data analysed and compared with the total HSC cell proteome.

**Results:** Significant reduction in collagen in the transplanted mice livers compared to untreated fibrotic controls was observed with both the cells and exosomes transplantation. Comparative analysis revealed distinct enrichment of proteins and signaling pathways associated with extracellular matrix regulation, cellular communication, and metabolism in exosomes. Notably, these pathways are prominently represented in the exosomal fraction, suggesting a selective packaging of functional mediators.

**Conclusion:** This study suggests the potential role of HSCs in regulating the complex liver homeostasis via exosomal network of proteins that contribute significantly to liver repair by ECM remodelling and growth factor-mediated signalling to regulate metabolism, fibrosis and liver regeneration.

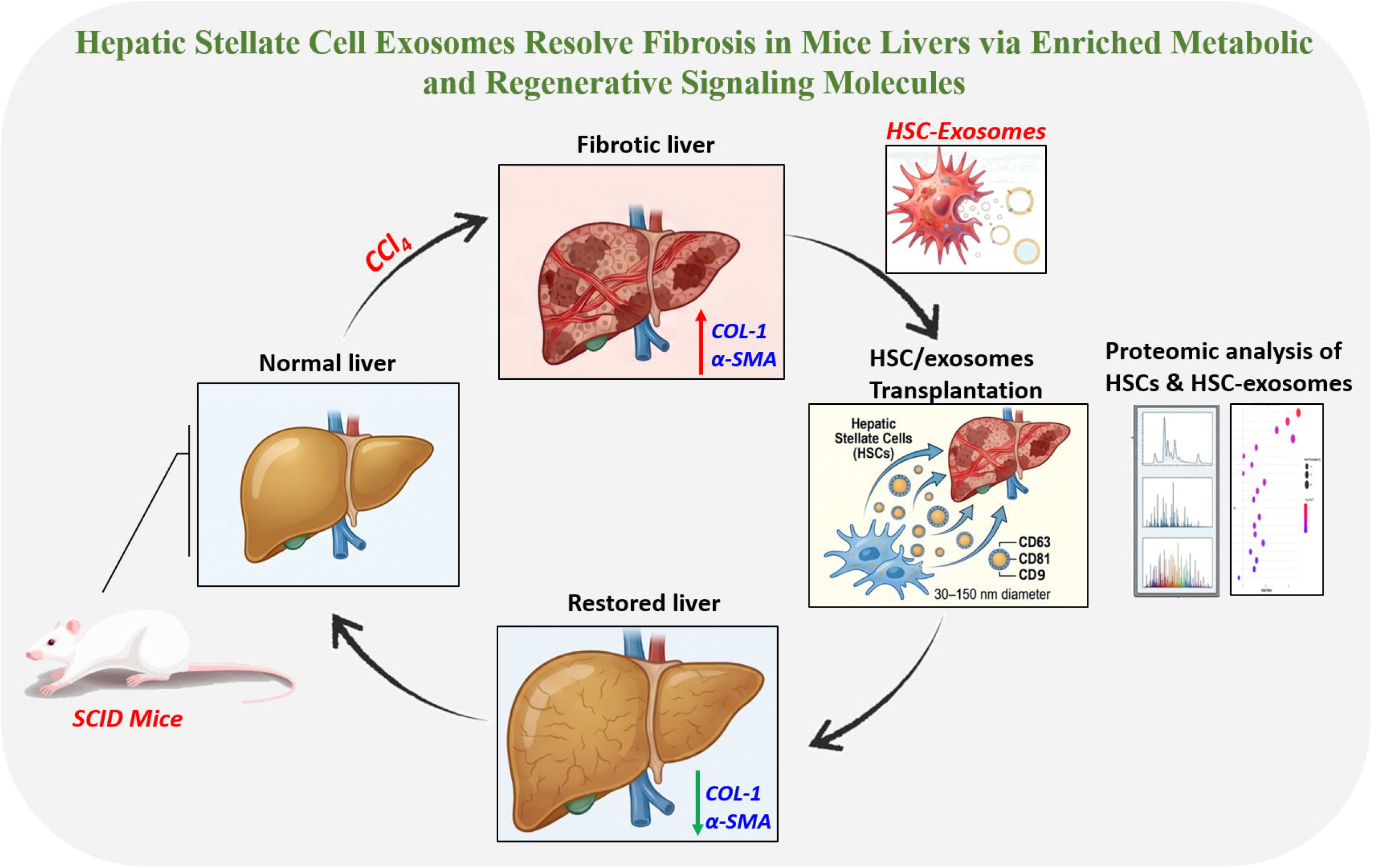

## INTRODUCTION

HSCs are mesenchymal cells constituting 5-8% of the total liver population and form a very important component of the liver. In a normal liver, these cells store retinol esters and are one of the most interactive cell types due to expression of several ligands and receptors making these ‘star cells’ unique and responsible in maintaining liver homeostasis.^1–2^ Chronic ongoing injury and inflammation leads to activation and trans-differentiation of these normally quiescent HSCs to a myofibroblast phenotype that secretes excessive collagen leading to a fibrotic scar.^3^ HSCs have been primarily associated for their role in fibrosis initiation and progression. Emerging studies reveal their importance in liver recovery and regeneration, thus these cells are considered as double edge swords playing unique role both in injury and regeneration.^4^ The direct trans-differentiation of HSCs to hepatocytes and cholangiocytes, contributing to liver regeneration post hepatectomy in a rat model was a direct evidence of these cells role as liver progenitor cells.^5^ Further, confirmation of the presence of hepatic progenitor cells within the HSC population was shown by differentiation to hepatocytes *in vitro* when cultured with appropriate growth factors.^6–7^ Also, by secreting matrix modulating enzymes such as metalloproteinases and pro-regenerative factors both quiescent and early activated HSCs are known to aid in liver regeneration.^8^

HSCs share functional similarities with Mesenchymal stromal cells (MSCs) such as immunomodulatory activity, response to external stimuli (hypoxia), release of cytokines such as VEGF, HGF, angiopoietins and insulin-like growth factors. HSCs also modulate the behaviour of other cell types such as endothelial cells and are thus considered as MSCs of the liver.^9^ Several basic and clinical investigations have previously shown that MSC-based cell therapy, including MSCs pre-treated with cytokines is effective in attenuating liver fibrosis and improving liver functions.^10–12^ These antifibrotic and pro-regenerative effects of MSCs are thought to be exerted by either suppressing proliferation or by stimulating apoptosis. However, growing evidence confirms that this antifibrotic effect of MSCs from different tissues is exerted via secretion of microvesicles that promote the reduction of fibrosis.^13–15^

Exosomes are the smallest type (30-130nm) of Extracellular vesicles (EVs) that are membrane bound bodies serving as cellular messengers between the cells. Exosomes are packed with proteins and miRNA and are vital mediators of intercellular communication in nearby and distant microenvironments.^16^ A variety of hepatic cell types produce exosomes and carry a variety of cargos enriched in proteins, lipids, mRNAs, and microRNAs (miRs).^17–19^ Most of the exosomes serve as messengers between the cells in both normal functioning and disease.

LX-2, a human hepatic stellate cell line first derived by Xu et al, is most extensively used by several researchers as an *invitro* fibrosis model.^20^ While treating with transforming growth factor beta (TGF-β) or in presence of high serum, HSCs are activated to a fibrogenic phenotype, while low serum conditions maintains these cells in quiescent state.^21–22^ Interestingly, these cells exhibit trilineage differentiation, angiogenesis potential, low immunogenicity, non-tumorigenicity and also support hematopoiesis similar to bone marrow derived mesenchymal stem cells (BM-MSCs).^23–24^ LX-2 cells are involved in tissue remodelling by secretion of matrix remodelling complex comprised of pro-matrix metalloproteinase 2 (MMP-2), membrane-type 1 matrix metalloproteinase (MT1-MMP), and TIMP-2 thus, serving as human cell model to explore the ECM degradative pathways.^20^ With their striking similarity to MSCs, combined with their unique position in the liver, it is essential to understand the non-fibrogenic and homeostatic functions of HSCs in depth.

A very recent study by Sugimoto et al. showed that the soluble fraction of HSCs enhances hepatocyte proliferation. This study clearly demonstrated the novel role of HSCs in regulating hepatocyte functions, metabolism and regeneration.^25^ With newer studies unravelling the exosomal mediated regulation involved in several cellular processes in multiple cell types, our study focused on understanding the exosomal mediated HSCs role in fibrosis reduction, liver homeostasis and regeneration.

## 2. MATERIALS AND METHODS

### 2.1 Cell culture and DiD labelling of cells

LX-2 (Hepatic Stellate cells) were routinely cultured in DMEM media supplemented with 2% fetal bovine serum (FBS), Glutamax and 1X PenStrep. Cells were seeded into 12 well plates(1*10^5^ cells/well) and were induced for trilineage differentiation once they reached a confluency of 70-80% into adipocyte (HiAdipoXL-AL521), osteocyte (HiOsteoXL-AL522) and chondrocyte (HiChondroXL-AL523) differentiation media (Hi-Media, India) for a period of 18-21 days with media change every alternate day. Differentiated cells were fixed and stained with Oil Red O, Alizarin Red, Alcian blue to confirm the differentiation. Images were acquired using the Olympus Microscope (IX83). For 1,1’-Dioctadecyl-3,3,3’,3’-Tetramethylindodicarbocyanine (DiD) labelling of cultured cells, to 1*10^6^ LX-2 cell suspension in 1ml of serum free media, 5µl of DiD cell labelling solution was added, mixed gently by pipetting and incubated for 15-20 min at 37°C. Cells were centrifuged at 1500 rpm for 5min and the cell pellet was resuspended in warm media for 10 mins to recover the cells and washed twice before final resuspension in either PBS/FACS buffer. Uniform DiD labelling was confirmed by both flow cytometry and fluorescent microscopy.

### 2.2 Flow cytometry

LX-2 cells (1*10^6^) were harvested, washed in PBS and re-suspended in FACS buffer (0.5% BSA in PBS) and incubated with fluorescence conjugated primary antibodies (Fluorescein Isothiocyanate (FITC)-CD90, Allophycocyanin-Cyanine 7 conjugate (APC Cy7)-CD73, Allophycocyanin (APC)-CD105, Phycoerythrin (PE)-CD34 (Biolegend, USA) for 30 min at 4°C, washed and analysed on Beckman Coulter machine (Gallios).

### 2.3 Immuno cytochemistry (ICC)

LX-2 cells were fixed using 4% paraformaldehyde (PFA), permeabilized with 0.5% Triton-X 100 for 15 mins at room temperature followed by blocking using 10% goat serum in PBS. Cells were stained with primary antibodies CD63 and CD81 overnight at 4°C followed by secondary antibody incubation (Alexa-488) for 1 hour at room temperature mounted with DAPI and imaged using New Zeiss Apotome microscope.

### 2.4 Exosome isolation and processing

LX-2 cells at 8*10^5^cell number were seeded onto 100 mm dishes. Upon reaching 70% confluence, cells were serum starved by replacing the media with plain media without serum and the spent medium supernatant was collected after 48 hours. Collected medium (60ml) was centrifuged at 2000g for about 30 min at 4°C and the pellet containing cell debris was discarded and the supernatant was loaded onto Amicon centrifugal filters (10 kDa cut-off) at 3200g for 30 mins. The concentrated retentate was then collected to which 0.5 volumes of Total Exosome Isolation Reagent (Thermofisher, USA) was added and mixed vigorously by pipetting and left overnight at 4°C followed by centrifugation at 10000g for 1 hour at 4°C. Supernatant was discarded and the pellet was re-suspended in 600ul of 1X PBS for further downstream analysis or for transplantation.

### 2.5 Nanoparticle Tracking Analysis (NTA)

5 µl of the exosome resuspension was then further diluted to 5ml with PBS and then loaded into the NTA instrument (NanoSight, USA), movement of the exosome particles was captured and analysed frame by frame to track the movement of individual particles and particle concentration was calculated.

### 2.6 Transmission Electron Microscopy (TEM)

5µl of the exosome sample was fixed using 2.5% glutaraldehyde for an hour at room temperature (Final volume-100 µl), and 5µl of the fixed sample was loaded onto a copper grid coated with carbon. A contrast stain (UranyLess EM stain) was then added and images were taken after 20 mins.

### 2.7 Animals

SCID mice (around 9-12 weeks) available at the CSIR-CCMB animal house facility was used after obtaining the clearance from the Institutional animal ethics committee of CSIR-CCMB IAEC (111/2022). All the animals were examined thoroughly, weighed grouped and numbered into different cages before start of the experiment. Fibrosis model in SCID mice was developed as reported earlier.^26^ Mice were injected intra-peritoneally with 0.125ml/kg carbon tetrachloride (CCl_4_) and 0.180 ml/kg CCl_4_ for a period of 6 weeks. Mice were divided into the following groups: Corn oil controls, CCl_4_ + PBS, CCl_4_ + LX-2 cells, CCl_4_ + LX-2 EV. At the end of 4 weeks, mice from the control and CCl_4_ groups (n=3) were sacrificed to confirm the development of fibrosis.

### 2.8 Transplantation of cells and exosomes into fibrotic mice

DiD labelled cells (5*10^5^) and 100μl of exosomal preparation in PBS (1.19*10^7^ particles) were transplanted/injected via tail vein into the fibrotic mice. Fibrotic mice receiving PBS served as non-transplanted controls. CCl_4_ was continued further for a period of another 2 weeks with one injection/week). After a total of 2 weeks, mice were sacrificed, serum and liver tissue were collected for further analysis.

### 2.9 Serum analysis

100μl of blood was collected by orbital puncture before the sacrifice, serum was collected. Liver function tests like AST, ALT, ALP, Albumin, protein and bilirubin content were analysed using an automated analyser as per the manufacturer’s instructions.

### 2.10 Histological staining and Immuno histo chemistry (IHC)

Paraffin embedded liver tissue blocks were sectioned and stained with Hematoxylin and Eosin (H&E). Sirius Red (SR) staining was done to image and quantify the collagen deposition using ImageJ software. Three weeks after cell transplantation, frozen liver sections were fixed using 4% PFA followed by permeabilization and mounted with DAPI. For IHC, paraffin embedded tissue sections were deparaffinised, antigen retrieved, blocked with bovine serum albumin (BSA) and stained for primary antibody overnight at 4°C. Horse radish peroxidase (HRP) polymer was added to washed sections for 30 min and 3, 3’-diaminobenzidine (DAB) substrate was used to develop colour, counter stained with Gill’s Hematoxylin, dehydrated and mounted using Dibutylphthalate Polystyrene Xylene (DPX). Images were acquired on Olympus IX83 microscope.

### 2.11 Western Blotting

Protein was isolated from stored liver tissues, using Tissue Extraction Reagent (Thermofischer scientifics). For exosomal protein, lysis buffer was added to a small amount of the exosome pellet resuspended in PBS. Total protein was quantified and 30µg of protein was separated in a 10% SDS gel and transferred onto a polyvinylidene fluoride (PVDF) membrane using the semi-dry transfer technique, blocked with 5% skimmed milk in tris-buffered saline with Tween 20 (TBST). Blots were incubated with primary antibody overnight at 4^0^C and secondary secondary antibody for 1 hour and developed using Peroxide and luminol (ratio of 1:1) for chemiluminescence.

### 2.12 Label-Free Quantitative Proteomics and analysis

Total proteins were extracted from exosome samples and cell lysate control samples using urea-based protein solubilization buffer. Protein concentrations were quantified using the Amido Black assay, ^27^ with BSA as the standard. For Label-free quantitative tandem mass spectrometry (MS/MS) analysis, 100 µg of total protein from exosome sample and cell lysates was resolved on 10% SDS-PAGE, gels were stained with Coomassie R250, destained, and excised into five fractions based on molecular weight (250-150kDa, 150-75kDa, 75-37kDa, 37-25kDa, 25-10kDa). The excised gel bands were in-gel digested with trypsin, followed by purification of the digested peptides using C18 spin columns (Thermo Scientific). The purified peptides were reconstituted in 5% acetonitrile (ACN) and 0.2% formic acid, then subjected to Liquid Chromatography Tandem Mass Spectrometry (LC-MS/MS) analysis ^28^ using an Orbitrap Velos Nano analyzer (Q-Exactive HF). The experiment was performed with 2 independent biological replicates.

### 2.13 Pathway and Functional Enrichment Analysis

For identification of peptides, the raw data was analyzed on Proteome Discoverer 2.2.3 and searched against NCBI sequence database of Homosapiens released 2022.04 and database of non-contaminants. Data were processed with a 1% false discovery rate (FDR) percolator and XCorr (Score vs. Charge) filters. Proteome Discoverer 2.2.3 label free quantification feature was used to quantify the differences in abundance between the samples. Proteins with peptide counts 2 or higher were selected for further analysis. Proteins showing more than a one-log fold change were shortlisted for further study. The mass spectrometry proteomics data have been deposited to the ProteomeXchange Consortium via the PRIDE partner repository with the dataset identifier PXD060360. Gene Ontology (GO) and Kyoto Encyclopedia of genes and genomes (KEGG) pathway enrichment analysis were performed in R version 4.4.3 using the ClusterProfiler package version 4.14.6. GO annotations were used based on the Gene Ontology database 2021. KEGG Pathway analysis was conducted using the KEGG database accessed in 2021. Differentially expressed protein analysis was performed using the Python programming language version 3.12.7. Data processing and statistical analysis were carried out using Pandas (2.2.3), NumPy (1.26.4), SciPy (1.13.1), statsmodel (0.14.2), sklearn (1.5.1), matplotlib (3.10.0) and seaborn (0.13.2). Multiple testing correction was performed using FDR (Benjamin-Hochberg) method. Comparative overlap analysis of differentially expressed proteins was performed using InteractiVenn, an interactive web based tool for Venn diagram visualization. Significantly expressed proteins were defined based on Log2FC(>1) and adjusted p-value thresholds(<0.05) and visualized using Volca NoseR Additionally, protein–protein interaction networks were generated using the Search Tool for the Retrieval of Interacting Genes/Proteins (STRING) database to visualize and explore enriched pathways and interaction clusters.

### 2.14 Statistical analysis

Statistical analysis was done using GraphPad Prism software (Version 8.4.2). Data is represented as mean ± SEM. Statistical significance between groups was assessed using unpaired or paired Student’s t-tests, or one-way ANOVA, as appropriate. Differences were considered statistically significant at p < 0.05 and highly significant at p < 0.001.

## 3. RESULTS

### 3.1 LX-2 cells in culture retain MSC characteristics

LX-2 cells that were maintained and cultured in low serum (2%) were positive for expression of MSC markers CD90 (86.5%), CD73 (97.8%), CD105 (77.06%) and negative for CD34 (Fig 1A). Successful trilineage differentiation (adipogenic, osteogenic and chondrogenic) further confirmed that the LX-2 cells continue to retain their MSC characteristics even after several rounds of culture (Fig 1B). Upon TGF-β treatment these cells have upregulated expression of Collagen-1 and α-SMA as compared to the untreated control cells that are cultured in low serum (Figure S1). For this entire study, we have used cells maintained in low serum.

**Figure 1:**
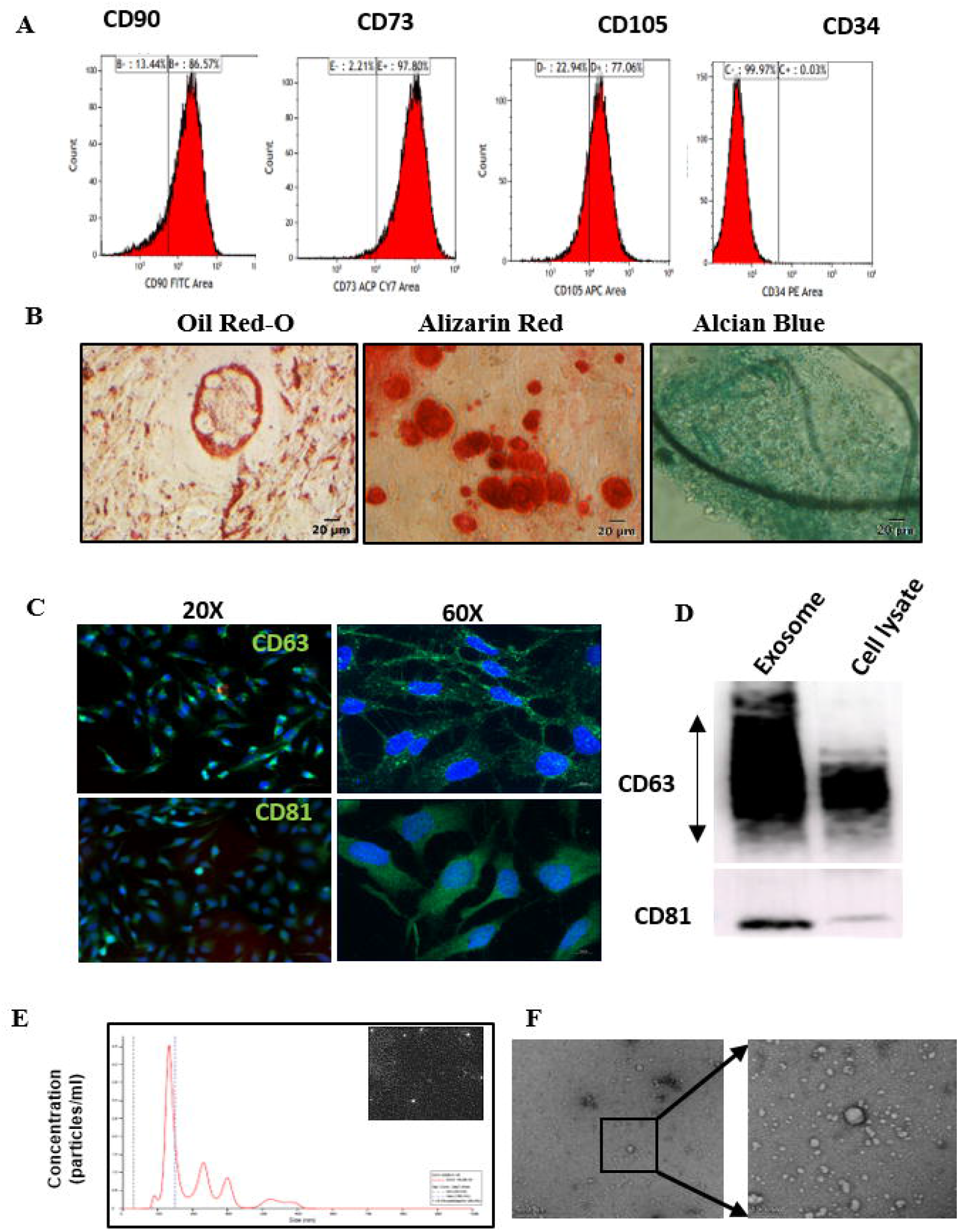
Characterization of LX-2 cells and exosomes (A) Flow cytometry profile of surface markers CD90, CD73, CD105 and CD34 (B) Confirmation of Tri-lineage differentiation by staining with Oil Red O (Adipocytes), Alizarin Red (Osteocytes) and Alcian Blue (Chondrocytes) (C) Immunocytochemistry on the cells to confirm expression of exosome specific markers, CD63 and CD81 (D) Immunoblot showing exosome specific markers-CD63 and CD81 in the exosomal preparation and in the cell lysate (E) Nanoparticle Tracking analysis (NTA) of isolated exosomes representing the concentration (number of particles/ml) vs size with the NTA image of exosomes (inset) (F) Transmission electron micrographs confirming the size of isolated exosomes as predicted from NTA analysis.

### 3.2 Characterization of exosomes by NTA, TEM and exosomal marker expression

The expression of exosomal specific markers CD81 and CD63 in the cells and exosomes was confirmed by immunocytochemistry (ICC) and immunoblot respectively (Figure 1C & D). The exosome preparation showed a mean radius distribution range with a major peak at 140 nm and the particle concentration/mL was found to be 1.19 * 10^8^ by NTA analysis (Figure 1E). TEM imaging also confirmed the size range (Figure 1F).

### 3.3 Elevated liver enzymes and histology confirmed liver fibrosis

Establishment of liver injury and fibrosis in SCID mice was confirmed after 4 weeks of CCl_4_ injection by histological and biochemical assessment of the liver tissues and serum respectively (Figure 2). Gross morphological appearance as nodulation on the liver surface and histological assessment of the livers by H&E showed infiltration of inflammatory cells in the livers of the CCl_4_ group mice (Figure 2B). The liver enzymes Serum Glutamic-Oxaloacetic transaminase (SGOT), Serum Glutamic-Pyruvic transaminase (SGPT), ALP and bilirubin levels were significantly higher in the CCl_4_-group as compared to the vehicle control group (Fig 2C). Further, α-SMA expression by IHC and Sirius red staining for collagen showed clear bridging network typical of fibrosis (Figure 2D & E). Quantification of Sirius red staining showed a significant increase in collagen deposition in the CCl_4_ group mice (Figure 2 E). Also, the total body weights of the fibrotic mice were lower than that of the vehicle control group (Figure S2 A). These results confirm that the mice livers show fibrosis and are ready for transplantation.

**Figure 2:**
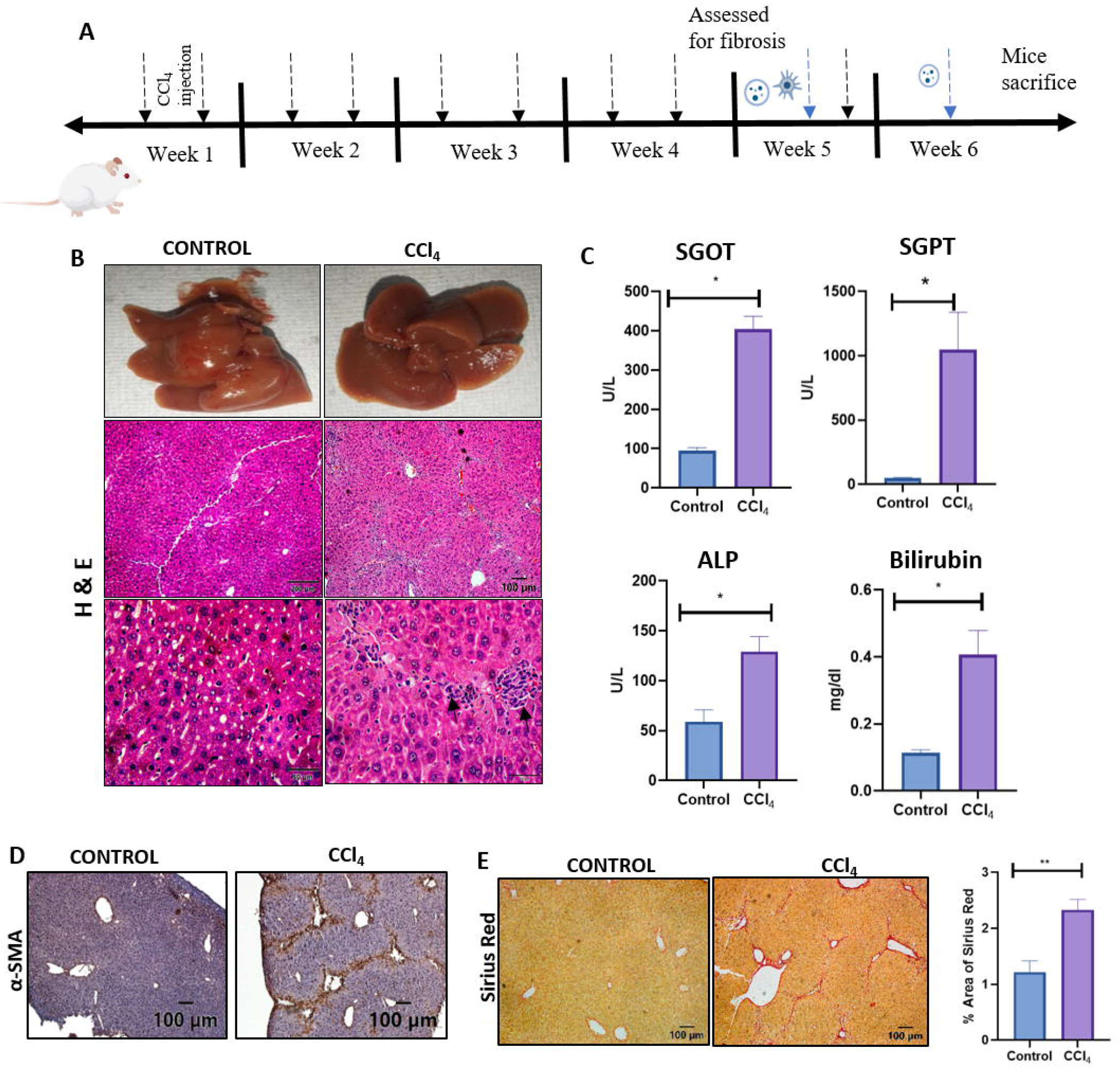
SCID mice model for liver fibrosis. (A) Timeline of the study (B) Representative macroscopic images of control and CCl_4_ mice livers and corresponding H&E images of liver sections (black arrows represent inflammatory cells)(C) Serum biochemical assays for liver function SGOT, SGPT, ALP and Bilirubin for control and CCl_4_ mice groups (n=3), analysed by unpaired t test (D) Representative images of immunohistochemical staining of α-SMA (E) Sirius Red stained liver sections and quantification (control group n=3, CCl_4_ injected group n=3; a total of 25-30 images/group were quantified), and analysed by unpaired t test. Data are presented as mean±SEM, *p□<□0.05, **p□<□0.01.

### 3.4 Significant decrease in fibrosis in the livers that received cells/exosomes as compared to the non-transplanted fibrotic livers

Cultured fresh LX-2 cells, labelled with DiD and the exosomal preparation from the same batch of cells were transplanted/injected into the fibrotic mice individually via tail vein injection. While cells were given a single dose, exosomes were injected twice with 1 week interval. In order to have an ongoing injury trigger, CCl_4_ dosage was continued with single injection/week. There was no lethality observed in the transplanted mice and body weight of the mice receiving the cells/exosomes were higher than the CCl_4_ group than the model control group (PBS alone) suggesting good recovery (Figure S2 B). Histological examination by H&E and Sirius red showed decrease in the inflammatory cells and significant reduction in fibrosis in the cells and EV group as compared to fibrosis model group suggesting contribution of both the cells/exosomes in reduced fibrosis (Figure 3A & B). α-SMA expression although not as significant as collagen levels was definitely lower than the untreated model group in both the cells and exosomes transplanted groups, (Figure 3 C&D).

**Figure 3:**
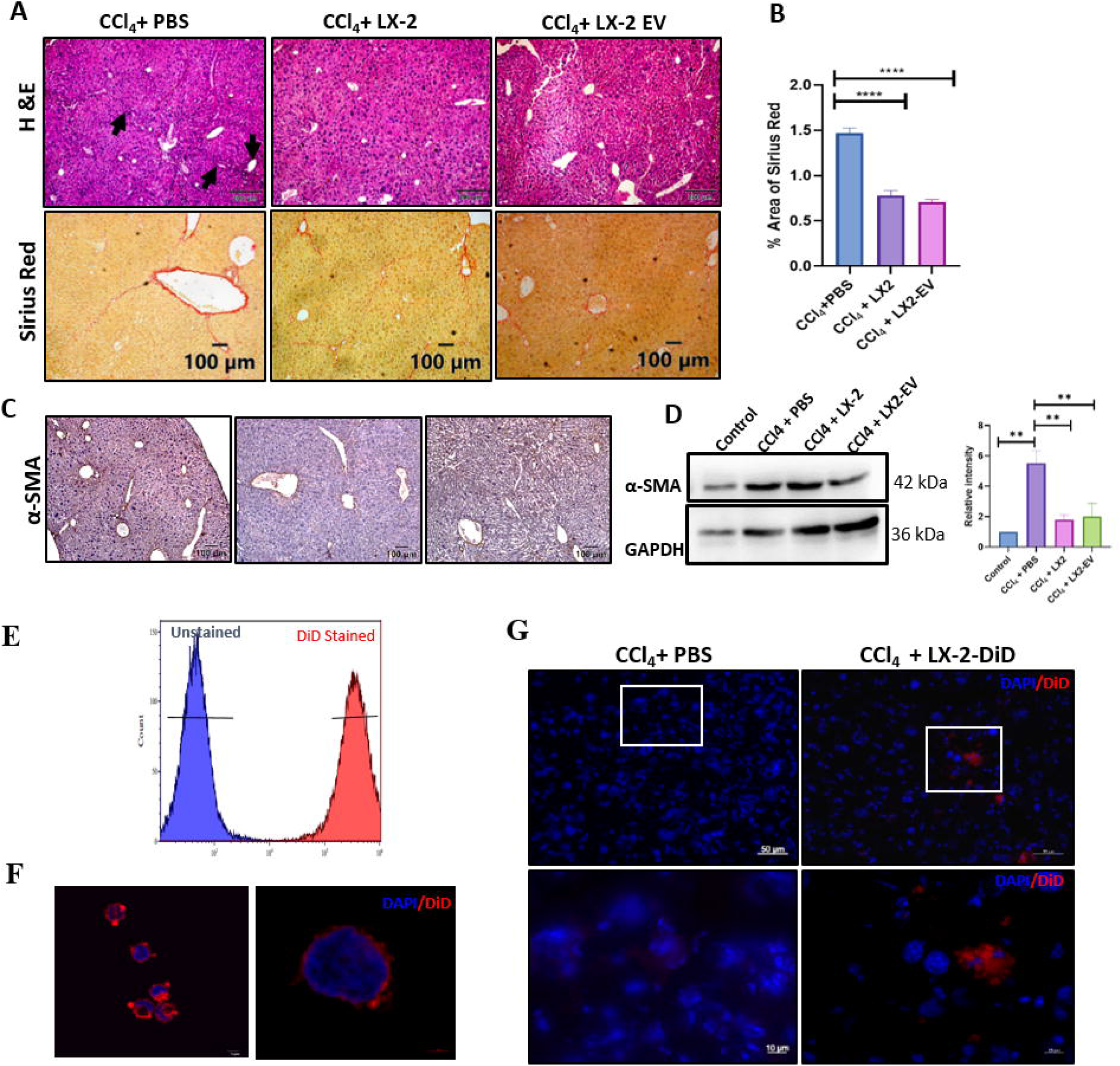
Post-transplantation/injection analysis of livers and DiD labelling and tracking of transplanted cells. (A) Representative images of H&E, SR and (B) Quantification of Sirius Red stained liver sections (n=5; a total of 25-30 images/ group were quantified) (C) α-SMA-IHC stained liver tissue sections of non-transplanted and post-transplantated with cells and exosomes (D) Western blot showing expression of α-SMA and quantified by normalisation with GAPDH. Data are presented as mean±SEM, p<0.0001. (E) Flow cytometry analysis of DiD labelled LX2 cells confirms 100% labelling (F) Representative confocal images of cells showing localization of DiD to the plasma membrane of LX-2 cells (G) Fluorescent microscopy images of cryosections of liver tissue of model group and cell transplanted group stained with DAPI and imaged showing co-localization of DiD only in the transplanted mice.

### 3.5 Presence of DiD labelled cells in the transplanted livers confirms homing of cells/exosomes to the liver

We have earlier shown lipophilic membrane dye DiD as an efficient method for tracking transplanted cells.^26^ In order to confirm that the observed effects were indeed due to homing of the cells/exosomes transplanted via the tail vein route to the liver, LX-2 cells were labelled with DiD. Flow cytometry and immuno fluorescence before transplantation confirmed uniform and 100% membrane labelling (Figure 3E & F). The presence of DiD signal only in the CCl_4_ liver tissue sections that received the labelled cells and not in the CCl_4_ livers that received PBS alone (Figure 3 G), confirmed the migration and homing of cells/exosomes to the liver via tail vein route.

### 3.6 Total proteome analysis reveals unique signatures in the exosomes

To further understand the protein signatures in the exosomes, the LX-2 cell lysates and their derived exosomes were subjected to label free quantitative LC/MS-MS analysis. A total of 2,137 proteins were identified in the cell lysate, while in the exosomes 1,546 proteins were identified, considering those with at least a one-fold log change. The exosomal proteome was compared with the known exosome databases, Exocarta and Vesiclepedia and a total of 644 common proteins were found among the three and 305 unique protein signatures were seen in our exosomes (Figure 4A). The proteome signature of the LX-2 exosomes shared a higher number of common proteins with the Exocarta database (1225) than with Vesiclepedia (660). Since, LX-2 cells exhibited properties similar to MSC cells, the exosomal proteins were compared with the MSC-Exocarta database. Out of the 1500 proteins, nearly 1000 were found to be common further confirming that MSCs and HSCs share many protein signatures attributing to the similarity in their behavior/function (Figure 4B). Between the cell lysate and exosomes of this study, 1,078 proteins were found to be common and 1059 and 468 proteins were found to be unique to the cell lysate and exosome respectively (Figure 4C). Since both the transplanted LX-2 cells and the isolated exosomes lead to reduced fibrosis, the LX-2 cells post transplantation are thought to exert their antifibrosis and regenerative effect via the exosomes. Hence, the proteomes common to the cellular and exosomal compartments were then compared. Systematic differences in protein abundance were identified that were supported by log2 fold-change, p-value distribution, log2 abundances analysis etc (Supplementary Figure S3) that support a selective and regulated mechanism governing protein incorporation into exosomes.

**Figure 4:**
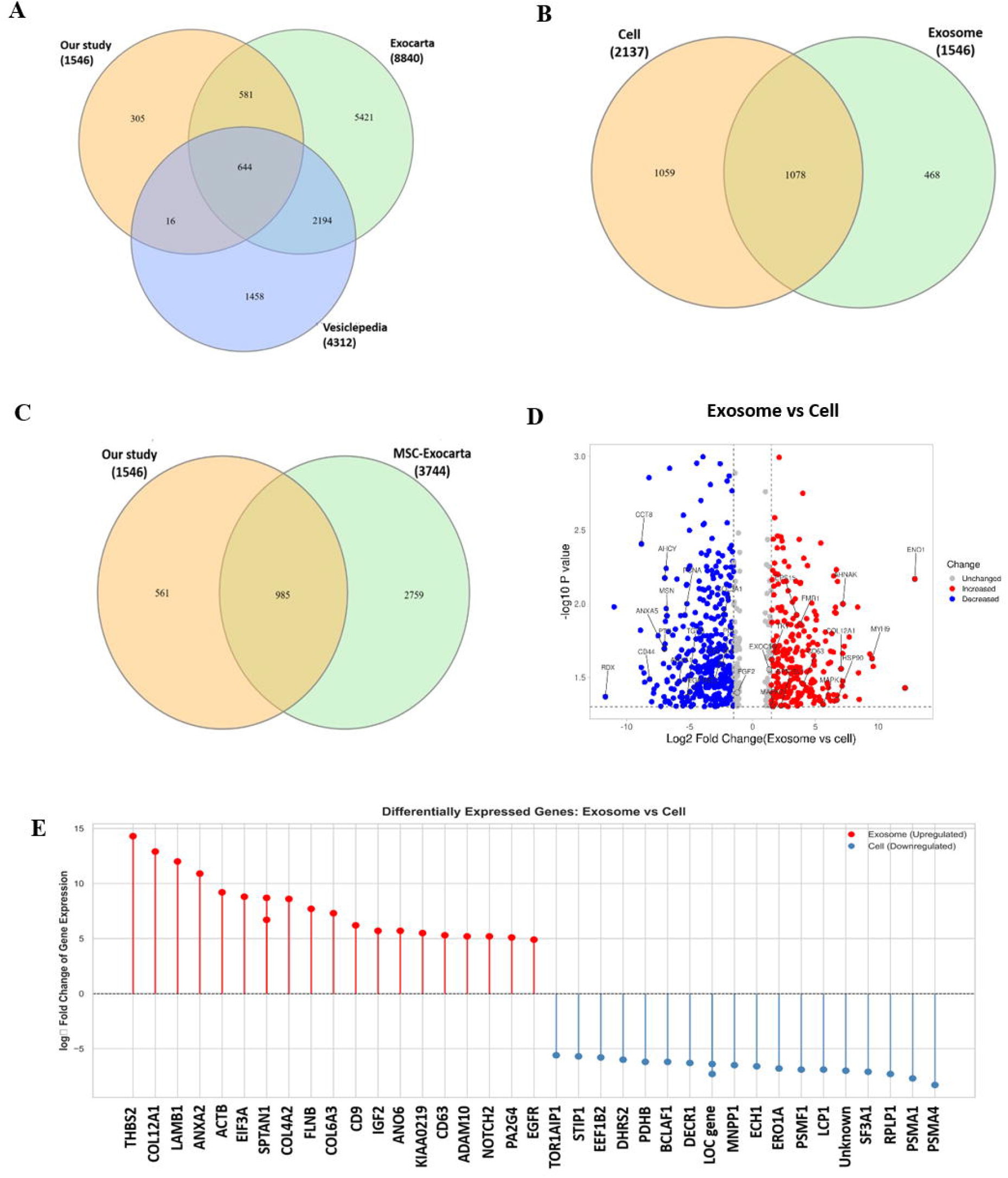
Venn diagrams showing (A) Exosomes (this study) Vs total available proteome signatures from exocarta and vesiclepedia (B) LX-2 Exosomes (this study) vs MSC-Exosomes (C) LX-2 Exosomes (this study) Vs LX-2 cell lysate (D) Volcano plot showing distribution of the up and down regulated proteins in exosomes vs LX-2 cell lysate. (E) Lollipop plot shows differentially expressed proteins between exosomal and cellular fractions. (Matplotlib, seaborn).

The volcano plot shows the selective upregulated and down regulated proteins in the exosomal fraction vs cell lysate (Figure 4D). The comparative expression analysis between exosomal and cellular fractions revealed a distinct profile of gene regulation, highlighting active molecular sorting during exosome biogenesis. The log□ fold change of the exosome vs cells indicates a total of 176 proteins upregulated, while 318 proteins downregulated. The fold change expression of the top upregulated and downregulated proteins highlight the selective enrichment of regenerative and signalling molecules within exosomes (Figure 4E). Compared to cells, exosomes exhibited significant upregulation of proteins related to extracellular matrix organization (Thrombospondin-2 (THBS2), Collagen Type XII Alpha 1 chain (COL12A1), Laminin Subunit Beta 1 (LAMB1)), cytoskeletal structure (Beta-actin (ACTB), Filamin B (FLNB)), and regenerative signaling pathways (Epidermal growth factor receptor (EGFR), A Disintegrin and metalloproteinase domain-containing protein 10 (ADAM10), Notch receptor 2 (NOTCH2)). Conversely, genes involved in mitochondrial metabolism and apoptosis regulation (Dehydrogenase/Reductase 2 (DHRS2), Pyruvate dehydrogenase E1 beta subunit (PDHB), BCL2 associated transcription factor 1 (BCLAF1), 2,4-Dienoyl-CoA Reductase 1 (DECR1), Enoyl-CoA Hydratase 1 (ECH1)) were relatively downregulated in exosomes. A selective upregulation of the exosomal markers CD9 and CD63 was observed in the exosomal preparation confirming the preparation.

The GO enrichment for cellular components showed the proteins from the exosomes to be enriched in the secretory cellular components such as membrane bound organelles, extracellular regions or organelles, vesicles, endomembrane system etc. confirming the unique nature of these proteins localized to the extracellular vesicular spaces (Figure 5A). GO enrichment for biological process showed these proteins were predominantly involved in the cell differentiation, cytoskeletal organization, tissue development, cell cycle, cell morphogenesis, response to wound healing etc. (Figure 5B). Further, the predominant molecular functions were mapped to the catalytic activity, and binding to RNA or protein and cell adhesion molecule etc. (Figure 5C). The KEGG pathway enrichment showed the top major pathways involved were in cellular stress response (Hypoxia-Inducible Factor 1 (HIF-1) signaling pathway), metabolism pathways (Glucagon and fatty acid degradation), and cellular proliferation pathways (Hippo and Wingless/Integrated (Wnt) signaling (Figure 5D). The other pathways involved in liver homeostasis such as Mitogen-Activated protein Kinase (MAPK), NOTCH, VEGF signaling were also enriched. The protein-protein interactions (PPI) networks of the interacting partners of top significant pathways such as the glucagon signaling (Figure 5E), fatty acid degradation (Figure 5F), Hippo and Wnt signaling (Figure 5 G-H) with highly significant p values suggested strong network connections.

**Figure 5:**
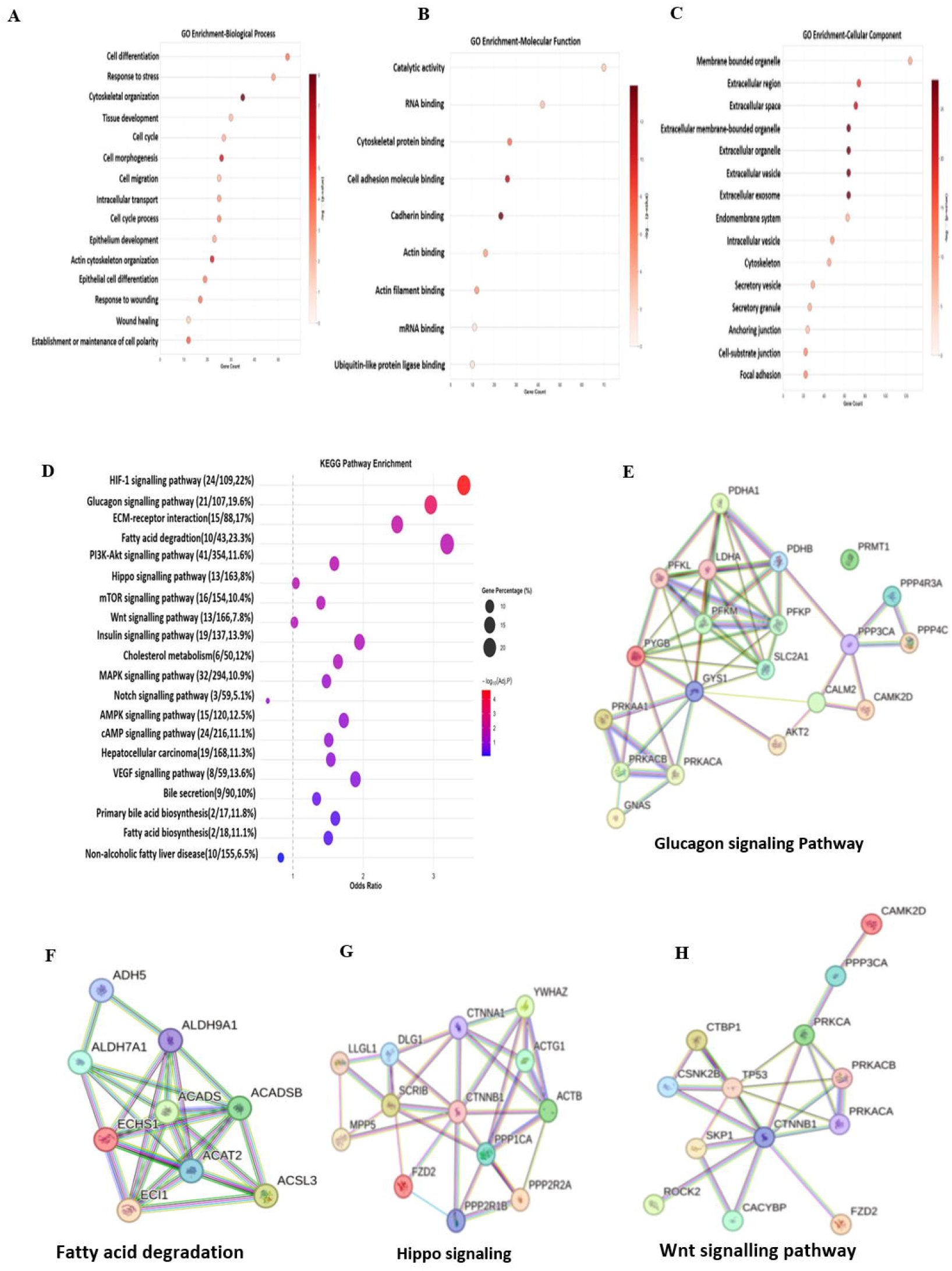
Enrichment pathways (A) GO enrichment for molecular components (B) Biological process (C) Molecular function and (D) KEGG enrichment for the top signaling pathways; Top signaling interaction pathways are shown (E) Glucagon signaling (F) Fatty acid degradation (G) Hippo signaling & (H) Wnt signaling pathway.

Further, analysis of the exosomal specific proteins with significant log2 abundance revealed the proteins to be involved majorly in liver growth (66) and regeneration (76), while a few were categorised into growth factor category (18) (Figure 6A). The log2 abundance of the high and medium proteins involved in liver growth and high and very high abundant proteins involved in liver regeneration are shown (Figure 6 B&C). The proteins in the liver regeneration category were essentially basement membrane proteins (Laminin Subunit Alpha 5 (LAMA5), Laminin Subunit Alpha 2 (LAMA2), LAMB1), matrix proteins (Fibronectin (FN1), Collagen Type VI Alpha 2 (Col6A2) etc) and matrix metalloproteinases (MMP2, MMP1) (Fig 6C). While the proteins grouped in the liver growth category were predominantly the complement proteins (Complement component 1r (C1R), Complement component 4a (C4A), Complement factor H (CFH) etc.), enzymes involved in different pathways such as the mitochondrial enzymes (Glutamic-Oxaloacetic Transaminase 2 (GOT2), Golgi enzymes Exostosin-1(EXT1), Protein Tyrosine Phosphatase 1B, Serine hydroxymethyltransferase 1 (SHMT1), Glutathione reductase (GSR1), all of which are involved in the metabolic function, cell growth and liver homeostasis.

**Figure 6:**
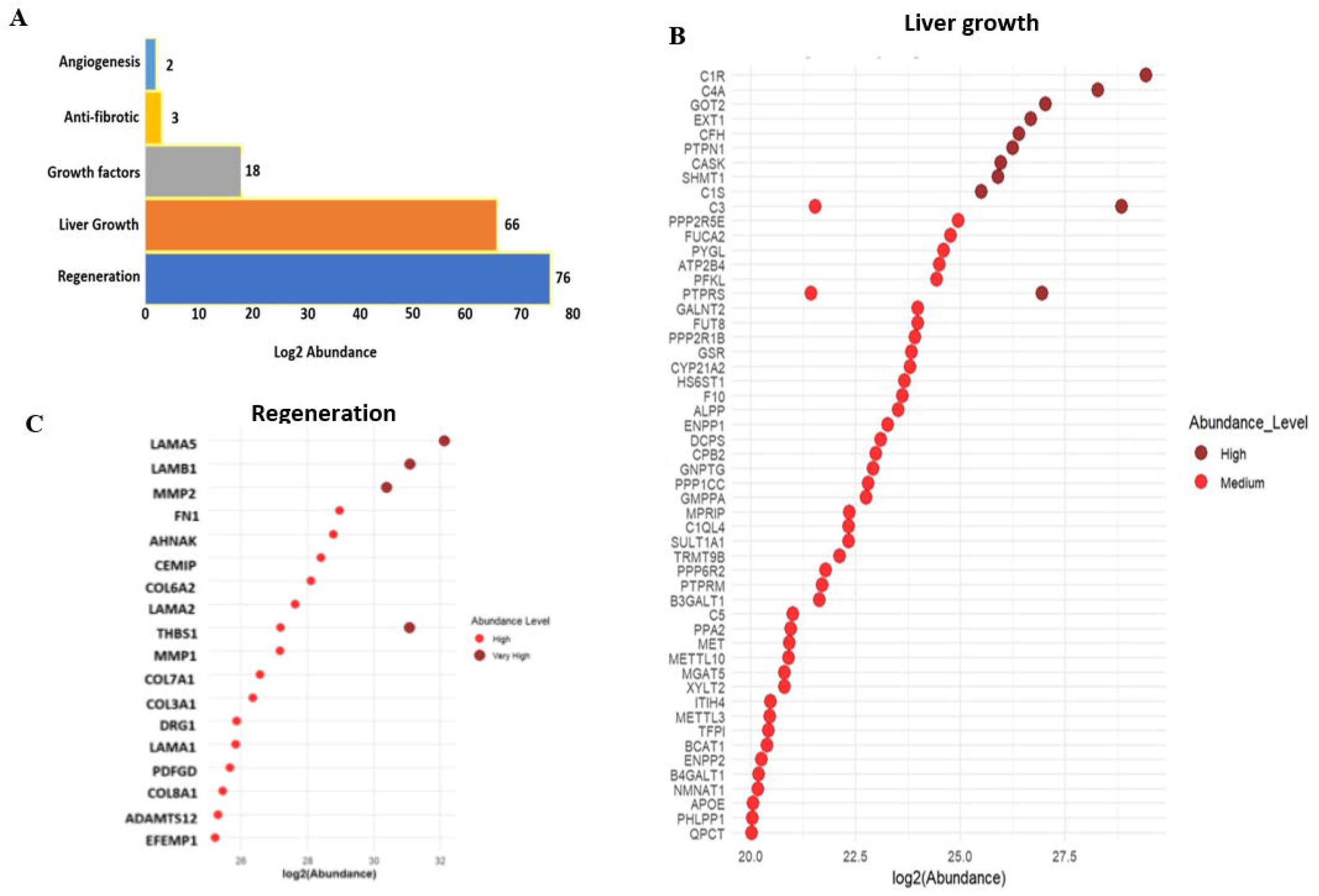
Proteins found enriched only in the exosomes (A) Log2 abundance for the proteins in the exosome fractions involved in different liver functions. Proteins in Log2 abundance scale involved in (B & C) Regeneration & Liver growth shown in log2 abundance scale.

## DISCUSSION

Hepatic stellate cells are responsible for maintaining homeostasis of the liver. Apart from the specialized function of retinol storage in normal liver and fibrogenesis during injury, HSCs significant role beyond repair includes hepatic metabolism and regeneration.^25^ In this study we show that exosomes derived from cultured HSCs (LX-2 cells) when introduced into fibrotic livers contributed to fibrosis reduction. Further proteomic profiling revealed several unique proteins involved in liver metabolism and regeneration to be enriched in the exosomes.

MSCs are immunomodulatory, anti-inflammatory, antifibrotic and are shown to efficiently alleviate liver fibrosis in murine models.^29^ Studies in rat model show that EVs derived from Adipose derived mesenchymal stem cells (AMSCs) are sufficient to suppress LF induced with CCl_4_.^30^ EVs promote the activation and proliferation of pro-inflammatory cells and suppress the secretion of anti-inflammatory cells thereby reducing fibrosis and promoting regeneration.^31–32^ With morphological and functional similarity to MSCs, HSCs are regarded as liver mesenchymal cells which could exert similar effects as MSCs on fibrosis. In line with this, studies have shown fibrosis reduction by downregulating connective tissue growth factor (CCN2) via exosomal transfer of miR-214, miR-199a-5p and the transcription factor Twist-1, between quiescent and activated HSCs.^33–34^ LF was also decreased by upregulated exosomal miR-181-5p and miR-122.^35–36^

Interestingly, despite being the primary effector cells in liver fibrosis, HSCs transplanted into healthy livers did not cause fibrosis.^37^ In another study, transplanted hepatic progenitor cells in a mouse model generated by thioacetamide (TAA) and cyclophosphamide monohydrate (CTX) settled in the injured livers and developed into functional hepatocyte like cells (HLC) and supported regeneration.^38^ These studies point to the healing and regenerative effects of HSCs when transplanted. In our study, we transplanted both the cells and exosomes. Incidentally, we found reduced fibrosis in both groups of mice receiving the cells or exosomes suggesting a common exosomal mediated mechanism to be involved.

The total proteome analysis of the cell lysate and exosomes showed that while global protein export into exosomes is limited and selective, a subset of proteins is actively enriched, supporting the hypothesis that exosomes are specifically and carefully packaged with proteins with a specified function. The combination of broad depletion, strong statistical significance, correlation with cellular abundance, and abundance-dependent variance provides a multi-faceted validation of exosomal proteome distinctiveness and highlights candidate proteins for future mechanistic studies of vesicle loading and release. The uregulation of CD9 and CD63, classical exosome markers first validated the vesicular origin of the isolated fraction. CD9 is a tetraspannin protein that is shown to participate in post injury mediated liver regeneration by promoting migration and proliferation of hepatic progenitor cells and is often used as a marker for identifying exosomes.^39–40^ CD9 also counteracts hepatic steatosis by regulating the genes involved in hepatic fatty acid synthesis and oxidation and as a mediator of glucagon receptor agonist in mitigation of liver steatosis.^40^ The proteins upregulated in the exosomes are majorly involved in extracellular matrix remodeling, cytoskeletal organization, and membrane trafficking. The proteins such as COL12A1, LAMB1, COL6A3, and FLNB are known to support tissue integrity and wound healing, suggesting that exosomal cargo may facilitate extracellular communication and regeneration.^41–43^ The key cytoskeletal and membrane-associated proteins such as ACTB, Annexin A2 (ANXA2), and Spectrin alpha-non erythrocytic 1 (SPTAN1) were also enriched, reflecting the incorporation of structural components during vesicle formation.^44^ Notably, Anoctamin6 (ANO6), ADAM10, NOTCH2, Proliferation associated 2G4 (PA2G4), and epidermal growth factor receptor (EGFR)— critical mediators of cell signaling and tissue regeneration—were significantly overexpressed in the exosomal compartment. The protease, ADAM10 is shown to as a central regulator of liver tissue homeostasis regulating the EGFR, HGF and TNFRI pathways, during liver injury and regeneration.^45–46^ These findings suggest a coordinated signaling axis (EGFR–ADAM10– NOTCH2) involved in cell proliferation and regenerative signaling, potentially influencing recipient cells via exosome-mediated communication. The presence of mitogen IGF2, additionally underscores an active role of exosomes in growth and metabolic regulation.^47^ Conversely, the downregulated cellular genes—including DHRS2, PDHB, BCLAF1, DECR1, and ECH1—are primarily associated with mitochondrial metabolism and apoptosis regulation, suggesting a cellular shift away from metabolic activity and toward vesicular export or repair processes.

Overall, this expression pattern further supports the hypothesis that exosomes act as vehicles for regenerative signaling and extracellular matrix modulation, selectively packaging signaling proteins and matrix components to coordinate intercellular communication during tissue recovery or stress response. A Very recent and interesting study, shows HSCs to regulate Wnt activity via R-spondin 3 (RSPO3) secretion, thereby controlling hepatocytes to determine the liver size, metabolic zonation in the liver, regulating liver function and regeneration.^25^ The enrichment of specific molecules involved in the major signaling pathways such as the glucagon signaling, Wnt signaling and Hippo signaling together with the ECM-receptor interactions and fatty acid degradation pathways in our exosomes is in agreement with the recent understandings of HSCs function and suggest a multidimensional role of exosomes in mitigating fibrosis and accelerating liver regeneration. Most of the studies reported earlier have shown the miRNA mediated mechanisms for fibrosis attenuation, however, for this study we have only profiled the total proteome to understand the proteome network. A further profiling of the transcriptome signatures in the exosomes would provide a greater understanding of the miRNAs involved in fibrosis regulation, liver regeneration and metabolic regulation by the exosomes.

Overall, our findings suggest that LX-2 cells derived exosomal proteins contribute significantly to liver repair by modulating key regenerative pathways, particularly through ECM remodeling, fibrotic tissue degradation, and growth factor-mediated signaling. As newer studies emerge, the role of HSCs from a fibrocentric point of view should shift to a homeostatic and regenerative view that would aid in focusing on modulating the HSC function towards regeneration rather than inhibiting or elimination mechanisms to restore liver functions.

## Abbreviations

ACN: Acetonitrile
ACTB: Beta-Actin
ADAM10: A Disintegrin And Metalloproteinase Domain-Containing Protein 10
AMSC: Adipose-Derived Mesenchymal Stem Cells
ANO6: Anoctamin6
ANXA2: Annexin A2
APC: Allophycocyanin
APC-Cy7: Allophycocyanin-Cyanine 7 conjugate
BCLAF1: BCL2 Associated Transcription Factor 1
BM-MSC: Bone Marrow-Derived Mesenchymal Stem Cells
BSA: Bovine Serum Albumin
C1R: Complement component 1R
C4A: Complement Component 4A
CCl□: Carbon Tetrachloride
CCN2: Connective tissue growth factor
CFH: Complement Factor H
COL12A1: Collagen Type XII Alpha 1 Chain
COL6A2: Collagen type VI Alpha 2
CTX: Cyclophosphamide monohydrate
DAB: 3,3’-Diaminobenzidine
DECR1: 2,4-Dienoyl-CoA Reductase 1
DHRS2: Dehydrogenase/Reductase 2
DiD: 1,1’-Dioctadecyl-3,3,3’,3’-Tetramethylindodicarbocyanine
DPX: Dibutylphthalate Polystyrene Xylene
ECH1: Enoyl-CoA Hydratase 1
ECM: Extracellular Matrix
EGFR: Epidermal Growth Factor Receptor
EV: Extracellular Vesicles
EXT1: Exostosin Glycosyltransferase 1
FBS: Fetal Bovine Serum
FDR: False Discovery Rate
FITC: Fluorescein Isothiocyanate
FLNB: Filamin B
FN1: Fibronectin 1
GO: Gene Ontology
GOT2: Glutamic-Oxaloacetic Transaminase 2 (mitochondrial AST)
GSR1: Glutathione Reductase
H&E: Hematoxylin and Eosin
HIF1: Hypoxia-Inducible Factor 1
HLC: Hepatocyte like cells
HRP: Horseradish Peroxidase
ICC: Immunocytochemistry
IHC: Immunohistochemistry
KEGG: Kyoto Encyclopedia of Genes and Genomes
LAMA2: Laminin Subunit Alpha 2
LAMA5: Laminin Subunit Alpha 5
LAMB1: Laminin Subunit Beta 1
LC-MS/MS: Liquid Chromatography–Tandem Mass Spectrometry
LF: Liver Fibrosis
LX2: Human Hepatic Stellate Cell Line (LX-2 cells)
MAPK: Mitogen-Activated Protein Kinase Pathway
miR: microRNA
MMP: Matrix Metalloproteinases
MMP2: Matrix Metalloproteinase-2
MS/MS: Tandem mass spectrometry
MSC: Mesenchymal Stem Cells
MT1-MMP: Membrane-Type 1 Matrix Metalloproteinase
NOTCH2: Notch Receptor 2
NTA: Nanoparticle Tracking Analysis
PA2G4: Proliferation associated 2G4
PDHB: Pyruvate Dehydrogenase E1 Beta Subunit
PE: Phycoerythrin
PFA: Paraformaldehyde
PPI: Protein protein interaction
PVDF: Polyvinylidene Fluoride
RSPO3: R spondin 3
SCID: Severe combined immune deficient
SDS: Sodium Dodecyl Sulfate
SGOT: Serum Glutamic-Oxaloacetic Transaminase (AST)
SGPT: Serum Glutamic-Pyruvic Transaminase (ALT)
SHMT1: Serine Hydroxymethyltransferase 1
SPTAN1: Spectrin alpha-non erythrocytic 1
SR: Sirius Red
STRING: Search Tool for the Retrieval of Interacting Genes/Proteins
TAA: Thioacetamide
TBST: Tris-Buffered Saline with Tween 20
TEM: Transmission electron microscopy
TGF-β: Transforming growth factor beta
THBS2: Thrombospondin-2
VEGF: Vascular Endothelial Growth Factor
WNT: Wingless/Integrated Signaling Pathway
α-SMA: Alpha-Smooth Muscle Actin

## DATA AVAILABILITY STATEMENT

All data generated or analysed during the study are included. All the supporting data for this research is uploladed as supplementay figures and tables 1& 2. All the proteomics raw data is uploaded in the PRIDE repository and is available as dataset identifier PXD060360 and 10.6019/PXD060360.

1. Supplementary Figs.
2. Table 1 List of Proteins unique to exosome
3. Table 2 List of proteins common between LX-2 exosomes and MSC exosomes (Exocarta)

## AUTHOR CONTRIBUTIONS

Conception and design: Chaturvedula Tripura. Performed research: Vyshnavi Bharat, PV Anusha. Collection and assembly of data: Vyshnavi Bharat, PV Anusha; Khushboo Singh. Data analysis and interpretation: Chaturvedula Tripura; Khushboo Singh, Vyshnavi Bharat and Mohammad M Idris; Manuscript writing: Chaturvedula Tripura, Vyshnavi Bharat, Khushboo Singh. Final approval of manuscript: All authors.

## ACKNOWLEDGMENTS

The authors acknowledge the kind gift of LX-2 cells from Dr. Scott Friedman. The authors thank Mr. Prashant at CSIR-CCMB for his contributions in the animal study. The authors are grateful to Director, CCMB for all the support in conducting the study.

## ETHICS APPROVAL

The animal experiments were approved by the Institutional animal ethics committee of CSIR-CCMB (IAEC 111/2022).

