## Supplementary information for "Hepatic Stellate Cell Exosomes Resolve Fibrosis in Mice Livers via Enriched Metabolic and Regenerative Signaling Molecules"

**Figure S1**

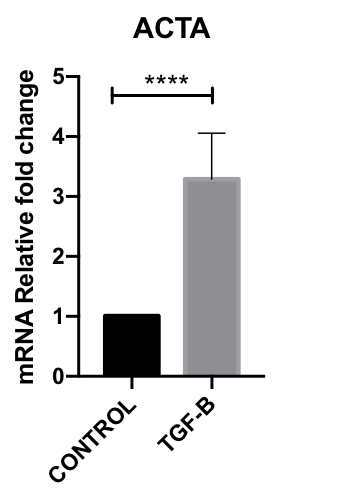

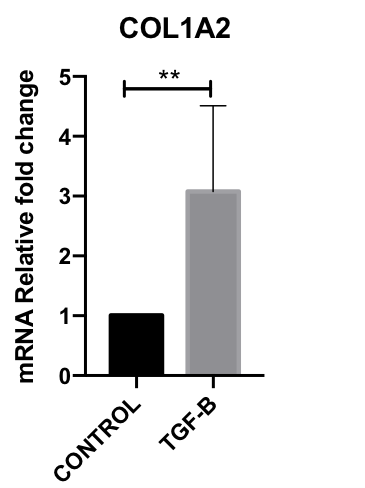

**A**

**B**

**Col1a	a-SMA**

**Figure S1: Changes in expression of col-1 and a-SMA in control and TGF-b treated cells**

LX-2 cells were serum deprived for 24h and then treated with TGF-b for 48 h. untreated cells served as controls. Cells were harvested and total RNA isolated. cDNA synthesised and the changes in the expression of the genes Col1 and a-SMA was assessed by qPCR. A significant increase in the expression of both the genes as compared to untreated controls confirmed the induction and fibrosis. Cells untreated with TGF-b and maintained in low serum (2%) continue to show lower expression of the fibrotic genes which is an indicator for undifferentiated cells.

**Figure S2**

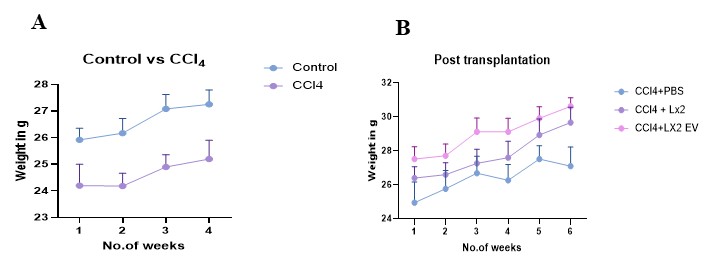

**Figure S2: Changes in the weights of mice with fibrosis induction and post transplantation.**

There is a loss of weight in the mice during fibrosis induction over a period of 4 weeks suggesting injury and weight loss (Fig A). Post transplantation a higher weight gain over a period of 6 weeks was seen in the mice receiving the EVs as compared to the mice that received only the cells, suggesting healing and recovery. The mice that received only PBS remained low in weight suggesting a lower recovery as compared to the transplantation groups.

**Figure 3**

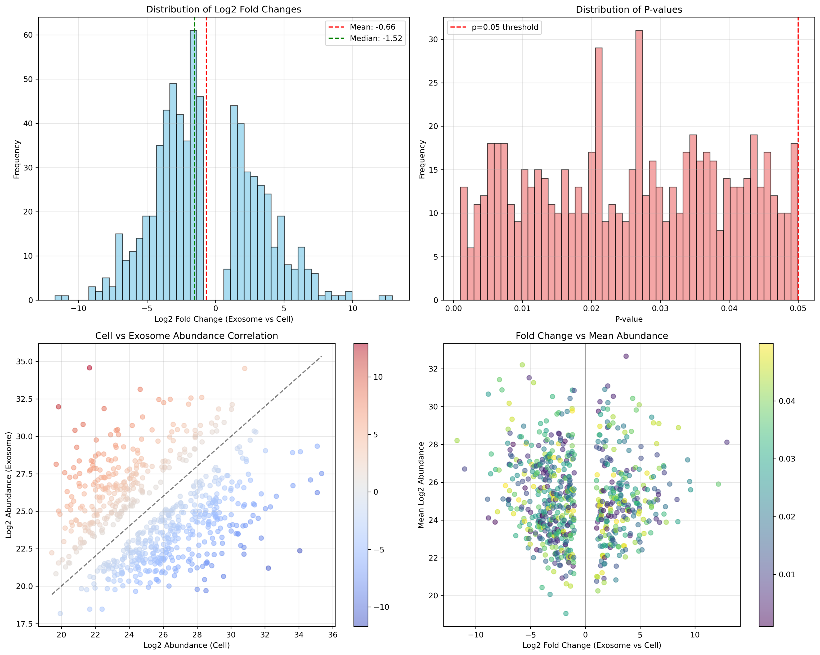

**A**

**B**

**C**

**D**

**Figure S3: Distribution of the data and correlation with the fold changes and abundances**

A) Distribution of log₂ fold changes (exosome vs cell). Vertical lines indicate median (-1.52) and mean (0.66).
B) Distribution of p-values from differential expression testing. Dashed line marks *p* = 0.05.
C) Correlation between log₂ abundance in cells and exosomes. The diagonal line represents equality.
D) MA plot of log₂ fold change versus mean log₂ abundance. Trend line (loess) highlights mean-dependent variance.
