## Supplementary Table 1 for "Hepatic Stellate Cell Exosomes Resolve Fibrosis in Mice Livers via Enriched Metabolic and Regenerative Signaling Molecules"

Table 1: Unique Proteins-Exosome

| PLG | A Chain A, Plasminogen Activator Inhibitor Type 1 |
| --- | --- |
| MMP2 | A Chain A, KDA TYPE IV COLLAGENASE |
| UROD | A Chain A, UROPORPHYRINOGEN DECARBOXYLASE |
| NMT1 | A Chain A, Glycylpeptide N-tetradecanoyltransferase 1 |
| PP1 | A Chain A, Serine/threonine protein phosphatase PP1-beta (or delta) catalytic subunit |
| MMP1 | B Chain B, Interstitial collagenase |
| PAPSS1 | A Chain A, Bifunctional 3'-phosphoadenosine 5'-phosphosulfate synthetase 1 |
| SAE1 | A Chain A, Ubiquitin-like 1 activating enzyme E1A |
| TSP1 | A Chain A, Thrombospondin 1 |
| PEPD | A Chain A, Xaa-pro Dipeptidase |
| SRI | C Chain C, Sorcin |
| MAPK2 | C Chain C, MAP kinase-activated protein kinase 2 |
| GINS3 | H Chain H, GINS complex subunit 3 |
| GLUL | E Chain E, Glutamine synthetase |
| MIF | B Chain B, Macrophage migration inhibitory factor |
| TFPI | tissue factor pathway inhibitor 2, isoform CRA_a [Homo sapiens] |
| VPS33A | B Chain B, VACUOLAR PROTEIN SORTING-ASSOCIATED PROTEIN 33A |
| SCFD1 | A Chain A, SYNTAXIN-BINDING PROTEIN 2 |
| HSP70 | A Chain A, Heat shock 70 kDa protein 1A/1B, Hsc70-interacting protein |
| SGSH | E Chain E, N-sulphoglucosamine sulphohydrolase |
| IDUA | B Chain B, Alpha-L-iduronidase |
| AP1S1 | S Chain S, AP-1 complex subunit sigma-1A |
| PFN2 | A Chain A, Profilin-1 |
| APRT | A Chain A, Adenine phosphoribosyltransferase |
| RHOA | A Chain A, Transforming protein RhoA,Rac GTPase-activating protein 1 |
| ALDH1A3 | H Chain H, Aldehyde dehydrogenase family 1 member A3 |
| PPIAL4G | C Chain C, Peptidyl-prolyl Cis-trans Isomerase A |
| PSMB2 | X Chain X, Proteasome subunit beta type-2 |
| HPCAL1 | A Chain A, Hippocalcin-like protein 1 |
| HRAS | A Chain A, GTPase HRas |
| SOD1 | H Chain H, Superoxide dismutase [Cu-Zn] |
| SNIP1 | D Chain D, Proteasome subunit beta type-8 |
| APE1 | A Chain A, DNA-(apurinic or apyrimidinic site) lyase |
| ROP7 | A Chain A, Ribonuclease P protein subunit p20 |
| CTSH | A Chain A, Pro-cathepsin H |
| C1R | B Chain B, Complement C1s subcomponent |
| SHMT2 | E Chain E, Serine hydroxymethyltransferase, mitochondrial |
| SHMT1 | A Chain A, Serine hydroxymethyltransferase, cytosolic |
| NPH1-1 | A Chain A, Tyrosine-protein phosphatase non-receptor type 1,NPH1-1 |
| GNAO1 | A Chain A, Guanine nucleotide-binding protein G(o) subunit alpha |
| RPL12 | Aq Chain Aq, 60S ribosomal protein L12 |
| CTPS2 | F Chain F, CTP synthase 2 |
| SLC12A8 | B Chain B, Solute carrier family 12 member 2 |
| DDB2 | L Chain L, DNA damage-binding protein 2 |
| ASNA1 | A Chain A, ATPase ASNA1 |
| PCSK9 | A Chain A, Proprotein convertase subtilisin/kexin type 9 |
| NAA25 | B Chain B, N-alpha-acetyltransferase 25, NatB auxiliary subunit |
| ATP6V1E1 | J Chain J, V-type proton ATPase subunit E 1 |
| CDK7 | J Chain J, Cyclin-dependent kinase 7 |
| CFI | A Chain A, iC3b1 alpha chain |
| MDM2 | CE Chain CE, Cell growth-regulating nucleolar protein |
| H2AC | n Chain n, Histone H2A type 2-A |
| RPS10 | K Chain K, 40S ribosomal protein S10 |
| VPS4B | A Chain A, Vacuolar protein sorting-associated protein 4B |
| SLC6A18 | neutral amino acid transporter [Homo sapiens] |
| RFC1 | replication factor C large subunit [Homo sapiens] |
| F2RL1 | cathepsin [Homo sapiens] |
| MMP28 | skin collagenase precursor [Homo sapiens] |
| HBB | beta-globin [Homo sapiens] |
| SCG2 | secretogranin II [Homo sapiens] |
| LOXL4 | lysyl oxidase-like protein [Homo sapiens] |
| TGFB2 | transforming growth factor-beta-2 precursor [Homo sapiens] |
| DLG2 | homolog of Drosophila discs large protein, isoform 2 [Homo sapiens] |
| C4A | complement component C4A [Homo sapiens] |
| COL6A2 | type VI collagen alpha 2 chain precursor [Homo sapiens] |
| FBXW7 | stomatin peptide [Homo sapiens] |
| MYLIP | myosin light chain 3 [Homo sapiens] |
| TGFBR1 | transmembrane receptor [Homo sapiens] |
| RPL26 | ribosomal protein L26 [Homo sapiens] |
| TNF | tumor necrosis factor [Homo sapiens] |
| PWP1 | IEF SSP 9502 [Homo sapiens] |
| RAF1 | p21-activated protein kinase [Homo sapiens] |
| PTPN13 | guanine nucleotide regulatory protein [Homo sapiens] |
| CFB | complement component C3 [Homo sapiens] |
| DNASE1L1 | DNase I-like protein [Homo sapiens] |
| ATP50 | assembly protein 50 [Homo sapiens] |
| JAG1 | transmembrane protein Jagged 1 [Homo sapiens] |
| EIF3B | Prt1 homolog [Homo sapiens] |
| XB | tenascin X [Homo sapiens] |
| FRZB | Fritz [Homo sapiens] |
| HADH | 3-hydroxyacyl-CoA dehydrogenase, isoform 2 [Homo sapiens] |
| C1S | complement component C4A, partial [Homo sapiens] |
| EXT1 | putative tumour suppressor/hereditary multiple exostoses candidate gene [Homo sapiens] |
| HMGB1 | secreted apoptosis related protein 2 [Homo sapiens] |
| PARG1 | PTPL1-associated RhoGAP [Homo sapiens] |
| COL1A1 | pro alpha 1(I) collagen [Homo sapiens] |
| SNX20 | sorting nexin 2 [Homo sapiens] |
| SEMA3F | semaphorin F homolog [Homo sapiens] |
| NRP2 | vascular endothelial cell growth factor 165 receptor/neuropilin [Homo sapiens] |
| SNX2 | sorting nexin 2 [Homo sapiens] |
| NYCO7 | antigen NY-CO-7 [Homo sapiens] |
| PPAT | glutamine PRPP amidotransferase [Homo sapiens] |
| FLNA | actin-binding protein homolog ABP-278 [Homo sapiens] |
| RAB3GAP2 | rab3-GAP regulatory domain [Homo sapiens] |
| ADAMTS | snake venom-like protease [Homo sapiens] |
| SPSB1 | spectrin SH3 domain binding protein 1 [Homo sapiens] |
| PAPPA | 2209266A plasma protein A |
| NRG1 | fusion protein, partial [Homo sapiens] |
| SQSTM1 | ubiquitin fusion-degradation 1 like protein [Homo sapiens] |
| EIF2S2 | translation initiation factor IF2 [Homo sapiens] |
| LUC7L | CGI-26 protein [Homo sapiens] |
| PGCP | blood plasma glutamate carboxypeptidase precursor [Homo sapiens] |
| HSPC027 | HSPC027 [Homo sapiens] |
| DYNC2LI1 | dynein light chain-A [Homo sapiens] |
| PDGF | secretory growth factor-like protein fallotein [Homo sapiens] |
| KIAA0291 | KIAA0291; similar rodent cytoplasmic linker protein CLIP-115 and restin [Homo sapiens] |
| CST4 | cathepsin F precursor [Homo sapiens] |
| MATR3 | matrin 3 [Homo sapiens] |
| COMMD10 | HSPC305, partial [Homo sapiens] |
| EHD3 | EH domain containing 3 [Homo sapiens] |
| PPA2 | inorganic pyrophosphatase 2 [Homo sapiens] |
| TGFBI | Transforming growth factor, beta-induced, 68kDa [Homo sapiens] |
| BASP1 | Brain abundant, membrane attached signal protein 1 [Homo sapiens] |
| CDCA8 | CDCA8 protein [Homo sapiens] |
| RPLP0 | Ribosomal protein, large, P0 [Homo sapiens] |
| TSG101 | Tumor susceptibility gene 101 [Homo sapiens] |
| LGMN | Legumain [Homo sapiens] |
| BLMH | Bleomycin hydrolase [Homo sapiens] |
| VTN | Vitronectin [Homo sapiens] |
| KPNA2 | Karyopherin alpha 2 (RAG cohort 1, importin alpha 1) [Homo sapiens] |
| TXNDC12 | Thioredoxin domain containing 12 (endoplasmic reticulum) [Homo sapiens] |
| CYCS | Cytochrome c, somatic [Homo sapiens] |
| IGFBP6 | Insulin-like growth factor binding protein 6 [Homo sapiens] |
| SRP19 | Signal recognition particle 19kDa [Homo sapiens] |
| CDC27 | CDC27 protein [Homo sapiens] |
| CD9 | CD9 molecule [Homo sapiens] |
| COL8A1 | Collagen, type VIII, alpha 1 [Homo sapiens] |
| KRT6A | Keratin 6A [Homo sapiens] |
| AP2A1 | AP2A1 protein [Homo sapiens] |
| CPOX | Coproporphyrinogen oxidase [Homo sapiens] |
| NIT2 | Nitrilase family, member 2 [Homo sapiens] |
| SRPX | Sushi-repeat-containing protein, X-linked [Homo sapiens] |
| SRPX2 | Sushi-repeat-containing protein, X-linked 2 [Homo sapiens] |
| FBNL1 | Fibulin 1 [Homo sapiens] |
| CHPF | Chondroitin polymerizing factor [Homo sapiens] |
| RRP9 | Ribosomal RNA processing 9, small subunit (SSU) processome component, homolog (yeast) [Homo sapiens] |
| VASP | Vasodilator-stimulated phosphoprotein [Homo sapiens] |
| VPS45 | Vacuolar protein sorting 45 homolog (S. cerevisiae) [Homo sapiens] |
| PRPS2 | PRPS2 protein, partial [Homo sapiens] |
| PODN | Podocan [Homo sapiens] |
| C9ORF23 | Chromosome 9 open reading frame 23 [Homo sapiens] |
| MAN2B2 | Mannosidase, alpha, class 2B, member 2 [Homo sapiens] |
| RPRD1B | Regulation of nuclear pre-mRNA domain containing 1B [Homo sapiens] |
| ARSK | ARSK protein, partial [Homo sapiens] |
| CDH2 | Cadherin 2, type 1, N-cadherin (neuronal) [Homo sapiens] |
| CNNM3 | Cyclin M3 [Homo sapiens] |
| C3ORF58 | Chromosome 3 open reading frame 58 [Homo sapiens] |
| IPO5 | Importin 5 [Homo sapiens] |
| MMP19 | Matrix metallopeptidase 19 [Homo sapiens] |
| FUCA2 | Fucosidase, alpha-L- 2, plasma [Homo sapiens] |
| EHD4 | EH-domain containing 4 [Homo sapiens] |
| CPXM1 | Carboxypeptidase X (M14 family), member 1 [Homo sapiens] |
| KIAA1967 | KIAA1967 [Homo sapiens] |
| MARCKSL1 | MARCKS-like 1 [Homo sapiens] |
| VASN | Vasorin [Homo sapiens] |
| CCNY | CCNY protein, partial [Homo sapiens] |
| B4GALT5 | UDP-Gal:betaGlcNAc beta 1,4- galactosyltransferase, polypeptide 5 [Homo sapiens] |
| EIF3D | Eukaryotic translation initiation factor 3, subunit D [Homo sapiens] |
| CPZ | Carboxypeptidase Z [Homo sapiens] |
| PTPN23 | Protein tyrosine phosphatase, non-receptor type 23 [Homo sapiens] |
| NT5DC1 | 5'-nucleotidase domain containing 1 [Homo sapiens] |
| FRMPD1 | FERM and PDZ domain containing 1 [Homo sapiens] |
| GLT8D1 | Glycosyltransferase 8 domain containing 1 [Homo sapiens] |
| KIF5B | Kinesin family member 5B [Homo sapiens] |
| ANO6 | ANO6 protein [Homo sapiens] |
| BMP1 | Bone morphogenetic protein 1 [Homo sapiens] |
| USP11 | Ubiquitin specific peptidase 11 [Homo sapiens] |
| CFH | Complement factor H [Homo sapiens] |
| SMARCD2 | SMARCD2 protein [Homo sapiens] |
| SMC2 | SMC2 protein [Homo sapiens] |
| CD151 | hemidesmosomal tetraspanin CD151 [Homo sapiens] |
| MTMR12 | phosphatidylinositol-3 phosphate 3-phosphatase adaptor subunit [Homo sapiens] |
| VPS13A | chorea-acanthocytosis [Homo sapiens] |
| HMCN1 | hemicentin [Homo sapiens] |
| B3GALT6 | beta-1,3-galactosyltransferase-6 [Homo sapiens] |
| LAMA5 | laminin alpha5 chain precursor [Homo sapiens] |
| URB | URB [Homo sapiens] |
| MMP9 | matrix metalloproteinase 9 (gelatinase B, 92kD gelatinase, 92kD type IV collagenase) [Homo sapiens] |
| FBLN7 | fibulin 2 [Homo sapiens] |
| CD109 | activated T-cell marker CD109 [Homo sapiens] |
| TUBB2A | class II beta tubulin isotype [Homo sapiens] |
| PHB | prohibitin [Homo sapiens] |
| NCSG135 | NCSG135 [Homo sapiens] |
| CPNE3 | copine III-like related protein [Homo sapiens] |
| GMFB | glia maturation factor, beta [Homo sapiens] |
| RRAS | related RAS viral (r-ras) oncogene homolog [Homo sapiens] |
| RECQ1 | RecQ protein-like (DNA helicase Q1-like) [Homo sapiens] |
| NUCB1 | nucleobindin 1 [Homo sapiens] |
| OLFML1 | olfactomedin-like [Homo sapiens] |
| SELE | selectin-like protein [Homo sapiens] |
| HRNR | hornerin precursor [Homo sapiens] |
| CPIN1 | cell proliferation-inducing protein 41 [Homo sapiens] |
| RAB11B | RAB11B, member RAS oncogene family [Homo sapiens] |
| SCGF | stem cell growth factor; lymphocyte secreted C-type lectin [Homo sapiens] |
| RABA2 | RAB2, member RAS oncogene family [Homo sapiens] |
| CEP55 | centrosomal protein 55 kDa [Homo sapiens] |
| RNASET2 | ribonuclease T2 precursor [Homo sapiens] |
| ESYT2 | extended-synaptotagmin 2 [Homo sapiens] |
| CD163 | scavenger receptor cysteine-rich glycoprotein [Homo sapiens] |
| WFDC6 | testicular secretory protein Li 57 [Homo sapiens] |
| FAM22A | YWHAE/FAM22A fusion protein, partial [Homo sapiens] |
| GNS | glucosamine -6-sulfatase isoform 1, partial [Homo sapiens] |
| PROS1 | protein S isoform 2, partial [Homo sapiens] |
| PSMB9 | PSMB9 [Homo sapiens] |
| KIAA0051 | KIAA0051 [Homo sapiens] |
| SMAD3 | Smad 3 [Homo sapiens] |
| PCOLCE | type 1 procollagen C-proteinase enhancer protein [Homo sapiens] |
| KIAA0640 | KIAA0640 protein [Homo sapiens] |
| KIAA0726 | KIAA0726 protein, partial [Homo sapiens] |
| KIAA0862 | KIAA0862 protein [Homo sapiens] |
| ASY | ASY [Homo sapiens] |
| CCBE1 | KIAA1983 protein [Homo sapiens] |
| RHOQ | TC10-like Rho GTPase variant, partial [Homo sapiens] |
| MAP3K7IP1 | mitogen-activated protein kinase kinase kinase 7 interacting protein 1 isoform alpha variant, partial [Homo sapiens] |
| DDX17 | DEAD box polypeptide 17 isoform p82 variant, partial [Homo sapiens] |
| SPOCK2 | sparc/osteonectin, cwcv and kazal-like domains proteoglycan precursor variant, partial [Homo sapiens] |
| AEBP1 | adipocyte enhancer binding protein 1 precursor variant, partial [Homo sapiens] |
| GLO1 | glyoxalase I variant, partial [Homo sapiens] |
| SERPING1 | Plasma protease C1 inhibitor precursor variant, partial [Homo sapiens] |
| THBS1 | thrombospondin 1 precursor variant, partial [Homo sapiens] |
| ANPEP | membrane alanine aminopeptidase precursor variant, partial [Homo sapiens] |
| CD146 | Melanoma cell adhesion molecule variant, partial [Homo sapiens] |
| PGRN | granulin variant, partial [Homo sapiens] |
| MATRIX | splicing factor 3a, subunit 3 variant, partial [Homo sapiens] |
| VIP36 | lectin, mannose-binding 2 variant, partial [Homo sapiens] |
| TXNDC9 | ATP binding protein associated with cell differentiation variant, partial [Homo sapiens] |
| RAB18 | RAB18, member RAS oncogene family variant, partial [Homo sapiens] |
| LIN7C | lin-7 homolog C variant, partial [Homo sapiens] |
| TSPAN9 | tetraspan NET-5 variant, partial [Homo sapiens] |
| BADH | p47 protein isoform a variant, partial [Homo sapiens] |
| NADPH | carbonyl reductase 3 variant, partial [Homo sapiens] |
| COMM | MURR1 variant, partial [Homo sapiens] |
| TPKR | SDC4-ROS1_S4;R32 fusion protein [Homo sapiens] |
| TBCD | beta-tubulin cofactor D [Homo sapiens] |
| TGFB1 | 1109243A transforming growth factor beta |
| LAMA1 | laminin A chain, partial [Homo sapiens] |
| VARS1 | valyl-tRNA synthetase [Homo sapiens] |
| FBN1 | fibrillin, partial [Homo sapiens] |
| TNC | human tenascin-C [Homo sapiens] |
| ACTR1A | alpha-centractin [Homo sapiens] |
| KRP | kinesin-related protein [Homo sapiens] |
| EMP3 | epithelial membrane protein-3 [Homo sapiens] |
| SEC23 | Sec23 protein [Homo sapiens] |
| PRSS11 | novel serine protease, PRSS11 [Homo sapiens] |
| NF2 | NF2 protein [Homo sapiens] |
| TKA1 | tyrosine kinase activator protein 1 (TKA-1) [Homo sapiens] |
| DKC1 | dyskerin [Homo sapiens] |
| CAF1 | unnamed protein product [Homo sapiens] |
| DHX36 | putative DExH/D RNA helicase [Homo sapiens] |
| IGFBP-5 | IGFBP5 [Homo sapiens] |
| VEGFC | VEGFC [Homo sapiens] |
| PPIH | PPIH [Homo sapiens] |
| PDGFRL | PDGFRL, partial [Homo sapiens] |
| Rab6A | RAB6A [Homo sapiens] |
| LAMA2 | laminin, alpha 2 (merosin, congenital muscular dystrophy) [Homo sapiens] |
| MTAP | methylthioadenosine phosphorylase [Homo sapiens] |
| MCM7 | MCM7 minichromosome maintenance deficient 7 (S. cerevisiae) [Homo sapiens] |
| ENPP1 | ectonucleotide pyrophosphatase/phosphodiesterase 1, isoform CRA_b, partial [Homo sapiens] |
| EVA1A | histidine triad nucleotide binding protein 3, isoform CRA_a [Homo sapiens] |
| LINJ | chromosome 10 open reading frame 119, isoform CRA_a [Homo sapiens] |
| IFI16 | interferon, gamma-inducible protein 16, isoform CRA_e [Homo sapiens] |
| GNB2 | guanine nucleotide binding protein (G protein), beta polypeptide 2-like 1, isoform CRA_c [Homo sapiens] |
| DNAJ | hCG2024613, isoform CRA_b [Homo sapiens] |
| SFRP-1 | hCG2039566, isoform CRA_b [Homo sapiens] |
| YTHDC1 | cadherin, EGF LAG seven-pass G-type receptor 2 (flamingo homolog, Drosophila) [Homo sapiens] |
| SORT1 | sortilin 1 [Homo sapiens] |
| C1QL4 | complement component 1, q subcomponent-like 4 [Homo sapiens] |
| KRT17 | keratin 17, isoform CRA_b [Homo sapiens] |
| FKBP10 | FK506 binding protein 10, 65 kDa, isoform CRA_b [Homo sapiens] |
| PDGFRB | platelet-derived growth factor receptor, beta polypeptide, isoform CRA_a [Homo sapiens] |
| FBN2 | fibrillin 2 (congenital contractural arachnodactyly), isoform CRA_a [Homo sapiens] |
| GSR | glutathione reductase, isoform CRA_c [Homo sapiens] |
| LRRFIP2 | leucine rich repeat (in FLII) interacting protein 2, isoform CRA_b [Homo sapiens] |
| LRIG1 | leucine-rich repeats and immunoglobulin-like domains 1, isoform CRA_b [Homo sapiens] |
| CD82 | CD82 antigen, isoform CRA_b [Homo sapiens] |
| PAMR1 | regeneration associated muscle protease [Homo sapiens] |
| PARVA | parvin, alpha, isoform CRA_c [Homo sapiens] |
| C19ORF10 | chromosome 19 open reading frame 10, isoform CRA_a [Homo sapiens] |
| CIRBP | cold inducible RNA binding protein, isoform CRA_b [Homo sapiens] |
| MYL1 | myosin, light polypeptide 1, alkali; skeletal, fast, isoform CRA_c [Homo sapiens] |
| FN1 | fibronectin 1, isoform CRA_j [Homo sapiens] |
| DTYMK | deoxythymidylate kinase (thymidylate kinase), isoform CRA_c [Homo sapiens] |
| CLSTN1 | calsyntenin 1, isoform CRA_a [Homo sapiens] |
| VBP1 | von Hippel-Lindau binding protein 1, isoform CRA_b [Homo sapiens] |
| FAM50A | family with sequence similarity 50, member A, isoform CRA_d [Homo sapiens] |
| C6ORF132 | hCG2016877, isoform CRA_c [Homo sapiens] |
| PRSS23 | protease, serine, 23, isoform CRA_b [Homo sapiens] |
| TOP1 | topoisomerase (DNA) I, isoform CRA_b [Homo sapiens] |
| DLGAP4 | discs, large (Drosophila) homolog-associated protein 4, isoform CRA_d [Homo sapiens] |
| TFPI2 | tissue factor pathway inhibitor 2, isoform CRA_a [Homo sapiens] |
| TBC1D23 | TBC1 domain family, member 23, isoform CRA_b [Homo sapiens] |
| NUMB | numb homolog (Drosophila), isoform CRA_d [Homo sapiens] |
| YY1 | YY1 transcription factor, isoform CRA_a, partial [Homo sapiens] |
| CALD1 | caldesmon 1, isoform CRA_a [Homo sapiens] |
| DNM2 | dynamin 2, isoform CRA_e [Homo sapiens] |
| COPE | coatomer protein complex, subunit epsilon, isoform CRA_g [Homo sapiens] |
| ITGB1 | integrin, beta 1 (fibronectin receptor, beta polypeptide, antigen CD29 includes MDF2, MSK12), isoform CRA_b [Homo sapiens] |
| DNM1L | dynamin 1-like, isoform CRA_c [Homo sapiens] |
| WNT5B | wingless-type MMTV integration site family, member 5B, isoform CRA_b [Homo sapiens] |
| CREG1 | cellular repressor of E1A-stimulated genes 1, isoform CRA_a [Homo sapiens] |
| TNFRSF11B | tumor necrosis factor receptor superfamily, member 11b (osteoprotegerin) [Homo sapiens] |
| SOD3 | superoxide dismutase 3, extracellular [Homo sapiens] |
| HS6ST1 | heparan sulfate 6-O-sulfotransferase 1, isoform CRA_a [Homo sapiens] |
| BTF3 | basic transcription factor 3, isoform CRA_b [Homo sapiens] |
| KRT18 | keratin 18, isoform CRA_a [Homo sapiens] |
| YEATS4 | YEATS domain containing 4, isoform CRA_a [Homo sapiens] |
| ERP29 | endoplasmic reticulum protein 29, isoform CRA_a [Homo sapiens] |
| OFD1 | oral-facial-digital syndrome 1, isoform CRA_d [Homo sapiens] |
| LTBP1 | latent transforming growth factor beta binding protein 1, isoform CRA_e [Homo sapiens] |
| TBCC | tubulin-specific chaperone c [Homo sapiens] |
| C6ORF108 | chromosome 6 open reading frame 108, isoform CRA_a [Homo sapiens] |
| CXCL9 | small inducible cytokine subfamily E, member 1 (endothelial monocyte-activating), isoform CRA_b [Homo sapiens] |
| SERPINE1 | SERPINE1 mRNA binding protein 1, isoform CRA_e [Homo sapiens] |
| KIAA0319 | KIAA0319-like, isoform CRA_a [Homo sapiens] |
| COL4A2 | collagen, type IV, alpha 2, isoform CRA_a [Homo sapiens] |
| DYNC1I2 | dynein, cytoplasmic 1, intermediate chain 2, isoform CRA_f [Homo sapiens] |
| PHF6 | PHD finger protein 6, isoform CRA_b [Homo sapiens] |
| C3 | complement C3 preproprotein [Homo sapiens] |
| F10 | coagulation factor X isoform 1 preproprotein [Homo sapiens] |
| CPM | carboxypeptidase M precursor [Homo sapiens] |
| DCTN4 | dynactin subunit 4 isoform a [Homo sapiens] |
| VDAC3 | voltage-dependent anion-selective channel protein 3 isoform 2 [Homo sapiens] |
| RABL6 | rab-like protein 6 isoform 3 [Homo sapiens] |
| HSPE1-MOB4 | HSPE1-MOB4 protein [Homo sapiens] |
| RAB5C | ras-related protein Rab-5C isoform b [Homo sapiens] |
| CCL5 | C-C motif chemokine 5 isoform 2 precursor [Homo sapiens] |
| FAT4 | protocadherin Fat 4 isoform 1 precursor [Homo sapiens] |
| PTPRK | receptor-type tyrosine-protein phosphatase kappa isoform c precursor [Homo sapiens] |
| RA32-3 | replication protein A 32 kDa subunit isoform 3 [Homo sapiens] |
| APOE | apolipoprotein E isoform a precursor [Homo sapiens] |
| GPRC5B | G-protein coupled receptor family C group 5 member B isoform 2 [Homo sapiens] |
| XRCC4 | DNA repair protein XRCC4 isoform 2 [Homo sapiens] |
| SEC24D | protein transport protein Sec24D isoform 2 [Homo sapiens] |
| TSKU | tsukushi isoform a precursor [Homo sapiens] |
| NRBF2 | nuclear receptor-binding protein isoform 2 [Homo sapiens] |
| JAK1 | tyrosine-protein kinase JAK1 isoform 1 [Homo sapiens] |
| CNOT7 | CCR4-NOT transcription complex subunit 7 isoform 3 [Homo sapiens] |
| WDR48 | WD repeat-containing protein 48 isoform 4 [Homo sapiens] |
| CRAT | carnitine O-acetyltransferase isoform 3 [Homo sapiens] |
| DCTN2 | dynactin subunit 2 isoform 4 [Homo sapiens] |
| SDCBP | syntenin-1 isoform 7 [Homo sapiens] |
| PACSIN2 | protein kinase C and casein kinase substrate in neurons protein 2 isoform D [Homo sapiens] |
| ADD1 | alpha-adducin isoform g [Homo sapiens] |
| SMARCB1 | SWI/SNF-related matrix-associated actin-dependent regulator of chromatin subfamily B member 1 isoform d [Homo sapiens] |
| ZC3HAV1 | zinc finger CCCH-type antiviral protein 1 isoform 3 [Homo sapiens] |
| CACNA2D3 | voltage-dependent calcium channel subunit alpha-2/delta-1 isoform 3 preproprotein [Homo sapiens] |
| KIF23 | kinesin-like protein KIF23 isoform 5 [Homo sapiens] |
| GREM1 | gremlin-1 isoform 1 precursor [Homo sapiens] |
| LARP7 | la-related protein 7 isoform 3 [Homo sapiens] |
| CPVL | probable serine carboxypeptidase CPVL isoform a precursor [Homo sapiens] |
| LIMS4 | LIM and senescent cell antigen-like-containing domain protein 4 isoform 2 [Homo sapiens] |
| NDRG1 | protein NDRG1 isoform 4 [Homo sapiens] |
| AB13BP | target of Nesh-SH3 isoform 8 precursor [Homo sapiens] |
| JAMA | junctional adhesion molecule A isoform 7 precursor [Homo sapiens] |
| ACAT1 | acetyl-CoA acetyltransferase, mitochondrial isoform a precursor [Homo sapiens] |
| CDC42BPA | serine/threonine-protein kinase MRCK alpha isoform F [Homo sapiens] |
| ITGA6 | integrin alpha-6 isoform 6 precursor [Homo sapiens] |
| PSMD14 | 26S proteasome non-ATPase regulatory subunit 14 [Homo sapiens] |
| COMMD3 | COMM domain-containing protein 3 [Homo sapiens] |
| PEF1 | peflin isoform 1 [Homo sapiens] |
| EVA1B | protein eva-1 homolog B [Homo sapiens] |
| RCN3 | reticulocalbin-3 precursor [Homo sapiens] |
| NFKB | transcription factor p65 isoform 1 [Homo sapiens] |
| TINAGL1 | tubulointerstitial nephritis antigen-like isoform 1 precursor [Homo sapiens] |
| HECTD3 | E3 ubiquitin-protein ligase HECTD3 [Homo sapiens] |
| RBM17 | splicing factor 45 [Homo sapiens] |
| CHST14 | carbohydrate sulfotransferase 14 [Homo sapiens] |
| AGRN | agrin isoform 2 precursor [Homo sapiens] |
| CARM1 | histone-arginine methyltransferase CARM1 isoform 1 [Homo sapiens] |
| PGRMC1 | RecName: Full=Membrane-associated progesterone receptor component 1; Short=mPR; AltName: Full=Dap1; AltName: Full=IZA |
| QSOX1 | RecName: Full=Sulfhydryl oxidase 1; Short=hQSOX; AltName: Full=Quiescin Q6; Flags: Precursor |
| CASK | RecName: Full=Peripheral plasma membrane protein CASK; Short=hCASK; AltName: Full=Calcium/calmodulin-dependent serine protein kinase; AltName: Full=Protein lin-2 homolog |
| DHX15 | RecName: Full=Pre-mRNA-splicing factor ATP-dependent RNA helicase DHX15; AltName: Full=ATP-dependent RNA helicase #46; AltName: Full=DEAH box protein 15 |
| LAMP1 | RecName: Full=Lysosome-associated membrane glycoprotein 1; Short=LAMP-1; Short=Lysosome-associated membrane protein 1; AltName: Full=CD107 antigen-like family member A; AltName: CD_antigen=CD107a; Flags: Precursor |
| COL18A1 | RecName: Full=Collagen alpha-1(XVIII) chain; Contains: RecName: Full=Endostatin; Contains: RecName: Full=Non-collagenous domain 1; Short=NC1; Flags: Precursor |
| RAB15 | RecName: Full=Ras-related protein Rab-15 |
| RPTPS | RecName: Full=Receptor-type tyrosine-protein phosphatase S; Short=R-PTP-S; AltName: Full=Receptor-type tyrosine-protein phosphatase sigma; Short=R-PTP-sigma; Flags: Precursor |
| H2AC21 | RecName: Full=Histone H2A type 2-B; AltName: Full=H2A-clustered histone 21 |
| MB21D2 | RecName: Full=Protein MB21D2; AltName: Full=Mab-21 domain-containing protein 2 |
| DCBLD1 | RecName: Full=Discoidin, CUB and LCCL domain-containing protein 1; Flags: Precursor |
| NPEPL1 | RecName: Full=Probable aminopeptidase NPEPL1; AltName: Full=Aminopeptidase-like 1 |
| IGSF8 | RecName: Full=Immunoglobulin superfamily member 8; Short=IgSF8; AltName: Full=CD81 partner 3; AltName: Full=Glu-Trp-Ile EWI motif-containing protein 2; Short=EWI-2; AltName: Full=Keratinocytes-associated transmembrane protein 4; Short=KCT-4; AltName: Full=LIR-D1; AltName: Full=Prostaglandin regulatory-like protein; Short=PGRL; AltName: CD_antigen=CD316; Flags: Precursor |
| PLXNA1 | RecName: Full=Plexin-A1; AltName: Full=Semaphorin receptor NOV; Flags: Precursor |
| SLIT2 | slit homolog 2 protein isoform X1 [Homo sapiens] |
| IL7R | interleukin-7 receptor subunit alpha isoform X1 [Homo sapiens] |
| LRRC17 | leucine-rich repeat-containing protein 17 isoform X1 [Homo sapiens] |
| PMPCB | mitochondrial-processing peptidase subunit beta isoform X1 [Homo sapiens] |
| CSNK1D | casein kinase I isoform X1 [Homo sapiens] |
| LIPG | endothelial lipase isoform X1 [Homo sapiens] |
| PDXK | pyridoxal kinase isoform X1 [Homo sapiens] |
| FAT1 | protocadherin Fat 1 isoform X1 [Homo sapiens] |
| EFEMP1 | EGF-containing fibulin-like extracellular matrix protein 1 isoform X1 [Homo sapiens] |
| CAMK1 | calcium/calmodulin-dependent protein kinase type 1 isoform X1 [Homo sapiens] |
| NID2 | nidogen-2 isoform X1 [Homo sapiens] |
| SART3 | squamous cell carcinoma antigen recognized by T-cells 3 isoform X1 [Homo sapiens] |
| MYOF | myoferlin isoform X1 [Homo sapiens] |
| POLB | DNA polymerase beta isoform X1 [Homo sapiens] |
| BRO1 | BRO1 domain-containing protein BROX isoform X3 [Homo sapiens] |
| UCHL5 | ubiquitin carboxyl-terminal hydrolase isozyme L5 isoform X1 [Homo sapiens] |
| RO60 Y | 60 kDa SS-A/Ro ribonucleoprotein isoform X2 [Homo sapiens] |
| GCA | grancalcin isoform X1 [Homo sapiens] |
| GDF11 | growth/differentiation factor 11 isoform X1 [Homo sapiens] |
| ITPA | inosine triphosphate pyrophosphatase isoform X1 [Homo sapiens] |
| NDRG3 | protein NDRG3 isoform X3 [Homo sapiens] |
| FHL1 | four and a half LIM domains protein 1 isoform X1 [Homo sapiens] |
| NIBAN1 | protein Niban 1 isoform X1 [Homo sapiens] |
| THBS3 | thrombospondin-3 isoform X1 [Homo sapiens] |
| TNFSF4 | tumor necrosis factor ligand superfamily member 4 isoform X1 [Homo sapiens] |
| RAB3GAP1 | rab3 GTPase-activating protein catalytic subunit isoform X1 [Homo sapiens] |
| MGAT5 | alpha-1,6-mannosylglycoprotein 6-beta-N-acetylglucosaminyltransferase A isoform X1 [Homo sapiens] |
| LPP | lipoma-preferred partner isoform X1 [Homo sapiens] |
| SEMA5A | semaphorin-5A isoform X1 [Homo sapiens] |
| LOC111313822 | MICAL-like protein 2 isoform X1 [Homo sapiens] |
| SEPTIN 7 | septin-7 isoform X1 [Homo sapiens] |
| AGO2 | protein argonaute-2 isoform X1 [Homo sapiens] |
| PPP2R4 | serine/threonine-protein phosphatase 2A activator isoform X1 [Homo sapiens] |
| EXT2 | exostosin-2 isoform X1 [Homo sapiens] |
| NEO1 | neogenin isoform X3 [Homo sapiens] |
| AP2B1 | AP-2 complex subunit beta isoform X1 [Homo sapiens] |
| KRT15 | keratin, type I cytoskeletal 15 isoform X1 [Homo sapiens] |
| SDF2 | stromal cell-derived factor 2 isoform X1 [Homo sapiens] |
| NAPG | gamma-soluble NSF attachment protein isoform X1 [Homo sapiens] |
| HSPBP1 | hsp70-binding protein 1 isoform X1 [Homo sapiens] |
| CDC37 | hsp90 co-chaperone Cdc37 isoform X1 [Homo sapiens] |
| COL4A5 | collagen alpha-5(IV) chain isoform X1 [Homo sapiens] |
| ROBO1 | roundabout homolog 1 isoform X1 [Homo sapiens] |
| GTF2F2 | general transcription factor IIF subunit 2 isoform X1 [Homo sapiens] |
| TNFAIP2 | tumor necrosis factor alpha-induced protein 2 isoform X1 [Homo sapiens] |
| PPP1CC | serine/threonine-protein phosphatase PP1-gamma catalytic subunit isoform X1 [Homo sapiens] |
| CALCOCO1 | calcium-binding and coiled-coil domain-containing protein 1 isoform X1 [Homo sapiens] |
| SEC24C | protein transport protein Sec24C isoform X1 [Homo sapiens] |
| UBR4 | E3 ubiquitin-protein ligase UBR4 isoform X1 [Homo sapiens] |
| PTPRF | receptor-type tyrosine-protein phosphatase F isoform X1 [Homo sapiens] |
| SNX24 | sorting nexin-24 isoform X1 [Homo sapiens] |
| PAM | peptidyl-glycine alpha-amidating monooxygenase isoform X1 [Homo sapiens] |
| ARL15 | ADP-ribosylation factor-like protein 15 isoform X1 [Homo sapiens] |
| TKFC | triokinase/FMN cyclase isoform X1 [Homo sapiens] |
| PGM2L1 | glucose 1,6-bisphosphate synthase isoform X1 [Homo sapiens] |
| LTBP3 | latent-transforming growth factor beta-binding protein 3 isoform X1 [Homo sapiens] |
| COL11A1 | collagen alpha-1(XI) chain isoform X1 [Homo sapiens] |
| DOCK10 | dedicator of cytokinesis protein 10 isoform X1 [Homo sapiens] |
| UPF0104 | carboxy-terminal domain RNA polymerase II polypeptide A small phosphatase 1 isoform X1 [Homo sapiens] |
| LZTFL1 | leucine zipper transcription factor-like protein 1 isoform X3 [Homo sapiens] |
| WNT5A | protein Wnt-5a isoform X1 [Homo sapiens] |
| RBJP | recombining binding protein suppressor of hairless isoform X1 [Homo sapiens] |
| ADAMTS12 | A disintegrin and metalloproteinase with thrombospondin motifs 12 isoform X1 [Homo sapiens] |
| PRIM2 | DNA primase large subunit isoform X1 [Homo sapiens] |
| PODNL1 | mRNA-capping enzyme isoform X1 [Homo sapiens] |
| WTAP | pre-mRNA-splicing regulator WTAP isoform X1 [Homo sapiens] |
| NUB1 | NEDD8 ultimate buster 1 isoform X1 [Homo sapiens] |
| LY96/MD-2 | lymphocyte antigen 96 isoform X1 [Homo sapiens] |
| FAT3 | protocadherin Fat 3 isoform X1 [Homo sapiens] |
| ARRB1 | beta-arrestin-1 isoform X1 [Homo sapiens] |
| BCAT1 | branched-chain-amino-acid aminotransferase, cytosolic isoform X1 [Homo sapiens] |
| COCH | cochlin isoform X1 [Homo sapiens] |
| FKBP3 | peptidyl-prolyl cis-trans isomerase FKBP3 isoform X1 [Homo sapiens] |
| GLCE | D-glucuronyl C5-epimerase isoform X1 [Homo sapiens] |
| TJP1 | tight junction protein ZO-1 isoform X1 [Homo sapiens] |
| TNFAIP1 | BTB/POZ domain-containing adapter for CUL3-mediated RhoA degradation protein 2 isoform X1 [Homo sapiens] |
| PSG4 | pregnancy-specific beta-1-glycoprotein 4 isoform X1 [Homo sapiens] |
| SRC | proto-oncogene tyrosine-protein kinase Src isoform X1 [Homo sapiens] |
| LOC118846801 | E3 ubiquitin-protein ligase HUWE1 isoform X1 [Homo sapiens] |
| ADD3 | gamma-adducin isoform X1 [Homo sapiens] |
| SmMYB113 | nucleobindin-2 isoform X1 [Homo sapiens] |
| CEMIP | cell migration-inducing and hyaluronan-binding protein isoform X1 [Homo sapiens] |
| RPN5A | 26S proteasome non-ATPase regulatory subunit 12 isoform X1 [Homo sapiens] |
| GTF2F1 | general transcription factor IIF subunit 1 isoform X1 [Homo sapiens] |
| NFASC | neurofascin isoform X1 [Homo sapiens] |
