## Supplementary Table 2 for "Hepatic Stellate Cell Exosomes Resolve Fibrosis in Mice Livers via Enriched Metabolic and Regenerative Signaling Molecules"

Table 2: List of proteins common between LX-2 exosomes and MSC exosomes (Exocarta)

| MSC-Exocarta | Exosome-Our study | Common |
| --- | --- | --- |
| RPL34 | PLG | SERPINB1 |
| SERPINB1 | ASL | RAF1 |
| PSMC2 | GSTP1 | GLT8D1 |
| RAF1 | PA2G4 | RAB14 |
| ACOX3 | UBB | YWHAG |
| RPL23P6 | SRP14 | GSS |
| YPEL5 | TXNL1 | CD9 |
| SKIV2L | MMP2 | XRCC5 |
| CLPTM1L | TYMS | PLXNA1 |
| VARS | TSN | GNB1 |
| GLT8D1 | DR1 | EIF3A |
| THUMPD3 | UROD | PDCD10 |
| TPRG1L | PDI | NIT2 |
| RAB14 | HORF6 | ASNA1 |
| SNX8 | NMT1 | RAB8A |
| GMDS | PP1 | ELAC2 |
| PPP6R1 | CSTB | MAPK1 |
| CUL7 | MMP1 | RAB9A |
| MYEF2 | #N/A | PSMB3 |
| SND1 | CUL2 | IPO9 |
| MPST | ISG20 | PRPS2 |
| YWHAG | PAPSS1 | RAB1B |
| GSS | SAE1 | MED1 |
| UBL3 | NUOB | SUPT16H |
| PSMF1 | PGAM1 | NMT1 |
| SMG9 | GSN | PLBD2 |
| ACOT1 | TSP1 | ALDH1A3 |
| ARF6 | GAPDH | TRMT6 |
| FMR1 | CLIC4 | ANXA5 |
| PKLR | AKR7A2 | SDCBP |
| FBXO9 | YWHAH | COL6A2 |
| ARHGAP35 | MDH | DHX9 |
| NUP155 | VPS26A | SQSTM1 |
| CD9 | SAR1A | YWHAZ |
| THOC2 | S100A13 | SNX2 |
| CITED1 | PHPT1 | KPNA2 |
| HSPB1P1 | PP2R1A | FLOT2 |
| IGHG3 | PEPD | EIF3G |
| ZDHHC5 | SRI | WNT5A |
| VPS18 | 6PGDH | USP9X |
| ILVBL | SNU13 | PDGFRB |
| NSUN2 | CALM1 | ATP1B1 |
| HPRT1 | SBDS | PSMC3 |
| XRCC5 | PRNP | UBE2M |
| MAP3K4 | UCH-L1 | LANCL1 |
| RPS4XP6 | SNX3 | MAT2A |
| SEC23IP | RAN | KRT6A |
| CLPX | RRP4 | PRKDC |
| TBC1D9B | SEC22B | BUB3 |
| USP4 | NAE1 NEDD8 | TRIP13 |
| NOP9 | HADH2 | NCSTN |
| PANX1 | DTD1 | UGGT1 |
| TBC1D10A | MAPK2 | IGFBP6 |
| ARHGEF18 | ENO1 | UGDH |
| GOLGA3 | PA2G4 | EIF3M |
| PLXNA1 | GINS3 | ANXA2 |
| TMPRSS9 | GLUL | RAC1 |
| GNB1 | ADSS2 | PRKAA1 |
| EIF3A | MAD2A | PCBP2 |
| PDCD10 | MUT | EDIL3 |
| NIT2 | RPA14 | LRRFIP2 |
| SVEP1 | LANCL1 | EIF2S2 |
| ASNA1 | BSG | STAT1 |
| SMYD3 | RP2 | UBA52 |
| UBAC1 | PGK1 | FEN1 |
| KRIT1 | SPDS | SHMT2 |
| TBL2 | TUBG1 | DLST |
| SDC2 | PMK | S100A13 |
| PSMB1 | N-ras | ATG5 |
| IL4I1 | VPS25 | PSMA2 |
| RAB8A | GNA1 | SRC |
| UACA | DCXR | HSPA13 |
| ADK | #N/A | FUBP1 |
| ELAC2 | POLD2 | RPL21 |
| MAPK1 | IDE | IDUA |
| IGL | TPP1 | CLIC1 |
| GALNT2 | CD47 | PDIA4 |
| SNAP23 | EIF4A1 | PPP1R7 |
| PDCL3 | EIF4A3 | HSPA5 |
| SYMPK | PDIA3 | HNRNPK |
| RCOR2 | UBE2K | UBA1 |
| U2AF2 | SEPHS1 | COL8A1 |
| KLHDC4 | GSN | CUL2 |
| RAB9A | GAPDH | COPE |
| CARKD | DDAH1 | PABPC4 |
| GRM2 | PRP8 | MFGE8 |
| DDX39A | MIF2 | IGF2R |
| COG7 | SDOS | TBCB |
| PSMB3 | FARSB | ATRN |
| CASP6 | MIF | HSPA4 |
| IPO9 | PRKACA | COL1A1 |
| GSK3A | AHCY | PIN1 |
| PRPS2 | ARFGAP1 | THBS2 |
| RBMX | WDR61 | SERPING1 |
| EEF1A2 | UGP1 | SLC16A3 |
| LACRT | PDCD10 | GNPDA1 |
| BRE | DCTN6 | PCNA |
| RAB1B | NSE | COMMD8 |
| MED1 | DBP | COL5A2 |
| TGFBR2 | AKR1B1 | SCARB2 |
| ACBD6 | RPL30 | TOR1AIP1 |
| ATP5H | MAP1LC3B | EIF2AK2 |
| HIST1H4C | GOT1 | ITGA3 |
| UCK2 | CAND1 | KPNB1 |
| STMN1 | PKM1 | CAPN2 |
| UBR1 | RPLP1 | CDC42BPA |
| SUPT16H | CFL1 | UBE3A |
| MPO | TFPI | DDX46 |
| MYADM | VPS33A | GBF1 |
| TRAPPC4 | ACAA2 | PPP2CA |
| PGRMC2 | CTNNA1 | FUCA2 |
| RAPH1 | SCFD1 | RPL3 |
| AARS | NQO1 | RABL6 |
| CHST12 | CSN4 | RAB13 |
| NMT1 | UBE2D2 | OSGEP |
| CSRP1 | SERPINE2 | RPS18 |
| MYLK2 | HARS1 | ILF3 |
| CAPZA2 | SERPINB1 | ZC3HAV1 |
| RPS27A | PLCB3 | CLTC |
| COL16A1 | SPR | SNX9 |
| RPP38 | PPP2CA | RPP30 |
| CHUK | NUP43 | TPI1 |
| HSPD1P6 | DARS1 | UPF1 |
| DDX6 | HSP70 | SMARCA1 |
| PLBD2 | CLIC1 | EGFR |
| ALDH1A3 | TKA | CAPN1 |
| CSAD | IDH1 | VASP |
| SAR1B | SARS1 | CD109 |
| ARMCX1 | SGSH | PPP2R5E |
| BRK1 | IDUA | SRP54 |
| TRMT6 | H2B2E | JAK1 |
| RELA | PSMD10 | RPS24 |
| FAT2 | YWHAG | IKBKE |
| ANXA5 | UBE2M | TXNL1 |
| RPL15P18 | AP1S1 | SRM |
| LIX1L | HSPA9 | HECTD3 |
| TERF1P3 | TXN | CUTA |
| PAF1 | QARS1 | TUBB |
| LAMTOR2 | UBE2D3 | ARPC5 |
| SDCBP | ATG5 | DDX39B |
| PFKFB2 | PIN1 | ECHDC1 |
| MXRA5 | RPL21 | PMPCB |
| CCDC93 | EEF2 | SDHA |
| BCAR1 | SNRPD1 | TUBG1 |
| NPC1 | DDB1 | COPA |
| COL6A2 | SNRPG | USP7 |
| C1QC | PFN2 | MYO1D |
| LPAR1 | APRT | RPN1 |
| DHX9 | GAL1 | SLC3A2 |
| SRXN1 | PRDX1 | OLA1 |
| APOA1BP | S100A6 | SLC44A2 |
| SPIRE1 | PSMA7 | PSME2 |
| SQSTM1 | RHOA | AP1M1 |
| MAN2A1 | TRMT6 | RRM1 |
| NOL6 | ALDH1A3 | MCM2 |
| ENO3 | PPIAL4G | PATL1 |
| PSMG2 | CDK2 | KRT1 |
| FMNL3 | HSPB1 | POR |
| IL10 | NR3C1 | PFN1 |
| MAGIX | HPRT | TKFC |
| MTX2 | FKBP12 | MIF |
| ZER1 | SF3B3 | PAFAH1B3 |
| TSPO | VCP | LMNB1 |
| YWHAZ | PYGB | HGS |
| CTNNB1 | TCEA1 | SORT1 |
| NECAP2 | ACP1 | FAT1 |
| NDUFV2 | ALDOA | GAPVD1 |
| SNX2 | ALDH2 | DIAPH1 |
| IL15RA | PSMA4 | TGFB2 |
| RPS18P12 | PSMA2 | EPB41L2 |
| ADCY9 | PSMB2 | TOLLIP |
| NOC4L | CDC34 | SFPQ |
| ERAP1 | ERF3A | ENPP1 |
| ERGIC1 | ERF1 | MPZL1 |
| KPNA2 | PRL1 | LTBP1 |
| GSPT1 | MAPK1 | PGAM1 |
| FLOT2 | NUDT5 | PRMT5 |
| PYCRL | VPS29 | CDC42 |
| CSF1 | RAB8A | ACAT2 |
| RPLP0P3 | RAB10 | SEC22B |
| TMCO1 | RAB1B | UBE2I |
| CLDND1 | PSMC6 | MATR3 |
| EIF3G | PSMD10 | EXT1 |
| CHMP2A | HPCAL1 | TUBB3 |
| GBP1 | IFIT2 | BASP1 |
| DUSP23 | RAN | ITCH |
| GALK1 | ATIC | VDAC1 |
| WNT5A | CCT5 | KRT17 |
| USP9X | H2BC11 | PSMC1 |
| IFT140 | PSMD7 | VASN |
| PDGFRB | RPS2 | C4A |
| VAV2 | HINT1 | AK1 |
| ATP1B1 | HRAS | PRDX4 |
| F5 | VDAC1 | ARPC5L |
| ARF5 | PRPF8 | DHRS1 |
| PGM3 | SNRPD2 | EIF3I |
| RER1 | RBBP4 | CYFIP2 |
| ITGA4 | EEF1E1 | ARHGAP18 |
| FBXO7 | NNMT | PLCG1 |
| IGFBP7 | SOD1 | EIF2B2 |
| PLXNB2 | SF1 | P4HA2 |
| HS1BP3 | SF1 | DNM1L |
| SRSF9 | SF3B1 | PSPC1 |
| PNO1 | SNRPA1 | TGFB1 |
| USP15 | YWHAB | PCMT1 |
| TARS | PMCA1 | CTSZ |
| ABTB2 | SF3B4 | RPL5 |
| LMOD1 | RRM1 | DDAH1 |
| ATP2B4 | SNIP1 | ISYNA1 |
| PSMC3 | RHEB | ITGAV |
| KCMF1 | APE1 | FLNA |
| UBE2M | NAA15 | CLDN11 |
| PARP16 | PRMT5 | RRAS |
| LANCL1 | ROP7 | SMC2 |
| MAT2A | ACTG1 | EHD3 |
| S100A4 | CTSH | PFN2 |
| KRT6A | NCBP2 | TRIM28 |
| MMS19 | RRP45 | HNRNPL |
| PRKDC | MTREX | CSTF1 |
| BUB3 | NT5C1A | PSMC5 |
| NDUFA13 | PSMA3 | DAG1 |
| TRIP13 | CALR | NCKAP1 |
| FAM129B | EIF2B2 | RPLP0 |
| NCSTN | C1R | THOC6 |
| SLFN5 | PCNA | RHEB |
| PHLDB1 | RPS6KB1 | SNAPIN |
| ACAP2 | SNRNP200 | BDH2 |
| UGGT1 | RPS9 | SERPINE2 |
| CLINT1 | RPS18 | NAPA |
| HPS5 | NELF-E | SEC24A |
| PARVG | DIS3 | TXNRD1 |
| IGFBP6 | SHMT2 | C1S |
| SLC22A4 | PPP1R7 | CDKN2A |
| UGDH | CK2 | RAP1GDS1 |
| EIF3M | MAGOHB | RAB3GAP1 |
| DHX16 | PRPF19 | FBN1 |
| FGB | RPS26 | CNP |
| ANXA2 | RPS26 | PFAS |
| PTGES2 | YKT6 | RPL7A |
| F8 | CTSL1 | SNX3 |
| SETD3 | ANXA5 | RPSA |
| TMEM55B | LARS1 | GSR |
| CCAR2 | PSME4 | PPAT |
| RAC1 | YWHAQ | ARF4 |
| KRT76 | TRIP13 | RAB10 |
| IGFBP2 | NCSTN | MAPRE1 |
| PRKAA1 | RPL38 | SCFD1 |
| HSPD1P1 | SLC16A1 | ITGA5 |
| NUP160 | SHMT1 | NCAPG |
| TUBBP1 | PSMD3 | CHPF |
| CHODL | PSMC4 | CLIC4 |
| PCBP2 | NPH1-1 | CDH2 |
| STK38 | EIF2S3 | MARCKS |
| PTPRA | EIF2S1 | ITPA |
| ACADM | GNAO1 | PPA2 |
| CARS | RPL12 | CYCS |
| LAMTOR5 | CTPS2 | AP1S1 |
| FLRT3 | NAA10 | VPS45 |
| PANK4 | SLC12A8 | VPS33A |
| EDIL3 | MAP2K1 | PLCB3 |
| LRRFIP2 | SMU1 | CORO1C |
| GAS6 | SNRPA1 | CCNY |
| OSBP | RAD21 | CREG1 |
| UBE4A | DDB1 | CLSTN1 |
| TARBP1 | DDB2 | KIF5B |
| GLA | PSMB3 | LASP1 |
| GLB1 | DDOST | TUBA1A |
| MDH2 | RPN1 | EIF4A3 |
| MYLK | ASNA1 | IFI16 |
| SMARCA4 | TP53 | LRP1 |
| ZNF598 | NAMPT | SRSF3 |
| IMPAD1 | CSNK2A1 | ARCN1 |
| RPL15P17 | FEN1 | CMAS |
| CERS1 | CHMP1B | GLUL |
| HSPD1P4 | PCSK9 | PPIA |
| BRI3 | WDR77 | HNRNPA1 |
| RHOC | ARPC3 | SART3 |
| EIF2S2 | CPSF6 | RAB31 |
| MOCS3 | GNB1 | HYOU1 |
| CIAO1 | NAA25 | PTPRF |
| RPL29P26 | ATG5 | CD151 |
| ASCC1 | RPL18 | PAICS |
| PMPCA | NUMA1 | SPR |
| STAT1 | SMC1A | DCTN2 |
| STRN | CTSA | RAD21 |
| PDXDC1 | ATP6V1E1 | PSMB9 |
| UBA52 | ATP5MC1 | EPRS |
| STC1 | ATP6V1A | STIP1 |
| TRPV2 | TUBA1A | SEC23A |
| MYL12A | RPS21 | VPS4B |
| KIAA1462 | CDK7 | SMARCD2 |
| PGLS | MCM2 | LAMP1 |
| FBLIM1 | ANP32A | ACSL4 |
| GDF15 | SF3B3 | COL18A1 |
| CCDC80 | GBF1 | PRPSAP1 |
| FMOD | MACROD2 | EXOC1 |
| HRSP12 | CFI | ACP1 |
| GOLT1B | ACTR2 | GYG1 |
| FNDC3B | PRKDC | PITPNB |
| CHMP4A | MDM2 | RPL9 |
| PRRC1 | RPS3A | ITGA6 |
| LIG4 | RPS19 | BLMH |
| PKN1 | RPS23 | RSU1 |
| FEN1 | LRRC47 | SUB1 |
| HSPB6 | RPS3 | CTSB |
| DEFA1 | TOP2A | LSS |
| COG4 | EIF3C | SH3GL1 |
| FAM126A | SIRT2 | APOE |
| SHMT2 | RUVBL2 | PRKACA |
| TM9SF2 | CYFIP2 | BMP1 |
| PLSCR3 | DDX39B | NAPG |
| WDR49 | H2B | GSTP1 |
| DLST | VPS35 | KRT15 |
| S100A13 | PPA1 | ATIC |
| ATP6V0A1 | PARK7 | CDC27 |
| MRPS34 | SLC3A2 | SLIT2 |
| NEGR1 | RSU1 | COPS5 |
| MYO1E | POLR1C | UTRN |
| HSPA12B | MTPN | SRPX2 |
| ATG5 | PSME2 | MYO1B |
| PSMA2 | MED1 | CUL3 |
| USP19 | H2AC | WDR1 |
| GNB4 | RPS10 | PFKP |
| NFKB2 | VCL | MYO1C |
| GZMA | MAT2A | YWHAQ |
| GSTZ1 | EIF2B1 | PODN |
| KLC1 | VPS4B | SCYL1 |
| RPL15P7 | GABRG2 | RPS15 |
| FAM84B | CCT8 | PAMR1 |
| SRC | SRP54 | HSPD1 |
| C16orf62 | RICTOR | GOT1 |
| IGF2BP2 | SLC6A18 | MACF1 |
| HSPB8 | THBS2 | DHX30 |
| ITGB3 | RFC1 | PRPF19 |
| AAAS | CDC42GAP | GLG1 |
| ADCY6 | NSF | PEPD |
| MBD2 | TXLNA | NCBP1 |
| KLF3-AS1 | NF45 | ADAMTS12 |
| RAB23 | PSMC1 | DYNC1I2 |
| COMP | F2RL1 | PSMD7 |
| HSPA13 | ANXA6 | TMPO |
| FUBP1 | MMP28 | OFD1 |
| SCPEP1 | SATB1 | PSME4 |
| TFRC | HBB | EXT2 |
| S100A16 | #N/A | CFB |
| ANGPTL2 | KBF1 | CFI |
| CCT6A | NOTCH2 | HEXA |
| RPL21 | CTNND1 | CAMK1 |
| PCYOX1L | RALB | EVA1B |
| IDUA | SCG2 | KHDRBS1 |
| CLIC6 | TPM1 | RAB5A |
| SLC4A7 | LOXL4 | FARSB |
| SLC7A11 | TGFB2 | PRSS23 |
| CLIC1 | DLG2 | FERMT2 |
| CADM1 | ACTN1 | CTNND1 |
| XRN2 | ANXA4 | MEMO1 |
| STK38L | HEXA | ACLY |
| EXOC5 | C4A | NDRG3 |
| M6PR | PAX5 | HCFC1 |
| RASA3 | COL6A2 | CSPG4 |
| TUBGCP4 | CTSB | C1R |
| AKAP12 | EEF1A1 | EIF4G1 |
| KIF1B | ESD | HSPA9 |
| TIPRL | TNK2 | EIF4E |
| PDIA4 | FBXW7 | ACO1 |
| PPP1R7 | GRB2 | EIF3B |
| KIAA2013 | GNAS | S100A10 |
| HSD17B12 | MARCKS | SNX5 |
| PTGR1 | MYLIP | HNRNPF |
| IGLV1-44 | PARP1 | ADH5 |
| HSPA5 | VDAC1 | POLD2 |
| HNRNPK | SNRPB | SPTBN1 |
| FOCAD | PDHB | KRT5 |
| RPL29 | GNB1 | LOXL2 |
| ZSWIM4 | RECQL | RPL30 |
| ARMCX3 | TGFBR1 | NUCB1 |
| HIST2H4A | RPL26 | ACTN1 |
| HSPA4L | CD90 | LIMK1 |
| DPP9 | TNF | STXBP3 |
| SLC25A6 | VIM | RAB5C |
| P4HA1 | PWP1 | KRAS |
| GSTM4 | RAF1 | GDI1 |
| CELF1 | TRAP1 | RPS10 |
| UBA1 | PTPN13 | TTC37 |
| COL8A1 | DAG1 | APEX1 |
| CUL2 | CFB | CKAP4 |
| MAP1A | RPS5 | LRRC40 |
| RUFY1 | #N/A | ENO1 |
| TRIM46 | DNASE1L1 | UBR4 |
| RPSAP18 | ATP50 | TJP1 |
| CTU1 | CD44 | PSMA7 |
| CSNK1G3 | LOX | COL6A1 |
| COPE | RAC1 | COL1A2 |
| TMOD3 | DAG1 | SNRNP200 |
| PABPC4 | JAG1 | CSRP2 |
| TTR | SCP2 | MYL6B |
| NCDN | EIF3B | RPS4X |
| AK5 | XB | IMPDH2 |
| DNAJA1 | PDIA5 | DYNC1LI2 |
| CSF2 | ADKL | MMP1 |
| IDI1 | HNRNPA0 | DDX1 |
| CCDC25 | FRZB | HSPA8 |
| PIK3C2A | HADH | GOT2 |
| MEST | PLG | GSTM3 |
| PIP | C1S | PROSC |
| CAMK2D | DLST | PPP6C |
| GTF2H4 | CCN1 | CYB5R3 |
| ATP9A | IFL2 | SLC44A1 |
| IFNG | EXT1 | MYO18A |
| MFGE8 | TBCB | RPS2 |
| DPM1 | C1S | GLO1 |
| HS3ST3A1 | HMGB1 | TALDO1 |
| TTC7B | DUT | COPB2 |
| IGF2R | PARG1 | RTN1 |
| SAAL1 | DHX9 | IARS2 |
| QPCTL | SFPQ | LIG1 |
| COTL1 | PPIA | LOX |
| PLOD3 | COL1A2 | KRT8 |
| NCKIPSD | COL1A1 | SPARC |
| TRAPPC11 | SNX20 | DNAJA2 |
| HSPG2 | DRP1 | VCL |
| TOMM34 | SEMA3F | CAND1 |
| GSTK1 | NRP2 | SEC24C |
| IFRD1 | LIMS1 | DPYSL3 |
| TBCB | SNX2 | DCTN1 |
| ATRN | NYCO7 | NEDD4 |
| SPPL2B | HSPH1 | ROCK1 |
| HSPA4 | EIF4E2 | HMGCS1 |
| KCNJ2 | KPNB1 | HPCAL1 |
| COL1A1 | SFRS10 | NPM1 |
| GOSR2 | PPAT | EEA1 |
| FYN | BUB3 | COL3A1 |
| MAP1LC3A | H2AFY | RAB35 |
| PIN1 | FLNA | C3 |
| C8A | RAB3GAP2 | FKBP10 |
| EMG1 | ADAMTS | EIF4A1 |
| LUC7L3 | SPSB1 | UBE2D2 |
| PXDN | ATP6V1G1 | TIA1 |
| MTA1 | TRIP1 | RAI14 |
| THBS2 | PAPPA | NXF1 |
| MMP14 | EIF5 | EEF1E1 |
| SERPING1 | PDIA3 | WDR77 |
| RBPJ | NRG1 | MARS |
| SLC16A3 | IKBKE | CS |
| IGHG2 | DCTN1 | RPL13 |
| TRAPPC12 | PRNP | CYFIP1 |
| GNPDA1 | EIF3G | MCM4 |
| MSN | AP1B1 | IQGAP1 |
| ADI1 | SQSTM1 | PGRMC1 |
| PCNA | NFS1 | GAA |
| SPATA5L1 | EIF2S2 | TNC |
| ANKRD30A | DHX9 | PEF1 |
| KRT8P9 | NXF1 | SPAG9 |
| PKP2 | EHMT2 | PLS3 |
| COMMD8 | THRAP3 | ANKRD13A |
| BIRC6 | LUC7L | SCRIB |
| COL5A2 | MRC2 | KRT14 |
| ACSL3 | PGCP | CYB5B |
| DCAF16 | HEBP1 | PDXK |
| SCARB2 | HSPCO24 | RAB27B |
| MTX1 | HSPC027 | HNRNPDL |
| ARID1A | COPB1 | NAA25 |
| TOR1AIP1 | DYNC2LI1 | GNS |
| WNK1 | G3BP2 | MRC2 |
| IL5 | GYG1 | FTSJ1 |
| KPNA4 | RAB21 | AKR1B1 |
| EIF2AK2 | PDGF | BCAM |
| EPM2AIP1 | RAB14 | SNRPA1 |
| HNRNPA3 | KIAA0291 | PSMD6 |
| DNMBP | COP9 | GDF11 |
| OSBPL9 | MG50 | NNMT |
| IGH | CST4 | HMOX2 |
| ITGA3 | MATR3 | SNRPG |
| PEA15 | PFDN2 | TXN |
| CNRIP1 | SEP09 | RICTOR |
| UGP2 | COMMD10 | NAA15 |
| FLRT2 | ARL6IP1 | SNX6 |
| NAP1L1 | EHD2 | LRRC59 |
| HERC3 | EHD3 | GAPDH |
| WDR36 | PHB2 | ZYX |
| ABCF2 | CBX1 | PLOD2 |
| BRD3 | HDCKB03P | PLG |
| KPNB1 | UGGT1 | DNM3 |
| CAPN2 | HNRNPH1 | RPS6KB1 |
| B4GAT1 | ATRN | TAGLN |
| UBE2Q1 | #N/A | SNRPB |
| ZSWIM8 | #N/A | GNB2 |
| RPS16P1 | DHRS1 | SRSF2 |
| GSTA5 | ZNF43 | CAV1 |
| CDC42BPA | CBFB | EEF1B2 |
| UBE3A | MYG1 | AP2A1 |
| DDX46 | PFN2 | PSMC4 |
| SYNCRIP | FKSG13 | PYGB |
| NUCB2 | EGFR | DDB1 |
| SPACA1 | PPA2 | CFL1 |
| GBF1 | #N/A | CDK7 |
| PPP2CA | NANS | KRT2 |
| LARP4B | ARPC5L | ANP32B |
| VPS41 | TGFBI | BLVRB |
| BLOC1S5 | HMGCS1 | SBDS |
| STARD10 | COPB2 | MAP1S |
| WRAP73 | PSMD7 | PTPN11 |
| FUCA2 | JUP | GBE1 |
| TNFRSF10B | BASP1 | RALB |
| SORBS3 | CDCA8 | VIM |
| SACM1L | DCI | VTN |
| RPL3 | NUDCD1 | WDR61 |
| PIK3R4 | CSRP2 | ANPEP |
| RASA4CP | RPLP0 | CD82 |
| RABL6 | TMEM109 | ITGB1 |
| RAB13 | SEC22B | CHP1 |
| IL17B | HNRNPR | DCTN4 |
| HEATR5B | XPNPEP3 | CLTA |
| OSGEP | APEX1 | SNX1 |
| FAP | EFTUD2 | ALDOA |
| DUS3L | TSG101 | CARM1 |
| RPS18 | SNRPF | CMPK1 |
| ILF3 | TUBB6 | COMMD3 |
| GPRC5A | KRT14 | AP2A2 |
| ZC3HAV1 | LGMN | CD63 |
| SARS | RAP2C | RAB32 |
| NARS | PRDX4 | ACTR1A |
| CLTC | BLMH | COL12A1 |
| DDX41 | HNRNPF | PARP1 |
| ITIH3 | PSMG3 | SEMA5A |
| TCP11L1 | VTN | PRDX1 |
| TBC1D8B | ATP5F1B | LRPPRC |
| SBF1 | KPNA2 | DOCK10 |
| CCL7 | GINS4 | APP |
| SLC9A1 | UNC-45 | TRIOBP |
| SNX4 | ILKAP | ANXA4 |
| SCAF4 | FSCN1 | SERPINE1 |
| NUTF2 | LASP1 | TGFBI |
| DCUN1D3 | GSS | CUL5 |
| SNX9 | IFITM3 | HNRNPA0 |
| RPL18A | PRDX3 | GRHPR |
| MRPL39 | PCMT1 | ICAM1 |
| ARHGEF40 | CAPN1 | TSKU |
| HGH1 | TXNDC12 | MMP2 |
| RPP30 | CYCS | VPS25 |
| TPI1 | IGFBP6 | MCM7 |
| KLHL9 | MYC | ALDH9A1 |
| LTF | SRP19 | AP3S1 |
| MAP7D1 | ATP6V1C1 | IGSF8 |
| UPF1 | CDC27 | FBN2 |
| OSBPL3 | ATP1B3 | ANXA1 |
| LRG1 | EIF2AK2 | TPT1 |
| SNAP47 | CAPNS1 | RAB11B |
| VPS51 | CD9 | ECE1 |
| SMARCA1 | GOLPH3 | CPSF1 |
| GPX8 | RTCD1 | SMARCC1 |
| SUSD6 | SERPINB2 | PDHB |
| DIP2A | KRAS | HNRNPR |
| KLHL21 | COL8A1 | ADAM10 |
| EGFR | JTV1 | RANBP2 |
| CCDC129 | KRT6A | CBR1 |
| IL1RAPL1 | AP2A1 | NAA10 |
| RANBP9 | VAT1 | CALR |
| HMGA1 | SF3A2 | EIF3C |
| PRKAG1 | DNAJA2 | SSR4 |
| SH3GLB2 | LRRC59 | HSPH1 |
| GNAQ | CPOX | GREM1 |
| PRPF31 | RAB5A | PGAM5 |
| GDF6 | GNBP1 | LDHB |
| TAOK1 | FBLN1 | CASK |
| POTEI | TUBB2C | CALU |
| TTLL12 | NIT2 | RECQL |
| SCYL2 | SRPX | ATP1B3 |
| CAPN1 | SRPX2 | NF2 |
| UNC13A | FBNL1 | NOTCH2 |
| KNTC1 | PABPC1 | COL4A2 |
| PES1 | CHPF | RPS3 |
| VASP | RRP9 | SAP18 |
| PGGT1B | GRPEL1 | ATP6V1A |
| SCGB2A1 | KRT5 | DIS3 |
| ADSL | GNAI3 | EHD4 |
| CD109 | CNN3 | PPP2R4 |
| LGALS8 | DYNC1LI2 | GCLM |
| PPP2R5E | VASP | STX2 |
| COPS4 | VPS45 | GNAS |
| MCM10 | OLA1 | RDX |
| ABHD14B | PRPS2 | PCOLCE |
| CP | PODN | RNMT |
| OGDH | PLBD2 | GART |
| SRSF4 | HSPA9 | ARPC1B |
| SRSF1 | EDIL3 | TNFAIP2 |
| RPSAP61 | C9ORF23 | PRIM2 |
| KRT74 | AXL | HK1 |
| SRP54 | OSGEP | CEMIP |
| JAK1 | MCM8 | MAT2B |
| NAA30 | EIF5B | PACS1 |
| RAP1B | MAN2B2 | PDE12 |
| RPS24 | VAT1L | RPL12 |
| DCTD | RPRD1B | PGK1 |
| MRGPRF | PRP4 | CUL4B |
| GBA | ARSK | SH3BGRL3 |
| RAB4B | CUL4B | LAMA1 |
| DOCK4 | HSPA13 | MOXD1 |
| GLIPR2 | CDH2 | BCAS3 |
| GNPNAT1 | KDELR2 | MON2 |
| IKBKE | PLOD2 | PAPPA |
| TXNL1 | CNNM3 | BZW1 |
| CPNE2 | C3ORF58 | PSAP |
| ACTA2 | AP3B1 | SNTB2 |
| LRPAP1 | COPA | H2AFV |
| DDX59 | CHD4 | RRP9 |
| SRM | STIP1 | XRCC1 |
| VRK1 | RAB5B | DBNL |
| HECTD3 | WASF2 | AP1B1 |
| VPS4A | COL1A2 | NRP2 |
| PPP6R2 | IPO5 | AQR |
| CUTA | PRKAA1 | G3BP2 |
| CYR61 | SLC9A391 | ARPC1A |
| UBA2 | MMP19 | EPS15L1 |
| AFP | SMARCC1 | TYMS |
| AP2M1 | NUP62CL | PHB |
| IFT27 | SFPQ | PPP2R2A |
| TUBB | FUCA2 | CDK13 |
| ST8SIA4 | UBE2I | ESD |
| HSPA1L | YWHAZ | HSP90AA1 |
| RPA3 | EHD4 | PSMD14 |
| ARPC5 | TMPO | IPO5 |
| APMAP | RPL22 | CAP2 |
| MTMR9 | SNX5 | ABCF1 |
| MAP4K4 | FSN | MYOF |
| UXS1 | CPXM1 | VPS13A |
| TMEM2 | RAB22A | DTYMK |
| PRKRA | KIAA1967 | YKT6 |
| QPCT | COL6A2 | POLR1C |
| CCT2 | CAD | EEF2 |
| DDX39B | MAT2B | ANO6 |
| DFNA5 | MARCKSL1 | MARCKSL1 |
| POLR3A | ANXA2 | ACTN4 |
| ECHDC1 | SLC25A5 | DDX3X |
| PMPCB | PDCD61P | PDLIM7 |
| TRMT11 | VASN | XRCC6 |
| SDHA | CCNY | EIF2S1 |
| HIST1H2BB | SERPINH1 | RALY |
| IGHA2 | EEF1A1 | EEF1D |
| TUBG1 | BICD2 | NAGLU |
| COPA | B4GALT5 | HNRNPC |
| TCF25 | DPYSL3 | TPP2 |
| WWP2 | CACYBP | MAP1B |
| FGA | EIF3D | LDLR |
| UPF2 | CPZ | DCXR |
| ST6GALNAC6 | PTPN23 | TUBA1C |
| URGCP | RPL13 | RPS15A |
| TOM1L2 | RAB13 | SYPL1 |
| USP7 | RAP1GDS1 | WBP2 |
| XPO4 | #N/A | LRRC17 |
| SWAP70 | RPL9 | RPS19 |
| MYO1D | GRIPAP1 | ATP6V1E1 |
| WDR13 | LMNB1 | HSPBP1 |
| NOP2 | FTH1 | PPT1 |
| STAT2 | PSMC3 | AHCY |
| HIST1H2BL | CSTF3 | NID2 |
| FHL3 | NT5DC1 | TPP1 |
| RPN1 | MAPRE1 | ARPC3 |
| STOM | HIP1 | VCP |
| MYO9B | PPP1R12A | PWP1 |
| SLC3A2 | ARHGAP18 | RPS9 |
| IPO7 | HSPA5 | VAT1 |
| OLA1 | ROCK1 | COL6A3 |
| SLC44A2 | TBR1 | FSCN1 |
| PRKCD | GEMIN5 | HNRNPA2B1 |
| CLN3 | EIF3A | VBP1 |
| PSME2 | FRMPD1 | SERPINH1 |
| AP1M1 | GLT8D1 | HEBP1 |
| KRT6B | HSPA4 | IFITM3 |
| RRM1 | L1CAM | EIF5B |
| MCM2 | KIF5B | RAB34 |
| PATL1 | UBA6 | SNX12 |
| KRT1 | AQR | CRAT |
| POR | ANO6 | EFHD2 |
| PFN1 | NEDD4 | PSMD3 |
| GDF5 | BMP1 | TUBB6 |
| HDAC1 | ZFR | KRT18 |
| ST3GAL1 | RCD1 | CD44 |
| FAM71F1 | IQGAP1 | LARP1 |
| SULT1E1 | AIFM1 | TTC9C |
| PSMG1 | USP11 | RPA1 |
| TMED4 | EIF4G1 | TPD52L2 |
| TKFC | CFH | RPL24 |
| COG3 | SMARCD2 | PPP5C |
| MIF | SMARCA1 | SMC1A |
| PAFAH1B3 | SMC2 | S100A11 |
| RPSAP8 | ITGA3 | UBE2Z |
| FAM91A1 | PFAS | AXL |
| FNTB | ALDH9A1 | PHB2 |
| IGBP1 | ERI1 | CTPS1 |
| SCAMP1 | CD151 | CDK2 |
| LMNB1 | MTMR12 | EHD1 |
| HGS | HOOK3 | PSMB2 |
| RPS2P12 | FBLN1 | PICALM |
| TNFSF18 | HNRNPQ3 | SCP2 |
| SORT1 | VPS13A | SH3GLB1 |
| MAP3K3 | HMCN1 | HOOK3 |
| FAT1 | HSRBC | TRAP1 |
| GAPVD1 | B3GALT6 | SHMT1 |
| FJX1 | COL3A1 | DST |
| DIAPH1 | TBC1D15 | CHMP1B |
| POU2F1 | LAMA5 | PRPF8 |
| TGFB2 | RAB27B | HNRNPUL1 |
| EPS8L2 | RAB32 | PSMA4 |
| CPAMD8 | URB | POLD1 |
| EPB41L2 | HSPA8 | CPSF7 |
| TOLLIP | GPR19 | PTPRK |
| IPO4 | SEPTIN6 | RPS23 |
| CNOT10 | CMAS | DNASE1L1 |
| SFXN3 | MMP9 | TSN |
| EPB41L3 | FBLN7 | PAPSS1 |
| DHX29 | CD109 | FTH1 |
| SFPQ | TUBB2A | CHID1 |
| ATP5B | TUBB4B | KHSRP |
| ENPP1 | SEPT8 | EHD2 |
| TTC28 | PHB | RPN2 |
| DPP8 | HZGJ | RAB12 |
| MPZL1 | NCSG135 | GNAI3 |
| STRAP | CPNE3 | PPP1CC |
| PPFIBP1 | GMFB | YWHAB |
| LYAR | RRAS | VEGFC |
| TK1 | NAP1L2 | OGT |
| LTBP1 | RECQ1 | FN3KRP |
| PCK2 | CAP1 | GNAI2 |
| TRAPPC6B | SMARCE1 | NUDCD1 |
| SPP2 | SNX12 | XPO7 |
| TERF1 | EEF1D | BSG |
| STK17B | IGF2R | SLC1A5 |
| PGAM1 | NUCB1 | ACAA2 |
| RAVER1 | GNG12 | LAMA2 |
| PRMT5 | CS | G6PD |
| MRPL21 | MYL9 | UBB |
| ATP6V1H | ICAM1 | RPS3A |
| SLC35B2 | CSPG4 | RPL22 |
| TMEM106B | P4HB | STAT3 |
| RAB3B | OLFML1 | AGRN |
| TMEM200A | TXNDC4 | RAB22A |
| RCC2 | SELE | RAB6A |
| KRT3 | HRNR | KRT10 |
| CDC42 | FSN | GLUD1 |
| LRRC57 | OCIAD1 | NQO1 |
| TNKS1BP1 | LAMP2 | AKR7A2 |
| HSPA7 | CPIN1 | CARHSP1 |
| ACAT2 | GNAI2 | NSF |
| SEC22B | CLDN11 | HBB |
| TNFRSF10A | RAB11B | PRNP |
| IGLC3 | MPZL1 | PRDX2 |
| MED4 | SCGF | CNN3 |
| ATP5C1 | RABA2 | RPS20 |
| RPSAP12 | NPM1 | ANXA6 |
| UBE2I | HNRNPK | ROCK2 |
| PLCD1 | CEP55 | RCC1 |
| KCTD12 | ACT | TSPAN9 |
| APEH | #N/A | FLNB |
| PLP2 | POLD1 | PFDN2 |
| MATR3 | TRIOBP | RAN |
| EXT1 | FHL2 | COPB1 |
| GLRX | PLCG1 | PTPN23 |
| MTCH2 | HSP90AA1 | PLEC |
| ARMCX2 | HSPB1 | DHX36 |
| FAH | SAFB2 | UBE2K |
| RAPGEF2 | RNASET2 | RPL35A |
| GMPR | ESYT2 | DKC1 |
| TUBB3 | TDP43 | RAE1 |
| SYDE1 | DCD | UBA6 |
| MYBBP1A | ARCN1 | ADD3 |
| PLAA | VIM | SMAD3 |
| COPS7B | EDDM3A | EIF3D |
| COL2A1 | EDDM3B | RCN1 |
| BASP1 | WFDC8 | SMU1 |
| MPI | CD163 | AEBP1 |
| UFM1 | WFDC2 | LAMP2 |
| ITCH | ELSPBP1 | PA2G4 |
| VDAC1 | ELSPBP1 | RPL38 |
| F8A1 | ELSPBP1 | CTPS2 |
| IQGAP2 | ELSPBP1 | IDH1 |
| FAM120B | ELSPBP1 | HIST1H4E |
| GSPT2 | ELSPBP1 | PPME1 |
| KRT17 | ELSPBP1 | PGM2L1 |
| GPD1 | OPA1 | SPTAN1 |
| EHBP1L1 | LRPPRC | ATP6V1C1 |
| NME2 | NF110B | PACSIN2 |
| GTF2I | PDIA4 | PSMA1 |
| MOGS | WFDC11 | PARK7 |
| SERINC5 | WFDC6 | JUP |
| GDAP2 | WFDC3 | BCAT1 |
| GLMN | FAM22A | ALDH2 |
| KIF1BP | KRT1 | DDOST |
| CLNS1A | CD63 | SEC31A |
| INTS9 | GFT2I | VAT1L |
| PSMC1 | GNS | HAT1 |
| VASN | PROS1 | SLC25A5 |
| RPL12P19 | HGS | NOMO2 |
| ITIH2 | FTSJ1 | CCT5 |
| C4A | HLAC | AP2S1 |
| AK1 | C1S | PRMT1 |
| PRDX4 | HLAC | PKM |
| ARPC5L | APP | NIT1 |
| NLE1 | TLN2 | NOP58 |
| DHRS1 | PRMT1 | VDAC3 |
| RPL19 | HSPA1A | WNT5B |
| CXCL2 | PSMB9 | RUVBL2 |
| RANBP6 | DNAJB1 | SRI |
| EIF3I | CTNNA3 | TCP1 |
| MPG | PREP | CDC34 |
| CYFIP2 | KIAA0051 | HRNR |
| LYPLA1 | S100A11 | SRP14 |
| PAN3 | RCN1 | HIP1 |
| ARHGAP18 | CDC47 | CALD1 |
| NCEH1 | PAFAH1B3 | CHST14 |
| PLCG1 | RPS13 | AHNAK |
| EIF2B2 | MYCBP | PDIA3 |
| ZAK | SMAD3 | MTMR12 |
| CDK17 | PCOLCE | ANXA11 |
| PLXND1 | GCP170 | DPYSL2 |
| ATP6AP1 | DDX46 | CORO1B |
| ACSS2 | KIAA0640 | MYH9 |
| P4HA2 | KIAA0726 | RAB5B |
| NAA50 | KIAA0801 | COPG2 |
| IGFBP4 | NARS1 | FN1 |
| MPP1 | CNP | SF3B1 |
| CCZ1 | ACTR3B | CDK5 |
| WDFY3 | KIAA0862 | SRPX |
| DNM1L | KIAA0908 | SMARCE1 |
| PSPC1 | ASY | QSOX1 |
| BRMS1 | DNAJC9 | MGAT5 |
| CHPF2 | KIAA1398 | UAP1 |
| TGFB1 | KIAA1470 | BTF3 |
| F13A1 | KIAA1499 | HMGB1 |
| HIST1H4B | RPN2 | APRT |
| RBPMS | #N/A | HNRNPH1 |
| PCMT1 | COPG2 | TBC1D15 |
| PUM1 | RAC1 | LPP |
| CTSZ | CCBE1 | RP2 |
| RPL5 | ABCF1 | RAB15 |
| FGF19 | #N/A | TUBB4B |
| DDAH1 | DHX8 | PHGDH |
| ISYNA1 | #N/A | PSMA3 |
| ITGAV | MRT1A | PPP3CA |
| PBLD | HMFT1638 | CPNE1 |
| OPLAH | XRCC1 | FAT4 |
| ZNF638 | GART | IDE |
| FLNA | PCBP2 | DDX17 |
| EPHB2 | CCT3 | CD47 |
| OSM | RPL5 | PCYT2 |
| CLDN11 | DDX3X | FLNC |
| ADAM23 | RFC5 | NCL |
| GAK | ACAT2 | NEO1 |
| RRAS | TRIM25 | RAP2C |
| EPHA2 | H2AFV | PLD3 |
| CEPT1 | COL5A2 | PREP |
| SMC2 | ARPC1A | AGL |
| LIPA | PYGB | RPLP1 |
| EHD3 | TIA1 | GMPS |
| DDX49 | RHOQ | ITGA2 |
| MOV10 | NCBP1 | H2AFY |
| PFN2 | SUB1 | RANGAP1 |
| TRIM28 | MAP3K7IP1 | PSMD2 |
| HERC5 | LDLR | MAP2K1 |
| FKBP15 | EBNA2 | AP2B1 |
| HNRNPL | LIMA1 | FBLN1 |
| CSTF1 | HNRNPC | VPS35 |
| UBE3B | TOLLIP | HNRNPD |
| GIGYF2 | STAU1 | CCT3 |
| DHCR7 | DDX17 | RBM3 |
| SZT2 | COLA1 | POLR2B |
| PSMC5 | SPOCK2 | COMMD10 |
| DAG1 | GBE1 | CAP1 |
| HLA-A | RPA1 | FLII |
| FAM64A | SLC1A5 | SEC24B |
| ACOT8 | AEBP1 | DNAJB1 |
| GSTM5 | WDR1 | RAB1A |
| MAPK3 | SPTBN1 | ARL1 |
| NCKAP1 | PSAP | CSNK2B |
| RPLP0 | PRPF4 | NUDC |
| CDK4 | COP9 | RAB21 |
| TRIM3 | GLO1 | GSN |
| FLAD1 | AP1M1 | EIF2B1 |
| HBS1L | PPP2R5E | DNM2 |
| MVK | GSTM3 | TSG101 |
| CHI3L1 | SERPING1 | CSNK1D |
| GLDC | PSMD2 | TUBB2A |
| S100A11P1 | CAPN2 | SERPINB6 |
| GPC5 | SPTAN1 | EIF2S3 |
| CDH13 | PLOD2 | EFTUD2 |
| THOC6 | ITGAV | P4HB |
| RHEB | THBS1 | ANKFY1 |
| SEC16A | ANPEP | ATP5A1 |
| SNAPIN | MAN2B1 | PDIA5 |
| ZW10 | Hsp40 | SNRPD1 |
| SH3BP4 | CD146 | PTBP1 |
| BDH2 | CAV1 | RPL26 |
| CASP8 | LRP1 | PPIB |
| RHOG | PGRN | PSMD10 |
| TRIM40 | MATRIX | PABPC1 |
| ACO2 | EIF3M | PSMC6 |
| TMX3 | VIP36 | NAGK |
| BAG2 | TXNDC9 | GLCE |
| BIN1 | NUP54 | DDX5 |
| NUDT4 | MARS | TRIM25 |
| SERPINE2 | PLS3 | ACAT1 |
| SLC25A4 | GLUD1 | NELFB |
| MAP4K3 | TEMD10 | HNRNPM |
| POLR2C | RAB18 | CANX |
| ACP2 | LIN7C | LAMA5 |
| NAPA | TSPAN9 | ACTR2 |
| SEC24A | PSMD11 | HSPB1 |
| ANG | VCAM1 | THRAP3 |
| TXNRD1 | GOT2 | SF3B3 |
| PHKG2 | BADH | TNFRSF11B |
| CD46 | NSFL1 | CFAP20 |
| C1S | HNRNPAB | PARVA |
| BLVRA | NADPH | PSMG3 |
| PRR36 | CKAP4 | EIF5A |
| SLIRP | COMM | ENAH |
| CDKN2A | YARS1 | TBCD |
| RAP1GDS1 | MYO1C | GEMIN5 |
| NPLOC4 | MYO1B | GRB2 |
| RAB3GAP1 | GOLGB1 | AP3B1 |
| ABCF3 | U2AF1 | CSNK2A1 |
| CCDC64B | HNRNPK | DYNC1H1 |
| FBN1 | BETA CATENINE | RAB7A |
| DECR1 | EIF2S2 | G3BP1 |
| CNP | DDX5 | KPNA6 |
| IL1RAP | SQSTM1-ALK | PPP1R12A |
| STXBP1 | TPKR | UROD |
| PFAS | TBCD | PFKM |
| MTA2 | GAPDH | MAN2B1 |
| CNOT1 | COL3A1 | EEF1A1 |
| RPL7A | TGFB1 | THBS1 |
| GOLGA7 | NDPK1 | MTAP |
| SNX3 | TCP1 | CTSD |
| SKIV2L2 | LAMA1 | TOR1B |
| MICAL1 | VARS1 | RPL18 |
| ACTR1B | EEF1B2 | SEC24D |
| RGL2 | FBN1 | CTNNA1 |
| UROS | DDX1 | DNAJC13 |
| RPSA | MCM4 | CST4 |
| GSR | TNC | CAPZA1 |
| PPAT | 90K | HINT1 |
| ARF4 | ACTR1A | CSTF3 |
| STARD13 | KRP | RNH1 |
| ESYT1 | EMP3 | GNG12 |
| PAPPA-AS1 | SLC25A1 | LMNA |
| PREB | SEC23 | ASL |
| QSOX2 | PRSS11 | CAD |
| COX6C | NF2 | NANS |
| RAB10 | FERMT2 | CTSA |
| MAPRE1 | KRT9 | RPS13 |
| SLC25A3 | TKA1 | NUP43 |
| UXT | HLAC | LRRC47 |
| TUFM | DKC1 | COL7A1 |
| NUCKS1 | CAF1 | PSMD11 |
| CANT1 | H/ACA | PPA1 |
| ACTR10 | NOGO-A | DCD |
| ITIH1 | SRPK1a | SAR1A |
| LMAN2L | OGT | CAPNS1 |
| KIF16B | DHX36 | ECHS1 |
| SCFD1 | ATP1B1 | DHX15 |
| CLU | RAB9A | PRPF4 |
| OLFML3 | MYH9 | NAMPT |
| ITGA5 | IGFBP-5 | CALM1 |
| SLC20A1 | PPIB | ERP29 |
| NCAPG | DNAJB1 | SLC16A1 |
| CHPF | VEGFC | RFC5 |
| ATP6V1D | PPIH | CHD4 |
| CAPN5 | PDGFRL | RAB3GAP2 |
| CLIC4 | Rab6A | LRIG1 |
| NAE1 | LAMA2 | RPS5 |
| CDH2 | HLAC | KRT9 |
| MARCKS | CORO1C | PAFAH1B2 |
| CNDP2 | ACYL-COA | TAOK3 |
| ITPA | MTAP | FHL2 |
| POGLUT1 | CR1 | ESYT2 |
| PAPSS2 | NACALM | EIF5 |
| RBM43 | MCM7 | VPS29 |
| KRT17P3 | ARPC1B | RAB18 |
| RASA1 | MLL4 | VAMP3 |
| TRAPPC9 | SNX9 | USP11 |
| PPA2 | MOXD1 | HSPA1A |
| ATL3 | ENPP1 | TNPO1 |
| CTBP2 | ECHDC1 | CCT8 |
| CYCS | EVA1A | PRDX3 |
| Sep-07 | KIAA0776 |  |
| PPM1F | anti-ALDH7A1 |  |
| RPS14 | AP3S1 |  |
| AP1S1 | LINJ |  |
| NMT2 | pcdA |  |
| LRRC41 | RPS15A |  |
| VPS45 | CRA_a |  |
| NME4 | RBM3 |  |
| VPS33A | CRA_b |  |
| OSBPL11 | SUPT6H |  |
| TSSC1 | L23a |  |
| PLCB3 | EFHD2 |  |
| ATP11A | ZYX |  |
| CORO1C | LIG1 |  |
| PAFAH1B1 | NIT1 |  |
| HSPA6 | NAP |  |
| TMA7 | IFI16 |  |
| GSK3B | LMNA |  |
| UNC80 | LMNC |  |
| ADNP | S100A10 |  |
| SPTBN4 | TfR1 |  |
| PPP2R5B | GNB2 |  |
| SEMA3C | IQGAP1 |  |
| RPL13A | SEPTIN10 |  |
| PIP5K1A | CDK5 |  |
| CCNY | HK1 |  |
| USO1 | DNAJ |  |
| LSM6 | ITGA2 |  |
| CREG1 | CAP2 |  |
| TFG | CRA_a |  |
| IKBKAP | SFRP-1 |  |
| SCCPDH | HIST1H4E |  |
| CLSTN1 | STXBP3 |  |
| KIF5B | SARS1 |  |
| B4GALT4 | YTHDC1 |  |
| TLN1 | SORT1 |  |
| RSL1D1 |  |  |
| DCHS1 | BLVRB |  |
| ABI3BP | BCAM |  |
| TENM3 | NAPA |  |
| EXOC6B | PTK9 |  |
| TXNIP | C1QL4 |  |
| XRN1 | CLTA |  |
| TCIRG1 | C9ORF19 |  |
| RRAS2 | ACO1 |  |
| GALNT1 | RPL3 |  |
| LASP1 | XRCC6 |  |
| TUBA1A | KRT10 |  |
| PRLHR | KRT17 |  |
| UBE2N | FKBP10 |  |
| SSB | DBNL |  |
| SNX27 | BSG |  |
| EIF4A3 | LARP1 |  |
| CNOT6L | G3BP1 |  |
| DIP2B | PDGFRB |  |
| IFI16 | IK |  |
| MPHOSPH6 | #N/A |  |
| MMP11 | SKP1 |  |
| KRT8P45 | P4HA2 |  |
| RPL23A | FBN2 |  |
| LRP1 | ANXA1 |  |
| RRP12 | TXNL1 |  |
| NPEPPS | PROSC |  |
| SRSF3 | GSR |  |
| ARCN1 | PPP2R2A |  |
| ANAPC13 | LOXL2 |  |
| UROC1 | ASAH1 |  |
| LTBP2 | LRRFIP2 |  |
| DPP3 | COL7A1 |  |
| CMAS | TMEM113 |  |
| EPS8 | PDE12 |  |
| IL11 | PSMD6 |  |
| STOML2 | LRIG1 |  |
| TRPT1 | SNX6 |  |
| DPP4 | SUPT16H |  |
| VPS36 | TXNDC10 |  |
| PARP14 | #N/A |  |
| GLUL | DCN1 |  |
| ARHGEF17 | PDGFRL |  |
| RPSAP58 | CUL5 |  |
| NTRK1 | RDX |  |
| RPS4Y1 | TAGLN |  |
| TPM4 | CD82 |  |
| SHPK | PAMR1 |  |
| PRKAB1 | PSMA1 |  |
| MSH2 | PARVA |  |
| PPIA | CRA_a |  |
| HNRNPA1 | HNRNPM |  |
| SART3 | ELAV1 |  |
| THG1L | KHSRP |  |
| CYTH1 | C19ORF10 |  |
| RAB31 | SH3GL1 |  |
| HYOU1 | LMNB2 |  |
| RPSAP15 | CIRBP |  |
| YRDC | POLRAE |  |
| HIST2H2AA3 | LIMK1 |  |
| ILK | FH |  |
| CCBL2 | HSPD1 |  |
| TUBAP2 | NRP2 |  |
| PTPRF | MYL1 |  |
| SLC7A1 | FN1 |  |
| CD151 | XRCC5 |  |
| ABLIM3 | NCL |  |
| PAICS | COL6A3 |  |
| FAM160B2 | DTYMK |  |
| DPH5 | RBM8A |  |
| RPL10A | CLSTN1 |  |
| SPR | SRM |  |
| ELL2 | POR |  |
| DCTN2 | RPS4X |  |
| PGAM2 | VBP1 |  |
| RAD21 | FAM50A |  |
| FARSA | SSR4 |  |
| FGF16 | AGL |  |
| PIK3R2 | C20ORF27 |  |
| DGCR2 | FBLN1 |  |
| UBR2 | C1ORF198 |  |
| ZNF614 | TTC9C |  |
| PSMB9 | C6ORF132 |  |
| PRKACB | EHD1 |  |
| ACTA1 | PACS1 |  |
| GSTT2B | SERPINH1 |  |
| EPRS | PRSS23 |  |
| HAS1 | TPD52L2 |  |
| TMEM192 | CTSZ |  |
| DLG5 | NCOA5 |  |
| MAP2K2 | CTSA |  |
| ARHGEF10 | TOP1 |  |
| TSR1 | C20ORF77 |  |
| RHBDF1 | DLGAP4 |  |
| STIP1 | ITGB4BP |  |
| ANAPC1 | ITCH |  |
| SPG11 | RALY |  |
| SEC23A | CHMP4B |  |
| RPS3P3 | TFPI2 |  |
| ERLIN1 | ADAM10 |  |
| TLE1 | SNX1 |  |
| TOR1A | SMC4L1 |  |
| TRIM56 | PFN2 |  |
| VPS4B | CRA_a |  |
| SERPINB3 | PDIA5 |  |
| STX5 | RPL24 |  |
| EIF3F | TBC1D23 |  |
| LDLRAP1 | PPP4C |  |
| SMARCD2 | TFIID |  |
| RIC8A | MYO1D |  |
| STX16 | CRA_c |  |
| HERC2 | RTN1 |  |
| SVIL | RB25 |  |
| TMEM263 | NUMB |  |
| PYGL | YY1 |  |
| LAMP1 | DYNC1H1 |  |
| ALOX12P2 | PLEC |  |
| PSMD5 | SCRIB |  |
| MTOR | EEF1D |  |
| TSNAX | SNTB2 |  |
| TBXA2R | FLNC |  |
| NLGN2 | CALD1 |  |
| MSRA | ILF3 |  |
| A1BG | DNM2 |  |
| ACSL4 | DNM2 |  |
| COL18A1 | PRDX2 |  |
| HGSNAT | GLT25D1 |  |
| TECPR1 | COPE |  |
| PRPSAP1 | PDLIM7 |  |
| EXOC1 | THOC6 |  |
| DDAH2 | HAGH |  |
| ACP1 | ITGB1 |  |
| SCAMP3 | PFKP |  |
| GYG1 | COPS5 |  |
| FCHSD1 | HSPA5 |  |
| EPPK1 | AK1 |  |
| CDK9 | TOR1B |  |
| PLOD1 | RPL7A |  |
| SMURF2 | COL1A2 |  |
| PITPNB | DNM1L |  |
| TSC1 | WNT5B |  |
| B3GAT3 | WBP2 |  |
| RPL9 | GAA |  |
| TRAPPC1 | FN3KRP |  |
| PSME1 | #N/A |  |
| RABL2A | TP53 |  |
| RPL28 | 0 |  |
| GGT1 | C1QBP |  |
| NUP93 | CREG1 |  |
| R3HCC1 | TIP41 |  |
| DENND6A | ARPC5 |  |
| HADHB | TPR |  |
| COG6 | IPO9 |  |
| TAS2R60 | #N/A |  |
| SNRPE | EIF3G |  |
| CXCL8 | TNFRSF11B |  |
| LOC400750 | RPL35A |  |
| PTPRG | CHP1 |  |
| ITGA6 | SOD3 |  |
| FAM65A | EPRS |  |
| BLMH | #N/A |  |
| ZBTB4 | CLTC |  |
| ADAM9 | SRSF2 |  |
| ISOC1 | #N/A |  |
| RSU1 | ITGA3 |  |
| PVRL2 | C1ORF33 |  |
| RPL4 | MAOA |  |
| TGM2 | HS6ST1 |  |
| SUB1 | KARS1 |  |
| EDC4 | GLG1 |  |
| NME1 | MAP1B |  |
| INHBB | BTF3 |  |
| GRB7 | DHFR |  |
| CTSB | KRT8 |  |
| ATP5O | KRT18 |  |
| NRG2 | HNRNPA1 |  |
| PTAR1 | ITGA5 |  |
| LSS | #N/A |  |
| ISG15 | CDK2 |  |
| BIN3 | FAM621 |  |
| PFKL | MYL6B |  |
| MCAM | ATP5F1B |  |
| NPTN | 0 |  |
| SH3GL1 | CPSF6 |  |
| IL23A | YEATS4 |  |
| SEC23B | ARL1 |  |
| APOE | ERP29 |  |
| RAD23B | OFD1 |  |
| TM9SF4 | HMGB3 |  |
| ERO1L | NAGK |  |
| WDR26 | RAB1A |  |
| COPG1 | LTBP1 |  |
| PRKACA | MFGE8 |  |
| PDCD5 | RNH1 |  |
| HIST1H4I | CTSD |  |
| RIOK2 | CSNK2B |  |
| PSMB10 | CUTA |  |
| BMP1 | SRSF3 |  |
| DHRS11 | TBCC |  |
| SRSF7 | C6ORF108 |  |
| ELAVL1 | CD2AP |  |
| NAP1L4 | HMGB2 |  |
| CXCL16 | POLR2B |  |
| ARMC8 | SCARB2 |  |
| NAPG | SEPTIN11 |  |
| MGLL | HNRNPD |  |
| PCBP1 | HNRNPDL |  |
| GSTP1 | SEC31A |  |
| KRT15 | ADH5 |  |
| HK2 | PPP3CA |  |
| DDX58 | CXCL9 |  |
| LTA4H | SERPINE1 |  |
| MTR | PABPC4 |  |
| SPOCK1 | KIAA0319 |  |
| KRT16 | RCC1 |  |
| RPL15P3 | DNAJC8 |  |
| SRPR | SH3BGRL3 |  |
| ATIC | COL4A2 |  |
| PPP1CA | LSS |  |
| EDC3 | PYGB |  |
| UBASH3A | SNRPB2 |  |
| PHKA1 | NSFL1 |  |
| WDR6 | HAT1 |  |
| SRRM2 | DYNC1I2 |  |
| CDC27 | PHF6 |  |
| TIMM50 | C3 |  |
| PDCD1LG2 | G6PD |  |
| SLIT2 | KRT2 |  |
| SYNE1 | F10 |  |
| COPS5 | NOMO2 |  |
| RAB11FIP5 | CPM |  |
| TRIM41 | CORO1B |  |
| UTRN | RAB12 |  |
| SRPX2 | ACOT9 |  |
| CDK5RAP3 | CTNND1 |  |
| ALDH1L1 | TXNRD1 |  |
| SLC25A13 | XPO7 |  |
| NISCH | RPP30 |  |
| CFAP44 | FLNA |  |
| MYO1B | PPP6C |  |
| CUL3 | GINS3 |  |
| CERCAM | EIF4E |  |
| GID8 | DCTN4 |  |
| WDR1 | VDAC3 |  |
| GOLIM4 | RPS24 |  |
| ICAM5 | RPS20 |  |
| PPCS | TPI1 |  |
| PFKP | RBMXL1 |  |
| VWA8 | FLNB |  |
| MYO1C | CYB5R3 |  |
| PIGR | RABL6 |  |
| YWHAQ | PCYT2 |  |
| PODN | DPYSL2 |  |
| SCYL1 | DPYSL3 |  |
| RPS15 | CALU |  |
| PAMR1 | HSPE1-MOB4 |  |
| CUL1 | SLC16A3 |  |
| SETD7 | BZW1 |  |
| RGS19 | AP2A2 |  |
| HSPD1 | RAB5C |  |
| UMPS | RPS3 |  |
| ABCA8 | TOR1AIP1 |  |
| VANGL1 | PPME1 |  |
| CCDC22 | COPZ1 |  |
| ACBD3 | RANGAP1 |  |
| GOT1 | CCL5 |  |
| MACF1 | STIP1 |  |
| FLOT1 | HMOX2 |  |
| DHX30 | TPT1 |  |
| STK25 | FAT4 |  |
| ECI2 | PTPRK |  |
| TBC1D2 | TUBB |  |
| TRAPPC2L | RA32-3 |  |
| NCAPH | AP2S1 |  |
| BCAR3 | GDIA1 |  |
| RBM14 | APOE |  |
| CCND1 | TUBA1C |  |
| PRPF19 | ACAT2 |  |
| GLG1 | ACLY |  |
| PEPD | RPSA |  |
| PTTG1IP | GPRC5B |  |
| BANF1 | RPS15 |  |
| KRT77 | SPARC |  |
| LDHA | LDHB |  |
| NCBP1 | RRP40 |  |
| IL19 | XRCC4 |  |
| ADAMTS12 | SEC24D |  |
| DYNC1I2 | TSKU |  |
| INTS1 | NRBF2 |  |
| FZD2 | CPSF3 |  |
| PSMD7 | JAK1 |  |
| TMPO | CNOT7 |  |
| ATG4B | UAP1 |  |
| RPL36AL | CYFIP1 |  |
| SMTN | DNAJC13 |  |
| OFD1 | PAXX |  |
| KIAA1468 | SCYL1 |  |
| MAPKAPK2 | SMARCC2 |  |
| TNFAIP3 | PTPN11 |  |
| MYH10 | WDR48 |  |
| ANKRD50 | AHNAK |  |
| PSME4 | TAOK3 |  |
| EXT2 | CRAT |  |
| TUBBP2 | MYO18A |  |
| KIAA0368 | DCTN2 |  |
| TANGO6 | SDCBP |  |
| PTK2 | PACSIN2 |  |
| CFB | ZIP14 |  |
| CFI | TACC1 |  |
| RPS2P17 | PRPSAP2-6 |  |
| LRRC15 | C1R |  |
| FAM171A1 | PFKM |  |
| HEXA | ADD1 |  |
| CACNA2D4 | SMARCB1 |  |
| CAMK1 | ZC3HAV1 |  |
| MLKL | SLC44A2 |  |
| MYH14 | CANX |  |
| PALLD | HNRNPH1 |  |
| KIF15 | OXCT1 |  |
| Sep-02 | ECMA9 |  |
| EVA1B | CACNA2D3 |  |
| KHDRBS1 | KIF23 |  |
| RALGPS2 | GREM1 |  |
| MECP2 | LARP7 |  |
| RAB5A | CPVL |  |
| DDX21 | LIMS4 |  |
| FARSB | SUMO1 |  |
| PRSS23 | NDRG1 |  |
| GNAI1 | SPTAN1 |  |
| CSK | AB13BP |  |
| PIP4K2C | PFN1 |  |
| FERMT2 | CLIP1 |  |
| SQLE | IARS1 |  |
| RPL10AP9 | SYPL1 |  |
| CTNND1 | TCERG1 |  |
| DCUN1D5 | JAMA |  |
| TBK1 | STAT1 |  |
| IL7 | MEMO1 |  |
| MEMO1 | HNRNPL |  |
| GNA13 | SERPINE1 |  |
| NUDT16L1 | ACAT1 |  |
| UFL1 | CDC42BPA |  |
| ANAPC7 | CLIP1 |  |
| RDH14 | PAICS |  |
| HEATR1 | MACF1 |  |
| GALT | EIF3I |  |
| ACLY | ITGA6 |  |
| HBE1 | GDI1 |  |
| NF1 | COL6A1 |  |
| ATP6AP2 | FGF2 |  |
| SGK3 | TNPO1 |  |
| ALDH16A1 | PRPSAP1 |  |
| FARP1 | VAPA |  |
| HIF1AN | CUL3 |  |
| HERC4 | CPNE1 |  |
| ENO2 | FSN |  |
| PPIAP31 | SERPINB9 |  |
| MAN1B1 | LAMA1 |  |
| NDRG3 | THOC4 |  |
| HCFC1 | PSMD14 |  |
| XPO5 | SAP18 |  |
| CSPG4 | TUBB3 |  |
| DIP2C | KHDRBS1 |  |
| NUP98 | CBX1 |  |
| UNC119B | COMMD3 |  |
| SGMS2 | PEF1 |  |
| C1R | SNAPIN |  |
| EIF4G1 | TTC37 |  |
| HSPA9 | MESD |  |
| EIF4E | NELFB |  |
| LAMB1 | NOP58 |  |
| MAP2K3 | IARS2 |  |
| ACO1 | EVA1B |  |
| ARL8B | RCN3 |  |
| EIF3B | NFKB |  |
| PLTP | TINAGL1 |  |
| PPIAP22 | HECTD3 |  |
| SFRP4 | CPSF7 |  |
| S100A10 | HNRNPUL1 |  |
| SNX5 | CHAMP1 |  |
| RPS25 | RBM17 |  |
| HNRNPF | CHST14 |  |
| NLN | RPIA |  |
| CCDC47 | PATL1 |  |
| KATNAL2 | AGRN |  |
| USP8 | CARM1 |  |
| TMEM51 | NCKAP1 |  |
| POP1 | PGRMC1 |  |
| ADH5 | QSOX1 |  |
| PTPRE | CASK |  |
| EIF6 | DHX15 |  |
| POLD2 | CYB5B |  |
| SLC2A1 | SEC24A |  |
| DOCK7 | HSPA8 |  |
| SPTBN1 | LAMP1 |  |
| NLRP8 | GRP94 |  |
| FST | CBR1 |  |
| GDF3 | ARF4 |  |
| GFPT2 | ECHS1 |  |
| KRT5 | CMPK1 |  |
| CD81 | TALDO1 |  |
| LOXL2 | COL18A1 |  |
| PYCR1 | TCP1 |  |
| ITIH4 | PPT1 |  |
| AKR7L | RAB7A |  |
| EXOSC2 | CAPZA1 |  |
| PCYT1A | PSA |  |
| ALDH3A2 | RAB15 |  |
| RAC2 | ACTC1 |  |
| CDK11A | FKBP52 |  |
| RPL30 | CSTF1 |  |
| NUCB1 | DHX9 |  |
| DHX38 | RPTPS |  |
| AURKA | FLNC |  |
| TBCA | RAB35 |  |
| UBA5 | EXT1 |  |
| ACTN1 | KTN1 |  |
| LIMK1 | H2AC21 |  |
| STXBP3 | MB21D2 |  |
| SFRP1 | DCBLD1 |  |
| CC2D1B | NPEPL1 |  |
| PPP2R1B | PSPC1 |  |
| RAB5C | ANP32B |  |
| DSCR3 | DDX17 |  |
| COG5 | IGSF8 |  |
| F2 | CRIP1 |  |
| NUP205 | PGAM5 |  |
| TGOLN2 | RPRD1A |  |
| KRAS | TPR2 |  |
| C5orf24 | UBE2Z |  |
| GDI1 | LRRC40 |  |
| RAB2A | NHP2 |  |
| CPS1 | MORF4L1 |  |
| NID1 | PLXNA1 |  |
| RPS10 | CFAP20 |  |
| NDUFB10 | HLAC |  |
| TTC37 | ITSN1 |  |
| APEX1 | ROCK2 |  |
| C9 | SLIT2 |  |
| VCPIP1 | IL7R |  |
| PTPN9 | VARS1 |  |
| CKAP4 | MDC1 |  |
| SDHB | HNRNPA2B1 |  |
| UBR5 | LRRC17 |  |
| LRRC40 | PMPCB |  |
| TWF1 | PLEC |  |
| ITGA11 | TNC |  |
| ATP6V0D1 | CD44 |  |
| ENO1 | ANKRD13A |  |
| GJA1 | TPP2 |  |
| UBR4 | CARHSP1 |  |
| TJP1 | CSNK1D |  |
| ZNHIT2 | FLII |  |
| MSH6 | SPAG9 |  |
| BGN | LIPG |  |
| PSMA7 | HNRNPUL1 |  |
| COL6A1 | PLD3 |  |
| EPHB1 | PTBP1 |  |
| ARMS2 | MAN2B1 |  |
| ADAMTS13 | COMMD7 |  |
| HSD17B4 | PLCG1 |  |
| FBLN5 | RAE1 |  |
| TRIM32 | PDXK |  |
| KDSR | UGDH |  |
| RAC3 | SEC24B |  |
| DAZAP1 | FAT1 |  |
| PKN2 | RANBP2 |  |
| ITGAL | EFEMP1 |  |
| STX6 | CAMK1 |  |
| RNF40 | EXOC1 |  |
| GNL1 | SEC23A |  |
| TMEM33 | NID2 |  |
| COL1A2 | CDC42 |  |
| PGP | GNPDA1 |  |
| IPO11 | SART3 |  |
| TMBIM1 | MYOF |  |
| WDR18 | ANXA11 |  |
| PLEKHA2 | KPNA6 |  |
| SNRNP200 | NASP |  |
| TRAPPC5 | SFPQ |  |
| GPC6 | HYOU1 |  |
| FAM168B | USP9X |  |
| PSMD1 | POLB |  |
| CSRP2 | PICALM |  |
| MYL6B | SH3GLB1 |  |
| RPS4X | BRO1 |  |
| IRF2BPL | UCHL5 |  |
| RPS12 | RO60 Y |  |
| DPYS | GCA |  |
| IMPDH2 | IMPDH2 |  |
| RAP1A | ANLN XI |  |
| GTF2A2 | SEPT 7 |  |
| KIAA1033 | SLC44A1 |  |
| DYNC1LI2 | LDHB |  |
| PGM1 | GDF11 |  |
| ARRDC1 | PKM |  |
| ARHGEF2 | FMNL1 |  |
| KRT79 | ITPA |  |
| ASCC2 | NDRG3 |  |
| MMP1 | FHL1 |  |
| FAF2 | HCFC1 |  |
| DDX1 | NIBAN1 |  |
| HSPA8 | THBS3 |  |
| GOT2 | TNFSF4 |  |
| GSTM3 | FBLN7 |  |
| MAEA | RAB3GAP1 |  |
| DNAAF5 | MGAT5 |  |
| AARSD1 | LPP |  |
| RABGAP1L | MELTF |  |
| ARL2 | DCUN1D1 |  |
| C7 | GMPS |  |
| WDR81 | RAI14 |  |
| BRIX1 | SEMA5A |  |
| NCAM2 | SLC29A1 |  |
| PSMD4 | SERPINB6 |  |
| S100P | LOC111313822 |  |
| HSPA12A | SEPTIN 7 |  |
| PROSC | AGO2 |  |
| PPP6C | UBE2V2 |  |
| UQCRC1 | CDKN2A |  |
| ARMT1 | PPP2R4 |  |
| CYB5R3 | NIBAN2 |  |
| NHLRC2 | CUL2 |  |
| GNAT3 | EXT2 |  |
| RABGAP1 | CHID1 |  |
| BAG6 | IPO5 |  |
| SLC44A1 | DNAJC3 |  |
| USP5 | NEO1 |  |
| PSMB4 | MYOIP3 |  |
| SMAP1 | PAFAH1B1 |  |
| MYO18A | RAEBP1 |  |
| ACSL1 | AP2B1 |  |
| RPS2 | KRT15 |  |
| EIF3K | SDF2 |  |
| PHKB | RNMT |  |
| ATG7 | NAPG |  |
| RPS10P11 | AP2S1 |  |
| KLC2 | AP2A1 |  |
| GLO1 | HSPBP1 |  |
| TALDO1 | CDC37 |  |
| COPB2 | N6 AMP LYASE |  |
| ARFGEF1 | EWSR1 |  |
| RTN1 | PITPNB |  |
| VPS37C | COL4A5 |  |
| IARS2 | MSN X1 |  |
| KCTD11 | ARFIP1 |  |
| RPS17 | BDH2 |  |
| EMILIN1 | DHX30 |  |
| WTH3DI | ROBO1 |  |
| LIG1 | GTF2F2 |  |
| LOX | COL12A1 |  |
| KRT8 | UTRN |  |
| SPARC | TNFAIP2 |  |
| DNAJA2 | CIT |  |
| CTSF | PPP1CC |  |
| RPTOR | CALCOCO1 |  |
| NUDT16 | EEA1 |  |
| VCL | SEC24C |  |
| CAND1 | ECE1 |  |
| SEC24C | UBR4 |  |
| DPYSL3 | PHGDH |  |
| FOXP1 | PTPRF |  |
| DCTN1 | SNX24 |  |
| NEDD4 | PAM |  |
| CAPG | ARL15 |  |
| FAM198B | TKFC |  |
| TJP2 | PGM2L1 |  |
| KDM1A | LTBP3 |  |
| ROCK1 | NUMA1 |  |
| TUBBP6 | NUDC |  |
| PNPLA6 | COL11A1 |  |
| HMGCS1 | TXLNA |  |
| TRIP12 | DNM3 |  |
| GCLC | GCLM |  |
| HPCAL1 | FUBP1 |  |
| TRIM38 | FUBP1 |  |
| TREML5P | VAMP3 |  |
| FAM114A1 | GFAT1 |  |
| METAP1 | PPM1B |  |
| LOXL1 | DOCK10 |  |
| BCR | UPF0104 |  |
| RABGEF1 | LZTFL1 |  |
| OSBPL2 | WNT5A |  |
| GTPBP2 | RBJP |  |
| SF3B2 | COMMD8 |  |
| ARPC2 | NCAPG |  |
| RNF213 | WDR1 |  |
| NPM1 | SDHA |  |
| RIPK1 | ADAMTS12 |  |
| EEA1 | PRIM2 |  |
| FZD6 | DST |  |
| NAA16 | PODNL1 |  |
| PHKA2 | WTAP |  |
| HELZ | NUB1 |  |
| COL3A1 | TPST1 |  |
| RAB35 | CDK13 |  |
| C3 | LY96/MD-2 |  |
| HDAC5 | EIFE3 |  |
| FKBP10 | GAPVD1 |  |
| IST1 | FAT3 |  |
| CTR9 | ARRB1 |  |
| ITPRIPL2 | PAFAH1B2 |  |
| EIF4A1 | AHNAK |  |
| PELP1 | STX2 |  |
| UBE2D2 | MON2 |  |
| TIA1 | ERC1 |  |
| RAI14 | LRP1 |  |
| NXF1 | VPS29 |  |
| LEO1 | BCAT1 |  |
| EEF1E1 | SLC7A1 |  |
| GALNT5 | MST3 |  |
| FDPS | COCH |  |
| NPR2 | FKBP3 |  |
| HGD | ACTN1 |  |
| RANBP3 | GLCE |  |
| ARHGEF7 | DNAJC17 |  |
| WDR77 | TJP1 |  |
| MARS | Ta-Gpg |  |
| CS | USP7 |  |
| MTMR1 | ANKFY1 |  |
| RPL13 | TNFAIP1 |  |
| EZR | ASPSCR1 |  |
| ZC3H7B | RAB31 |  |
| RMND5A | ATP5A1 |  |
| PTPRS | MAP1S |  |
| PPIP5K2 | PPP5C |  |
| CMBL | PSG4 |  |
| TTN | EPS15L1 |  |
| CNNM4 | UPF1 |  |
| DDX3Y | UBA52 |  |
| CYFIP1 | ACTN4 |  |
| HLA-C | GNAS |  |
| MCM4 | SRC |  |
| TRIM22 | LOC118846801 |  |
| NAV3 | UBA1 |  |
| RPL29P12 | EPB41L2 |  |
| IQGAP1 | CPSF1 |  |
| PGRMC1 | GRHPR |  |
| NBEAL1 | ADD3 |  |
| AACS | BCAS3 |  |
| CFTR | ENAH |  |
| TSPAN4 | ILK1 |  |
| CCR5 | SmMYB113 |  |
| GAA | SCYL1 |  |
| IL3 | FACT |  |
| TNC | LOC142304469 |  |
| NEDD4L | RACGAP1 |  |
| TRAPPC8 | COMMD4 |  |
| PEF1 | CEMIP |  |
| RPS6 | UBE3A |  |
| CLDN1 | XRCC1 |  |
| TBCE | EIF5A |  |
| LRBA | FLOT2 |  |
| HIST4H4 | NAGLU |  |
| SPAG9 | P4HB |  |
| OXSR1 | PSMC5 |  |
| NFKB1 | RPN5A |  |
| PLS3 | ELAC2 |  |
| ANKRD13A | STAT3 |  |
| CASP14 | RAB34 |  |
| RPL12P14 | TRIM28 |  |
| AKR1E2 | GTF2F1 |  |
| ARAP3 | ISYNA1 |  |
| NCAM1 | TMEM158 |  |
| SLC7A5 | ACSL4 |  |
| SCRIB | SEPTIN 2 |  |
| SLITRK4 | MAP4 ISO XI |  |
| GNB2L1 | CTPS1 |  |
| FADD | (FDPS) |  |
| POFUT2 | NFASC |  |
| COPS3 | DIAPH1 |  |
| GTF3C3 |  |  |
| KRT14 |  |  |
| DNAJC11 |  |  |
| RBP1 |  |  |
| SCUBE3 |  |  |
| UBE2V1 |  |  |
| EIF3H |  |  |
| CYB5B |  |  |
| SEC14L4 |  |  |
| PMVK |  |  |
| PDXK |  |  |
| RAB27B |  |  |
| ATP13A3 |  |  |
| HNRNPDL |  |  |
| PDS5A |  |  |
| IRAK4 |  |  |
| NAA25 |  |  |
| STON2 |  |  |
| ALDOB |  |  |
| MCTS1 |  |  |
| WASL |  |  |
| COG2 |  |  |
| EFR3A |  |  |
| ARHGEF1 |  |  |
| GNS |  |  |
| PLD1 |  |  |
| HMGA2 |  |  |
| HYI |  |  |
| MRC2 |  |  |
| CD276 |  |  |
| POLR1B |  |  |
| MTMR6 |  |  |
| LAMTOR1 |  |  |
| SPATA5 |  |  |
| FAM134C |  |  |
| RNPEP |  |  |
| MGRN1 |  |  |
| FTSJ1 |  |  |
| ATP13A1 |  |  |
| AKR1B1 |  |  |
| BCAM |  |  |
| PRKCDBP |  |  |
| GMPPB |  |  |
| FBP1 |  |  |
| SNRPA1 |  |  |
| PRRC2C |  |  |
| ABCD3 |  |  |
| POP4 |  |  |
| PSMD6 |  |  |
| NARFL |  |  |
| SLC15A4 |  |  |
| EIF2B4 |  |  |
| EML4 |  |  |
| MAPKAP1 |  |  |
| RIN1 |  |  |
| SURF4 |  |  |
| PDGFC |  |  |
| ARHGDIA |  |  |
| LARS |  |  |
| PSMD12 |  |  |
| GDF11 |  |  |
| DNPEP |  |  |
| RARRES1 |  |  |
| MTCL1 |  |  |
| FLG2 |  |  |
| ZNF134 |  |  |
| ILF2 |  |  |
| ALDH1A1 |  |  |
| IGLC2 |  |  |
| PLEKHO1 |  |  |
| FAM129A |  |  |
| RPS10P22 |  |  |
| NNMT |  |  |
| HMOX2 |  |  |
| CD59 |  |  |
| MGAT1 |  |  |
| YTHDC2 |  |  |
| SNRPG |  |  |
| TXN |  |  |
| PBXIP1 |  |  |
| RICTOR |  |  |
| NAA15 |  |  |
| CTSC |  |  |
| SNX6 |  |  |
| LRRC59 |  |  |
| ALB |  |  |
| KANK2 |  |  |
| GAPDH |  |  |
| NUBP2 |  |  |
| C8B |  |  |
| MYD88 |  |  |
| ZYX |  |  |
| SLC12A2 |  |  |
| PLOD2 |  |  |
| MAPKAPK3 |  |  |
| PLG |  |  |
| DNM3 |  |  |
| NUDT21 |  |  |
| RPS6KB1 |  |  |
| HSP90AB4P |  |  |
| GOPC |  |  |
| HMGCS2 |  |  |
| TAGLN |  |  |
| NOLC1 |  |  |
| SAMD4A |  |  |
| SERPINB7 |  |  |
| SNRPB |  |  |
| ARHGAP23 |  |  |
| UEVLD |  |  |
| HIST1H4J |  |  |
| TMED9 |  |  |
| GNB2 |  |  |
| VDAC2 |  |  |
| VPS13C |  |  |
| SEC61A1 |  |  |
| SRSF2 |  |  |
| ALDH18A1 |  |  |
| A2M |  |  |
| ANKRD28 |  |  |
| CAV1 |  |  |
| TRMT61A |  |  |
| EEF1B2 |  |  |
| SERPINA7 |  |  |
| CPNE7 |  |  |
| BHMT2 |  |  |
| AP2A1 |  |  |
| PSMC4 |  |  |
| INPP5K |  |  |
| CTBP1 |  |  |
| HGF |  |  |
| VAC14 |  |  |
| GSTM1 |  |  |
| COG1 |  |  |
| PYGB |  |  |
| TRIM16 |  |  |
| CCR4 |  |  |
| DDB1 |  |  |
| RPS8 |  |  |
| HEG1 |  |  |
| CFL1 |  |  |
| DSG1 |  |  |
| AIMP2 |  |  |
| RPL11 |  |  |
| CDK7 |  |  |
| TKT |  |  |
| TIFAB |  |  |
| C14orf166 |  |  |
| NBAS |  |  |
| KRT2 |  |  |
| MYL6 |  |  |
| ANP32B |  |  |
| PURA |  |  |
| HIST1H2BA |  |  |
| RPL23 |  |  |
| UAP1L1 |  |  |
| BLVRB |  |  |
| PRDX6 |  |  |
| SBDS |  |  |
| SCFD2 |  |  |
| UNC45A |  |  |
| MAP1S |  |  |
| PTPN11 |  |  |
| CTSG |  |  |
| UBAP1 |  |  |
| GLS |  |  |
| FUCA1 |  |  |
| COL4A3 |  |  |
| PIK3C3 |  |  |
| CMTM3 |  |  |
| RPL32 |  |  |
| PRR12 |  |  |
| GBE1 |  |  |
| RALB |  |  |
| NOP56 |  |  |
| DNAH6 |  |  |
| DHX57 |  |  |
| HIST1H1B |  |  |
| LAP3 |  |  |
| NAA35 |  |  |
| CLCN7 |  |  |
| POLR2A |  |  |
| VIM |  |  |
| MAPK10 |  |  |
| VTN |  |  |
| UFD1L |  |  |
| ATP8B3 |  |  |
| MLLT4 |  |  |
| PAAF1 |  |  |
| FAU |  |  |
| MAN1A2 |  |  |
| PPP6R3 |  |  |
| NOTCH3 |  |  |
| UGGT2 |  |  |
| TRIM21 |  |  |
| WDR61 |  |  |
| CAB39 |  |  |
| HNRNPLL |  |  |
| RND3 |  |  |
| PCDH7 |  |  |
| ANPEP |  |  |
| PNKP |  |  |
| DUS2 |  |  |
| GOLGA2 |  |  |
| PTGES3 |  |  |
| MGEA5 |  |  |
| CDIPT |  |  |
| CD82 |  |  |
| TTL |  |  |
| BCCIP |  |  |
| SEH1L |  |  |
| ITGB1 |  |  |
| CHP1 |  |  |
| GNG10 |  |  |
| HEBP2 |  |  |
| IL1RL2 |  |  |
| HAS2 |  |  |
| LSM1 |  |  |
| SRR |  |  |
| DCTN4 |  |  |
| PI4K2A |  |  |
| RPLP0P6 |  |  |
| CLTA |  |  |
| COL14A1 |  |  |
| GMPPA |  |  |
| LRRC7 |  |  |
| PXN |  |  |
| SNX1 |  |  |
| P3H4 |  |  |
| SLC38A2 |  |  |
| HECTD1 |  |  |
| RFC4 |  |  |
| ALDOA |  |  |
| GGH |  |  |
| CARM1 |  |  |
| NMRAL1 |  |  |
| MON1B |  |  |
| CMPK1 |  |  |
| SLC17A5 |  |  |
| GMPR2 |  |  |
| PXK |  |  |
| APOM |  |  |
| SLC22A2 |  |  |
| SMC4 |  |  |
| DPYD |  |  |
| PSD3 |  |  |
| CDCP1 |  |  |
| COMMD3 |  |  |
| APPL2 |  |  |
| MKRN2 |  |  |
| LONP1 |  |  |
| MRPS27 |  |  |
| LAMB2 |  |  |
| VPS11 |  |  |
| FBL |  |  |
| PARP4 |  |  |
| CAPZB |  |  |
| RPL10AP6 |  |  |
| AHCYL1 |  |  |
| SLC30A7 |  |  |
| PSMA6 |  |  |
| FNTA |  |  |
| ATG3 |  |  |
| SCAMP2 |  |  |
| SARNP |  |  |
| TRMT1 |  |  |
| AP2A2 |  |  |
| SMAD1 |  |  |
| OGFR |  |  |
| CD63 |  |  |
| RAB32 |  |  |
| COASY |  |  |
| PFKFB4 |  |  |
| LRP8 |  |  |
| SAMHD1 |  |  |
| ACTR1A |  |  |
| COL12A1 |  |  |
| NT5DC2 |  |  |
| PDDC1 |  |  |
| ADGRL4 |  |  |
| HBA1 |  |  |
| PARP1 |  |  |
| FLT1 |  |  |
| SEMA5A |  |  |
| TUBB4A |  |  |
| PEAK1 |  |  |
| TRNT1 |  |  |
| EXOC8 |  |  |
| PRDX1 |  |  |
| DCHS2 |  |  |
| ACAA1 |  |  |
| SLC2A6 |  |  |
| EIF3L |  |  |
| LRPPRC |  |  |
| ABI2 |  |  |
| ATP6V1B2 |  |  |
| PDPK1 |  |  |
| UCHL1 |  |  |
| MMP10 |  |  |
| RPL27A |  |  |
| GEMIN8 |  |  |
| HDAC2 |  |  |
| KIF13B |  |  |
| TAB1 |  |  |
| OAT |  |  |
| DOCK10 |  |  |
| APP |  |  |
| NT5C2 |  |  |
| TM9SF1 |  |  |
| TRIOBP |  |  |
| HSP90AA2P |  |  |
| ANXA4 |  |  |
| NUDCD2 |  |  |
| SERPINE1 |  |  |
| MGMT |  |  |
| FRG1 |  |  |
| TGFBI |  |  |
| SDSL |  |  |
| RHOBTB3 |  |  |
| CUL5 |  |  |
| SRGAP1 |  |  |
| HNRNPA0 |  |  |
| IL6ST |  |  |
| TMED10 |  |  |
| PHLDB2 |  |  |
| BST1 |  |  |
| CDK1 |  |  |
| MFHAS1 |  |  |
| CKLF |  |  |
| TMX2 |  |  |
| NUDT11 |  |  |
| PPID |  |  |
| GRHPR |  |  |
| SGCD |  |  |
| ICAM1 |  |  |
| SAMM50 |  |  |
| MCC |  |  |
| HNRNPH3 |  |  |
| SERPINC1 |  |  |
| TSKU |  |  |
| YIPF5 |  |  |
| CTNNBL1 |  |  |
| CPPED1 |  |  |
| RALA |  |  |
| MOS |  |  |
| TMEM43 |  |  |
| TEX10 |  |  |
| SFN |  |  |
| OPRM1 |  |  |
| PCYOX1 |  |  |
| ZDHHC8 |  |  |
| MAMDC2 |  |  |
| LMO7 |  |  |
| COPS2 |  |  |
| CNTN1 |  |  |
| LYPLA2 |  |  |
| EXOC3 |  |  |
| SLC12A9 |  |  |
| TIMM8A |  |  |
| RPS10P4 |  |  |
| MMP2 |  |  |
| IGLV4-3 |  |  |
| NUP188 |  |  |
| IDH3A |  |  |
| KCTD10 |  |  |
| S100A8 |  |  |
| RPL6 |  |  |
| API5 |  |  |
| VPS25 |  |  |
| ARAF |  |  |
| TMEM55A |  |  |
| MCM7 |  |  |
| RBBP7 |  |  |
| C19orf26 |  |  |
| ALDH9A1 |  |  |
| AP3S1 |  |  |
| IGSF8 |  |  |
| IGLV3-21 |  |  |
| FBN2 |  |  |
| ANXA1 |  |  |
| UNC79 |  |  |
| TPT1 |  |  |
| SYT9 |  |  |
| COMT |  |  |
| NRAS |  |  |
| RAB11B |  |  |
| FAM49B |  |  |
| ERBB2IP |  |  |
| VTI1B |  |  |
| APOB |  |  |
| SLK |  |  |
| POSTN |  |  |
| SCRN1 |  |  |
| FGF18 |  |  |
| HIST1H4H |  |  |
| FBXL18 |  |  |
| LOC646127 |  |  |
| RPL12P35 |  |  |
| SESN2 |  |  |
| WDR5 |  |  |
| ERGIC3 |  |  |
| ECE1 |  |  |
| HP |  |  |
| CTNNA2 |  |  |
| UTP15 |  |  |
| CLUH |  |  |
| CDC23 |  |  |
| FAM3B |  |  |
| CPSF1 |  |  |
| SMARCC1 |  |  |
| PDHB |  |  |
| TUBGCP6 |  |  |
| PDCD6IP |  |  |
| MYCBP2 |  |  |
| HNRNPR |  |  |
| BTAF1 |  |  |
| ADAM10 |  |  |
| RANBP2 |  |  |
| CBR1 |  |  |
| LPXN |  |  |
| KRT78 |  |  |
| NAA10 |  |  |
| ATXN2L |  |  |
| ENG |  |  |
| CALR |  |  |
| NDE1 |  |  |
| TRIO |  |  |
| EIF3C |  |  |
| CDC5L |  |  |
| CTDNEP1 |  |  |
| LGR6 |  |  |
| AP3D1 |  |  |
| SSR4 |  |  |
| FGFRL1 |  |  |
| HSPH1 |  |  |
| NTPCR |  |  |
| EIF3E |  |  |
| SLC26A2 |  |  |
| RPS18P5 |  |  |
| ANXA2P1 |  |  |
| ALDH1L2 |  |  |
| BAG3 |  |  |
| CCDC15 |  |  |
| NBL1 |  |  |
| ACTL6A |  |  |
| GREM1 |  |  |
| ARSG |  |  |
| FBXW8 |  |  |
| MANBA |  |  |
| INSR |  |  |
| H2AFX |  |  |
| PGAM5 |  |  |
| ABI1 |  |  |
| LDHB |  |  |
| UGCG |  |  |
| SCIN |  |  |
| COL5A1 |  |  |
| DNAJB2 |  |  |
| ALDH1B1 |  |  |
| PPFIA1 |  |  |
| NT5E |  |  |
| VEPH1 |  |  |
| PVRL3 |  |  |
| IMP3 |  |  |
| RAB6C |  |  |
| FTO |  |  |
| CDK16 |  |  |
| INPPL1 |  |  |
| CASK |  |  |
| STAM |  |  |
| DUSP3 |  |  |
| CALU |  |  |
| BTN2A1 |  |  |
| RECQL |  |  |
| RPL36 |  |  |
| GTPBP4 |  |  |
| H3F3A |  |  |
| ATP1B3 |  |  |
| METRNL |  |  |
| PNP |  |  |
| AKAP11 |  |  |
| SERPINF2 |  |  |
| PAK2 |  |  |
| GALE |  |  |
| PLD2 |  |  |
| IGFBP3 |  |  |
| MVB12A |  |  |
| PUM2 |  |  |
| NF2 |  |  |
| DRG2 |  |  |
| NOTCH2 |  |  |
| ATL1 |  |  |
| COL4A2 |  |  |
| CLEC11A |  |  |
| ABCE1 |  |  |
| CYB5R1 |  |  |
| TRADD |  |  |
| ADAM12 |  |  |
| RPS3 |  |  |
| PTGFRN |  |  |
| SPC25 |  |  |
| CRYZ |  |  |
| F3 |  |  |
| BPGM |  |  |
| SAP18 |  |  |
| RPL7 |  |  |
| SPRY4 |  |  |
| ATP6V1A |  |  |
| DIS3 |  |  |
| POFUT1 |  |  |
| NXN |  |  |
| EHD4 |  |  |
| SYT1 |  |  |
| NBN |  |  |
| PPP2R4 |  |  |
| FIG4 |  |  |
| GCLM |  |  |
| U2SURP |  |  |
| IREB2 |  |  |
| PDCD4 |  |  |
| HECTD4 |  |  |
| MXRA8 |  |  |
| FIBP |  |  |
| STX2 |  |  |
| GALNT10 |  |  |
| EXOSC4 |  |  |
| SMPD1 |  |  |
| TRA2B |  |  |
| WDR91 |  |  |
| CHSY1 |  |  |
| RB1 |  |  |
| AMPD2 |  |  |
| RRM2B |  |  |
| INPP5A |  |  |
| TPM3 |  |  |
| LARP1B |  |  |
| GNAS |  |  |
| SMCHD1 |  |  |
| ASS1 |  |  |
| TPI1P1 |  |  |
| DYRK1A |  |  |
| CCDC132 |  |  |
| FCN3 |  |  |
| APCS |  |  |
| TMED7 |  |  |
| ROGDI |  |  |
| RXRB |  |  |
| DFFA |  |  |
| ARHGAP17 |  |  |
| IFITM2 |  |  |
| DSP |  |  |
| DDX19A |  |  |
| BPIFB1 |  |  |
| DPYSL4 |  |  |
| ARHGEF11 |  |  |
| RDX |  |  |
| EPHB4 |  |  |
| UBE3C |  |  |
| UBE2D1 |  |  |
| WDR37 |  |  |
| PCOLCE |  |  |
| RNMT |  |  |
| RBM39 |  |  |
| EDEM3 |  |  |
| GART |  |  |
| PSMD8 |  |  |
| FAM160B1 |  |  |
| SOD2 |  |  |
| MAN1A1 |  |  |
| RIMBP3 |  |  |
| ARPC1B |  |  |
| TNFAIP2 |  |  |
| SUGT1 |  |  |
| LUM |  |  |
| CPNE8 |  |  |
| CCL2 |  |  |
| IGHA1 |  |  |
| PRIM2 |  |  |
| MSI2 |  |  |
| MBD3 |  |  |
| SLC2A14 |  |  |
| METAP2 |  |  |
| RPS10P7 |  |  |
| DCK |  |  |
| TXNDC17 |  |  |
| HK1 |  |  |
| BCLAF1 |  |  |
| NACA |  |  |
| CEMIP |  |  |
| AIDA |  |  |
| MAT2B |  |  |
| TCN1 |  |  |
| GNA11 |  |  |
| CDH11 |  |  |
| LCMT2 |  |  |
| ARMC9 |  |  |
| ACTB |  |  |
| THADA |  |  |
| PACS1 |  |  |
| CA12 |  |  |
| PDE12 |  |  |
| TNFRSF12A |  |  |
| RAB33B |  |  |
| PLK1 |  |  |
| PCDHGC3 |  |  |
| IPO13 |  |  |
| RPL12 |  |  |
| MARK2 |  |  |
| PGK1 |  |  |
| GIPC1 |  |  |
| MCM6 |  |  |
| CUL4B |  |  |
| SH3BGRL3 |  |  |
| YIF1A |  |  |
| RUVBL1 |  |  |
| FASN |  |  |
| LAMA1 |  |  |
| MOXD1 |  |  |
| AHNAK2 |  |  |
| BCAS3 |  |  |
| MON2 |  |  |
| PAPPA |  |  |
| RGN |  |  |
| DIAPH3 |  |  |
| BZW1 |  |  |
| ASB6 |  |  |
| RRAGA |  |  |
| ANXA7 |  |  |
| PSAP |  |  |
| STEAP3 |  |  |
| RPS2P11 |  |  |
| RAB43 |  |  |
| OSBPL8 |  |  |
| ANKHD1 |  |  |
| EIF4G2 |  |  |
| TYMP |  |  |
| SNTB2 |  |  |
| LDHAL6B |  |  |
| PRCP |  |  |
| H2AFV |  |  |
| DDT |  |  |
| PELO |  |  |
| RRP9 |  |  |
| XPOT |  |  |
| ANXA3 |  |  |
| XRCC1 |  |  |
| PRDM16 |  |  |
| TRHDE |  |  |
| DBNL |  |  |
| HEXB |  |  |
| TG |  |  |
| AP1B1 |  |  |
| PRKG1 |  |  |
| NRP2 |  |  |
| AQR |  |  |
| SPTLC1 |  |  |
| G3BP2 |  |  |
| ARPC1A |  |  |
| KRT19 |  |  |
| IL13 |  |  |
| SAA4 |  |  |
| LARS2 |  |  |
| SLC39A10 |  |  |
| ACADVL |  |  |
| GCN1L1 |  |  |
| LOC100130100 |  |  |
| EPS15L1 |  |  |
| GPSM1 |  |  |
| TYMS |  |  |
| TBC1D13 |  |  |
| XPR1 |  |  |
| PHB |  |  |
| RPL12P32 |  |  |
| PLEKHG3 |  |  |
| IGFALS |  |  |
| PLCD3 |  |  |
| RRBP1 |  |  |
| B2M |  |  |
| PPP2R2A |  |  |
| CDK13 |  |  |
| CKAP5 |  |  |
| EXTL2 |  |  |
| ESD |  |  |
| MAT1A |  |  |
| ETFB |  |  |
| ARL3 |  |  |
| STC2 |  |  |
| SACS |  |  |
| HSP90AA1 |  |  |
| YWHAE |  |  |
| LGALS1 |  |  |
| TMEM47 |  |  |
| YWHAEP5 |  |  |
| STRN4 |  |  |
| ETFA |  |  |
| CLK2 |  |  |
| GSTM2 |  |  |
| PSMD14 |  |  |
| PSPH |  |  |
| BAG5 |  |  |
| SHOC2 |  |  |
| SORD |  |  |
| CIRH1A |  |  |
| P3H3 |  |  |
| DEK |  |  |
| ZZEF1 |  |  |
| METTL13 |  |  |
| IPO5 |  |  |
| TROVE2 |  |  |
| CAP2 |  |  |
| RECK |  |  |
| ABCF1 |  |  |
| CRYM-AS1 |  |  |
| MYOF |  |  |
| VPS13A |  |  |
| DTYMK |  |  |
| KRT18P19 |  |  |
| UBA7 |  |  |
| NUP133 |  |  |
| CSNK1G1 |  |  |
| MAP3K1 |  |  |
| RTN4 |  |  |
| TF |  |  |
| YKT6 |  |  |
| POLR1C |  |  |
| EEF2 |  |  |
| TTC27 |  |  |
| DMXL1 |  |  |
| MFSD10 |  |  |
| LRRC8C |  |  |
| FAM175B |  |  |
| ANO6 |  |  |
| MARCKSL1 |  |  |
| SLC30A1 |  |  |
| RPL8 |  |  |
| ACTN4 |  |  |
| VRK2 |  |  |
| DAD1 |  |  |
| RPLP2 |  |  |
| NOMO1 |  |  |
| CDC123 |  |  |
| DDX3X |  |  |
| PDLIM7 |  |  |
| DUSP12 |  |  |
| LGALS3 |  |  |
| INTS6 |  |  |
| XRCC6 |  |  |
| SLC5A6 |  |  |
| CAT |  |  |
| PCM1 |  |  |
| SHC1 |  |  |
| HTRA1 |  |  |
| C17orf49 |  |  |
| VPS16 |  |  |
| EIF2S1 |  |  |
| LRRTM1 |  |  |
| DAB2 |  |  |
| GBA3 |  |  |
| KARS |  |  |
| RALY |  |  |
| EEF1D |  |  |
| GLOD4 |  |  |
| NAGLU |  |  |
| HNRNPC |  |  |
| RAB39B |  |  |
| ARPC4 |  |  |
| CFL2 |  |  |
| TXNDC5 |  |  |
| RPS6KA4 |  |  |
| HDLBP |  |  |
| TNIK |  |  |
| IMPDH1 |  |  |
| STAT5B |  |  |
| HAUS5 |  |  |
| RPS2P20 |  |  |
| PSMB8 |  |  |
| PI4KA |  |  |
| PTPN1 |  |  |
| TYRO3 |  |  |
| TPP2 |  |  |
| MAP1B |  |  |
| OCRL |  |  |
| STUB1 |  |  |
| GPX7 |  |  |
| SLC38A3 |  |  |
| CD248 |  |  |
| SCOC |  |  |
| LDLR |  |  |
| SQRDL |  |  |
| DCXR |  |  |
| TUBA1C |  |  |
| RPS15A |  |  |
| STX8 |  |  |
| ADSS |  |  |
| SYPL1 |  |  |
| SHANK3 |  |  |
| WBP2 |  |  |
| RRAGC |  |  |
| LRRC17 |  |  |
| VAMP7 |  |  |
| TANK |  |  |
| SRSF10 |  |  |
| RPS19 |  |  |
| ATP6V1E1 |  |  |
| EPHX2 |  |  |
| RAP2B |  |  |
| EMC2 |  |  |
| ARL6IP5 |  |  |
| GFRA3 |  |  |
| COPS7A |  |  |
| LRRC14 |  |  |
| LZTR1 |  |  |
| TSTA3 |  |  |
| HSPBP1 |  |  |
| NDUFS1 |  |  |
| TST |  |  |
| PHLDA2 |  |  |
| DENND5A |  |  |
| CCT7 |  |  |
| PPT1 |  |  |
| MED24 |  |  |
| AHCY |  |  |
| ZCCHC6 |  |  |
| NID2 |  |  |
| TPP1 |  |  |
| OXTR |  |  |
| SPTBN5 |  |  |
| GNE |  |  |
| TUBGCP3 |  |  |
| QTRT1 |  |  |
| ZNF503 |  |  |
| ARPC3 |  |  |
| COPS6 |  |  |
| RPSAP9 |  |  |
| HP1BP3 |  |  |
| LCN1 |  |  |
| GANAB |  |  |
| SPTLC2 |  |  |
| VCP |  |  |
| TUBA3E |  |  |
| PWP1 |  |  |
| FANCL |  |  |
| HIST1H4K |  |  |
| USP10 |  |  |
| CPB2 |  |  |
| GEMIN4 |  |  |
| GAPDHS |  |  |
| SLC27A5 |  |  |
| LRRC32 |  |  |
| DCLK2 |  |  |
| ACOT7 |  |  |
| RPS9 |  |  |
| PROCR |  |  |
| VAT1 |  |  |
| KIRREL |  |  |
| VPRBP |  |  |
| CEACAM8 |  |  |
| KRT27 |  |  |
| ANP32E |  |  |
| COL6A3 |  |  |
| NAT1 |  |  |
| FSCN1 |  |  |
| RPS10P13 |  |  |
| PLXNA3 |  |  |
| EXOC2 |  |  |
| GALK2 |  |  |
| GPX1 |  |  |
| DARS2 |  |  |
| HNRNPA2B1 |  |  |
| FKBP4 |  |  |
| VBP1 |  |  |
| SRSF6 |  |  |
| SERPINH1 |  |  |
| CBWD1 |  |  |
| CHMP3 |  |  |
| SYT11 |  |  |
| CDC42BPB |  |  |
| TERF1P2 |  |  |
| DKK3 |  |  |
| HEBP1 |  |  |
| IGF2BP3 |  |  |
| SLC6A6 |  |  |
| PRMT3 |  |  |
| IFITM3 |  |  |
| PC |  |  |
| EIF5B |  |  |
| ME1 |  |  |
| PDGFA |  |  |
| OSTF1 |  |  |
| STXBP5 |  |  |
| RAB34 |  |  |
| SNX12 |  |  |
| ENTPD4 |  |  |
| HPD |  |  |
| ACACA |  |  |
| RPSAP29 |  |  |
| GSTO1 |  |  |
| CRAT |  |  |
| JCHAIN |  |  |
| ERP44 |  |  |
| EFHD2 |  |  |
| PSMD3 |  |  |
| NDUFA9 |  |  |
| WNT8B |  |  |
| TP53BP1 |  |  |
| TUBB6 |  |  |
| SCRN2 |  |  |
| KRT18 |  |  |
| CDC16 |  |  |
| CD44 |  |  |
| CBS |  |  |
| LARP1 |  |  |
| TTC9C |  |  |
| ATP5L |  |  |
| RPA1 |  |  |
| TPD52L2 |  |  |
| FAM45A |  |  |
| RPL24 |  |  |
| PPP5C |  |  |
| SMC1A |  |  |
| DIRAS2 |  |  |
| GALNS |  |  |
| TP53RK |  |  |
| SIPA1L1 |  |  |
| TSPAN10 |  |  |
| S100A11 |  |  |
| PHLDA1 |  |  |
| LOC442497 |  |  |
| TBC1D2B |  |  |
| ITPR2 |  |  |
| ADAMTSL1 |  |  |
| DOCK11 |  |  |
| UBE2Z |  |  |
| FREM3 |  |  |
| DMBT1 |  |  |
| LRP10 |  |  |
| CNN2 |  |  |
| PPP1R18 |  |  |
| SULF1 |  |  |
| P3H1 |  |  |
| GYLTL1B |  |  |
| LRRC8A |  |  |
| AXL |  |  |
| PHB2 |  |  |
| CTPS1 |  |  |
| ANGPTL4 |  |  |
| XPNPEP1 |  |  |
| CDK2 |  |  |
| LNPEP |  |  |
| EHD1 |  |  |
| HUWE1 |  |  |
| PSMB2 |  |  |
| PLEKHG2 |  |  |
| MMP3 |  |  |
| AKAP13 |  |  |
| PICALM |  |  |
| EXOC7 |  |  |
| IFIT5 |  |  |
| TTLL3 |  |  |
| EIF2A |  |  |
| HIST2H2AB |  |  |
| CUL4A |  |  |
| MRE11A |  |  |
| MGAT4B |  |  |
| ITGB5 |  |  |
| SCP2 |  |  |
| KRT16P2 |  |  |
| CSE1L |  |  |
| SH3GLB1 |  |  |
| DDX20 |  |  |
| HOOK3 |  |  |
| RPL27 |  |  |
| EPG5 |  |  |
| GRM3 |  |  |
| RPL15 |  |  |
| TRAP1 |  |  |
| PUS1 |  |  |
| YARS |  |  |
| PNPT1 |  |  |
| TBPL1 |  |  |
| LMAN1 |  |  |
| LCMT1 |  |  |
| HSPD1P5 |  |  |
| DPP7 |  |  |
| NME1-NME2 |  |  |
| ZDHHC20 |  |  |
| LAMC1 |  |  |
| PLIN3 |  |  |
| EXOSC6 |  |  |
| EIF2D |  |  |
| SHMT1 |  |  |
| SEC61B |  |  |
| TSPAN14 |  |  |
| SPRYD3 |  |  |
| ATG9A |  |  |
| GPS1 |  |  |
| SIRT5 |  |  |
| COG8 |  |  |
| THSD4 |  |  |
| DIS3L2 |  |  |
| TREML2 |  |  |
| BAZ1A |  |  |
| CACNA2D1 |  |  |
| CLMP |  |  |
| DNAJC2 |  |  |
| RPS29 |  |  |
| DST |  |  |
| NAV1 |  |  |
| CHMP1B |  |  |
| SPRYD7 |  |  |
| PRPF8 |  |  |
| PML |  |  |
| NRP1 |  |  |
| PPP3CC |  |  |
| HNRNPUL1 |  |  |
| PSMA4 |  |  |
| XPO1 |  |  |
| HSP90B1 |  |  |
| STARD3NL |  |  |
| CMIP |  |  |
| ADAT3 |  |  |
| POLD1 |  |  |
| CPSF7 |  |  |
| PTPRK |  |  |
| RPS23 |  |  |
| SERPIND1 |  |  |
| DNASE1L1 |  |  |
| BDKRB1 |  |  |
| COL4A1 |  |  |
| KIAA1715 |  |  |
| CBLB |  |  |
| SNX18 |  |  |
| GDF1 |  |  |
| ABCA1 |  |  |
| OTC |  |  |
| TSN |  |  |
| PAPSS1 |  |  |
| ARHGAP31 |  |  |
| PLAU |  |  |
| FTH1 |  |  |
| PTPN12 |  |  |
| HIST2H4B |  |  |
| CHID1 |  |  |
| KHSRP |  |  |
| ASPH |  |  |
| TNS1 |  |  |
| DAPK1 |  |  |
| EHD2 |  |  |
| HAPLN1 |  |  |
| RPN2 |  |  |
| LAMTOR3 |  |  |
| RAB12 |  |  |
| LYZ |  |  |
| PPIL1 |  |  |
| PAPOLA |  |  |
| EIF4A2 |  |  |
| GNAI3 |  |  |
| DBN1 |  |  |
| XPO6 |  |  |
| PPP1CC |  |  |
| YWHAB |  |  |
| VEGFC |  |  |
| RPS27AP11 |  |  |
| OGT |  |  |
| SERPINB12 |  |  |
| CC2D1A |  |  |
| FN3KRP |  |  |
| C18orf8 |  |  |
| RAP2A |  |  |
| ANXA8 |  |  |
| GNAI2 |  |  |
| NUDCD1 |  |  |
| PSMD13 |  |  |
| ALDH6A1 |  |  |
| SYNGR2 |  |  |
| GRWD1 |  |  |
| XPO7 |  |  |
| HSPA2 |  |  |
| BSG |  |  |
| APOA1 |  |  |
| RPS7 |  |  |
| SLC1A5 |  |  |
| RARS |  |  |
| RPL22L1 |  |  |
| SSR1 |  |  |
| SRGAP2 |  |  |
| FXR1 |  |  |
| ACAA2 |  |  |
| PSIP1 |  |  |
| LAMA2 |  |  |
| IQGAP3 |  |  |
| KRT6C |  |  |
| DDR2 |  |  |
| ASB3 |  |  |
| G6PD |  |  |
| KRT7 |  |  |
| PEBP1 |  |  |
| UBB |  |  |
| HIST1H4D |  |  |
| RPS3A |  |  |
| DDR1 |  |  |
| RPL22 |  |  |
| IGHV4-31 |  |  |
| STAT3 |  |  |
| HPS6 |  |  |
| AGRN |  |  |
| PDIA6 |  |  |
| ARHGAP22 |  |  |
| HSP90AB1 |  |  |
| IFNGR1 |  |  |
| RAB22A |  |  |
| AASDHPPT |  |  |
| ATP2B1 |  |  |
| NADK2 |  |  |
| PARP9 |  |  |
| OTULIN |  |  |
| RAB6A |  |  |
| MAGED2 |  |  |
| KRT10 |  |  |
| DCBLD2 |  |  |
| TARDBP |  |  |
| RAB8B |  |  |
| LOC100133211 |  |  |
| PIK3CA |  |  |
| GLUD1 |  |  |
| GTPBP1 |  |  |
| RPL35 |  |  |
| NQO1 |  |  |
| MRTO4 |  |  |
| QPRT |  |  |
| AAK1 |  |  |
| RASA4 |  |  |
| FTL |  |  |
| ELMOD2 |  |  |
| CORO7 |  |  |
| RPL29P11 |  |  |
| AKR7A2 |  |  |
| PSMB6 |  |  |
| MYCBPAP |  |  |
| CCT4 |  |  |
| GIT2 |  |  |
| EPO |  |  |
| CARHSP1 |  |  |
| NSF |  |  |
| PTP4A1 |  |  |
| DENND4C |  |  |
| VPS37B |  |  |
| WDR82 |  |  |
| CLTCL1 |  |  |
| ZFYVE1 |  |  |
| PRKAR1A |  |  |
| DARS |  |  |
| ECM1 |  |  |
| PSAT1 |  |  |
| C7orf60 |  |  |
| PRPS1 |  |  |
| PARG |  |  |
| KRT72 |  |  |
| CDK5R2 |  |  |
| STK11IP |  |  |
| LRP6 |  |  |
| SLC25A24 |  |  |
| PON2 |  |  |
| TIMP3 |  |  |
| HBB |  |  |
| ABCB10 |  |  |
| KRT4 |  |  |
| BRAT1 |  |  |
| AKAP10 |  |  |
| NEU1 |  |  |
| MTMR2 |  |  |
| PRNP |  |  |
| HIST1H4F |  |  |
| PRDX2 |  |  |
| WDR70 |  |  |
| SIPA1 |  |  |
| AHRR |  |  |
| FGFR1 |  |  |
| F2R |  |  |
| PITRM1 |  |  |
| GALC |  |  |
| PPP4R1 |  |  |
| ARMC1 |  |  |
| HNRNPUL2 |  |  |
| ARHGAP1 |  |  |
| KIAA0196 |  |  |
| CNN3 |  |  |
| TEP1 |  |  |
| THY1 |  |  |
| SDC4 |  |  |
| RPS20 |  |  |
| ANXA6 |  |  |
| TCEB2 |  |  |
| IGLV1-40 |  |  |
| ROCK2 |  |  |
| IL36G |  |  |
| MFAP4 |  |  |
| RCC1 |  |  |
| TAAR2 |  |  |
| KLHDC8B |  |  |
| TSPAN9 |  |  |
| FLNB |  |  |
| PRKAR2A |  |  |
| QDPR |  |  |
| TRIP6 |  |  |
| NUP214 |  |  |
| HARS |  |  |
| UCKL1 |  |  |
| XYLB |  |  |
| LIN7A |  |  |
| PUS7 |  |  |
| TMEM59 |  |  |
| CBR3 |  |  |
| PFDN2 |  |  |
| MTMR10 |  |  |
| MAP4K5 |  |  |
| RNF123 |  |  |
| RAN |  |  |
| UHRF1 |  |  |
| COPB1 |  |  |
| PTPN23 |  |  |
| GIPC2 |  |  |
| PLEC |  |  |
| DSTN |  |  |
| BROX |  |  |
| CNGB1 |  |  |
| DHX36 |  |  |
| CSNK1A1 |  |  |
| RPS2P55 |  |  |
| GDI2 |  |  |
| DSG2 |  |  |
| ASNS |  |  |
| GNL3 |  |  |
| RTCB |  |  |
| PRKCI |  |  |
| EIF2B5 |  |  |
| PDLIM5 |  |  |
| PTRF |  |  |
| EXOSC5 |  |  |
| ACTR3 |  |  |
| COLGALT1 |  |  |
| ELP3 |  |  |
| GATAD2B |  |  |
| SCLT1 |  |  |
| BHMT |  |  |
| RPS16P10 |  |  |
| TENM4 |  |  |
| DOCK5 |  |  |
| ZBED9 |  |  |
| UBE2K |  |  |
| RPL35A |  |  |
| PIGK |  |  |
| AFM |  |  |
| FBXL8 |  |  |
| COL13A1 |  |  |
| ZCCHC8 |  |  |
| TTYH3 |  |  |
| DKC1 |  |  |
| PPP1CB |  |  |
| RAE1 |  |  |
| HSPE1 |  |  |
| PLEKHO2 |  |  |
| VPS39 |  |  |
| UBA6 |  |  |
| ADD3 |  |  |
| DNAJB4 |  |  |
| SMAD3 |  |  |
| EIF3D |  |  |
| ARHGAP5 |  |  |
| TUBGCP2 |  |  |
| CYB5R2 |  |  |
| NDST1 |  |  |
| DCPS |  |  |
| LYN |  |  |
| ANAPC5 |  |  |
| VPS52 |  |  |
| HADHA |  |  |
| DAB2IP |  |  |
| RAB4A |  |  |
| TLDC1 |  |  |
| ITFG3 |  |  |
| VPS8 |  |  |
| RNF149 |  |  |
| TTC39C |  |  |
| ABR |  |  |
| TCAF1 |  |  |
| ALDOC |  |  |
| AMPD3 |  |  |
| INHBA |  |  |
| RCN1 |  |  |
| SMU1 |  |  |
| VTI1A |  |  |
| NT5C |  |  |
| TRAF2 |  |  |
| TIMP2 |  |  |
| CTTN |  |  |
| GORASP2 |  |  |
| AEBP1 |  |  |
| TOR4A |  |  |
| RPL31 |  |  |
| LAMP2 |  |  |
| PCDH10 |  |  |
| TIGAR |  |  |
| IDH2 |  |  |
| ESM1 |  |  |
| PA2G4 |  |  |
| CHD3 |  |  |
| RPL38 |  |  |
| CTPS2 |  |  |
| SELENBP1 |  |  |
| INF2 |  |  |
| RDH11 |  |  |
| LTN1 |  |  |
| RPS16 |  |  |
| ADGRL2 |  |  |
| IDH1 |  |  |
| HIST1H4E |  |  |
| CTU2 |  |  |
| MESDC2 |  |  |
| PGD |  |  |
| PZP |  |  |
| ADAR |  |  |
| PPME1 |  |  |
| MDN1 |  |  |
| ERLIN2 |  |  |
| LRSAM1 |  |  |
| PGM2L1 |  |  |
| OGFRL1 |  |  |
| SPTAN1 |  |  |
| HTT |  |  |
| NTMT1 |  |  |
| STK39 |  |  |
| RBP4 |  |  |
| TPBG |  |  |
| ATP6V1C1 |  |  |
| AHSA1 |  |  |
| ALG2 |  |  |
| AVL9 |  |  |
| RAD50 |  |  |
| CRIPT |  |  |
| PACSIN2 |  |  |
| PSMA1 |  |  |
| SRGN |  |  |
| PARK7 |  |  |
| LARP6 |  |  |
| JUP |  |  |
| STXBP2 |  |  |
| BCAT1 |  |  |
| ALDH2 |  |  |
| WNT7B |  |  |
| HIST2H2BE |  |  |
| IRF2BP1 |  |  |
| DDOST |  |  |
| SEC31A |  |  |
| VAT1L |  |  |
| KIF1A |  |  |
| MOB3A |  |  |
| HAT1 |  |  |
| ASAP1 |  |  |
| ELP2 |  |  |
| SLC25A5 |  |  |
| PRAF2 |  |  |
| TBL3 |  |  |
| NOMO3 |  |  |
| TGFBRAP1 |  |  |
| ARHGEF12 |  |  |
| TRIP4 |  |  |
| NOMO2 |  |  |
| MYO5A |  |  |
| CCT5 |  |  |
| HIST1H1D |  |  |
| MAP2K6 |  |  |
| AP2S1 |  |  |
| SSFA2 |  |  |
| PRMT1 |  |  |
| MAP3K5 |  |  |
| FAM120C |  |  |
| DCAF8 |  |  |
| GLRX3 |  |  |
| PKM |  |  |
| NIT1 |  |  |
| ALCAM |  |  |
| DCTN3 |  |  |
| MST1 |  |  |
| USP28 |  |  |
| OTUB1 |  |  |
| PRDX5 |  |  |
| NOP58 |  |  |
| VDAC3 |  |  |
| ACVR1 |  |  |
| RFTN1 |  |  |
| WNT5B |  |  |
| TIMP1 |  |  |
| UBXN1 |  |  |
| RUVBL2 |  |  |
| SPIDR |  |  |
| NGLY1 |  |  |
| INPP4A |  |  |
| HSD17B10 |  |  |
| BZW2 |  |  |
| SLC25A10 |  |  |
| SRI |  |  |
| STX7 |  |  |
| PRPSAP2 |  |  |
| TCP1 |  |  |
| CDC34 |  |  |
| HRNR |  |  |
| DYNLT1 |  |  |
| TTC38 |  |  |
| IARS |  |  |
| AP4S1 |  |  |
| LUC7L2 |  |  |
| EFTUD1 |  |  |
| VIPAS39 |  |  |
| SRP14 |  |  |
| NDUFS2 |  |  |
| PARP10 |  |  |
| HIP1 |  |  |
| IGLV2-11 |  |  |
| FAHD2A |  |  |
| WDR4 |  |  |
| AKT1 |  |  |
| SERPINF1 |  |  |
| KRT73 |  |  |
| LLGL1 |  |  |
| CALD1 |  |  |
| TFCP2 |  |  |
| ARAP1 |  |  |
| AIP |  |  |
| RPS11 |  |  |
| CHST14 |  |  |
| SNX17 |  |  |
| TRA2A |  |  |
| AHNAK |  |  |
| PDIA3 |  |  |
| MTMR12 |  |  |
| ANXA11 |  |  |
| AKR1C1 |  |  |
| TTC21B |  |  |
| DPYSL2 |  |  |
| DYNC1LI1 |  |  |
| CORO1B |  |  |
| PIP4K2A |  |  |
| LHFPL2 |  |  |
| VIL1 |  |  |
| RPS27 |  |  |
| MYH9 |  |  |
| HGFAC |  |  |
| RAB5B |  |  |
| DPY30 |  |  |
| MAGOH |  |  |
| BRCC3 |  |  |
| COPG2 |  |  |
| STX3 |  |  |
| SNRNP40 |  |  |
| TERF1P5 |  |  |
| FN1 |  |  |
| ELMO2 |  |  |
| UFC1 |  |  |
| RNASEH2C |  |  |
| SF3B1 |  |  |
| WARS |  |  |
| LOXL3 |  |  |
| PPP2R5A |  |  |
| RPL14 |  |  |
| CCL20 |  |  |
| STX4 |  |  |
| CDK5 |  |  |
| SRPX |  |  |
| IGLC1 |  |  |
| PPP2R1A |  |  |
| PSMA5 |  |  |
| EML3 |  |  |
| SMARCE1 |  |  |
| RPS4XP13 |  |  |
| FOXK1 |  |  |
| QSOX1 |  |  |
| MGAT5 |  |  |
| UAP1 |  |  |
| BTF3 |  |  |
| HMGB1 |  |  |
| RABGGTA |  |  |
| APRT |  |  |
| C9orf91 |  |  |
| HNRNPH1 |  |  |
| RPL15P22 |  |  |
| TCEB1 |  |  |
| WDR7 |  |  |
| MAP2K4 |  |  |
| IL22RA1 |  |  |
| GDF9 |  |  |
| OBFC1 |  |  |
| MBLAC2 |  |  |
| DDX43 |  |  |
| TBC1D15 |  |  |
| LPP |  |  |
| DYNC2H1 |  |  |
| RP2 |  |  |
| SLC2A3 |  |  |
| SPON2 |  |  |
| PARP12 |  |  |
| TOM1 |  |  |
| SH3PXD2B |  |  |
| TNFRSF1A |  |  |
| C16orf13 |  |  |
| TREM1 |  |  |
| PRR4 |  |  |
| RAB15 |  |  |
| ANTXR2 |  |  |
| UFSP2 |  |  |
| FMNL2 |  |  |
| CRKL |  |  |
| IGHG1 |  |  |
| EHBP1 |  |  |
| TUBB4B |  |  |
| PHGDH |  |  |
| PSMA3 |  |  |
| KRT13 |  |  |
| PPP3CA |  |  |
| CPNE1 |  |  |
| UBASH3B |  |  |
| RQCD1 |  |  |
| PVR |  |  |
| GNAL |  |  |
| UBE2L3 |  |  |
| FAT4 |  |  |
| PSMB5 |  |  |
| IDE |  |  |
| DDX17 |  |  |
| MBNL1 |  |  |
| HEATR5A |  |  |
| MICAL2 |  |  |
| CD47 |  |  |
| MAGED1 |  |  |
| GPR176 |  |  |
| VPS28 |  |  |
| PTX3 |  |  |
| GPC1 |  |  |
| APOH |  |  |
| IKBIP |  |  |
| PCYT2 |  |  |
| PTK7 |  |  |
| NEK10 |  |  |
| FLNC |  |  |
| LYPLAL1 |  |  |
| KIDINS220 |  |  |
| BIRC3 |  |  |
| EEFSEC |  |  |
| NCL |  |  |
| HEATR3 |  |  |
| CORO2B |  |  |
| NEO1 |  |  |
| NDN |  |  |
| RAP2C |  |  |
| IGKV1-5 |  |  |
| TUBA1B |  |  |
| GPC4 |  |  |
| PLD3 |  |  |
| DDX47 |  |  |
| SERPINA1 |  |  |
| MED16 |  |  |
| CAMK2G |  |  |
| GYS1 |  |  |
| IKBKB |  |  |
| NADSYN1 |  |  |
| SARM1 |  |  |
| TBC1D17 |  |  |
| ITM2B |  |  |
| DCN |  |  |
| PFKFB3 |  |  |
| GATAD2A |  |  |
| PREP |  |  |
| AGL |  |  |
| RPLP1 |  |  |
| GMPS |  |  |
| GAN |  |  |
| NDUFV1 |  |  |
| CTNND2 |  |  |
| UQCRC2 |  |  |
| APOA4 |  |  |
| ADAM17 |  |  |
| ITGA2 |  |  |
| USP14 |  |  |
| ADARB1 |  |  |
| CCL28 |  |  |
| PACSIN3 |  |  |
| RPS2P51 |  |  |
| SMG1 |  |  |
| H2AFY |  |  |
| ATM |  |  |
| COPS8 |  |  |
| ADGRG1 |  |  |
| RPS27AP12 |  |  |
| EFEMP2 |  |  |
| RANGAP1 |  |  |
| PSMD2 |  |  |
| NME7 |  |  |
| WDR44 |  |  |
| ACTN2 |  |  |
| RPL12P2 |  |  |
| MAP2K1 |  |  |
| ALDH8A1 |  |  |
| TGFB1I1 |  |  |
| AP2B1 |  |  |
| STOML3 |  |  |
| ZNFX1 |  |  |
| FBLN1 |  |  |
| IGHV3-11 |  |  |
| RPA2 |  |  |
| EPB41 |  |  |
| VPS35 |  |  |
| ASCC3 |  |  |
| PBDC1 |  |  |
| ERCC2 |  |  |
| PGLYRP2 |  |  |
| HNRNPD |  |  |
| ADGRG4 |  |  |
| CCT3 |  |  |
| RBM3 |  |  |
| POLR2B |  |  |
| RDH5 |  |  |
| SLC12A4 |  |  |
| COMMD10 |  |  |
| IGLL5 |  |  |
| RPL10 |  |  |
| NEFH |  |  |
| RPL37A |  |  |
| IDH3B |  |  |
| TAGLN2 |  |  |
| SAMD9 |  |  |
| CAP1 |  |  |
| PDGFRA |  |  |
| FLII |  |  |
| SEC24B |  |  |
| AKR1B10 |  |  |
| OAS3 |  |  |
| DNAJB1 |  |  |
| RAB1A |  |  |
| YES1 |  |  |
| DRG1 |  |  |
| PFDN6 |  |  |
| ARL1 |  |  |
| DBF4B |  |  |
| HERC1 |  |  |
| SNRNP70 |  |  |
| UBE4B |  |  |
| NCAPD2 |  |  |
| HACE1 |  |  |
| CSNK2B |  |  |
| ASAP2 |  |  |
| PPP1R12C |  |  |
| LCN1P1 |  |  |
| DNAJC7 |  |  |
| NUDC |  |  |
| HPX |  |  |
| CLCN6 |  |  |
| TECR |  |  |
| EXOC4 |  |  |
| MVB12B |  |  |
| RAB21 |  |  |
| TOMM70A |  |  |
| GNG2 |  |  |
| PLA2G4A |  |  |
| DERA |  |  |
| GSN |  |  |
| EIF2B1 |  |  |
| RPLP0P2 |  |  |
| DNM2 |  |  |
| PSMB7 |  |  |
| TUBGCP5 |  |  |
| PWP2 |  |  |
| NONO |  |  |
| GCC1 |  |  |
| TSG101 |  |  |
| PBRM1 |  |  |
| CAPN7 |  |  |
| CSNK1D |  |  |
| HRG |  |  |
| MYO3B |  |  |
| EIF2AK4 |  |  |
| STRIP1 |  |  |
| DKK1 |  |  |
| TANC1 |  |  |
| PI4KB |  |  |
| DGKA |  |  |
| TELO2 |  |  |
| ASH1L |  |  |
| C6 |  |  |
| SLC7A10 |  |  |
| TUBB2A |  |  |
| ERGIC2 |  |  |
| NEK9 |  |  |
| SERPINB6 |  |  |
| EIF2S3 |  |  |
| CPSF3L |  |  |
| ATP2C1 |  |  |
| TRMT112 |  |  |
| MCM5 |  |  |
| DCAF7 |  |  |
| N6AMT1 |  |  |
| HSPA14 |  |  |
| WDR11 |  |  |
| EFTUD2 |  |  |
| P4HB |  |  |
| ANKFY1 |  |  |
| ATP5A1 |  |  |
| TTI1 |  |  |
| JMJD6 |  |  |
| PDIA5 |  |  |
| KIAA1549L |  |  |
| RPL29P9 |  |  |
| FGG |  |  |
| ATXN1 |  |  |
| PIEZO1 |  |  |
| SNRPD1 |  |  |
| USP24 |  |  |
| PTBP1 |  |  |
| SLC1A4 |  |  |
| SUPT5H |  |  |
| RPL26 |  |  |
| LRRK1 |  |  |
| KLHL22 |  |  |
| PPIB |  |  |
| PSMD10 |  |  |
| GARS |  |  |
| NPHP3 |  |  |
| RPS27AP16 |  |  |
| LMAN2 |  |  |
| BYSL |  |  |
| PPP1R13L |  |  |
| PABPC1 |  |  |
| TMEM132A |  |  |
| PPP1R8 |  |  |
| PSMC6 |  |  |
| RB1CC1 |  |  |
| C5 |  |  |
| FYCO1 |  |  |
| F11R |  |  |
| KRT28 |  |  |
| NAGK |  |  |
| LCN2 |  |  |
| GLCE |  |  |
| DDX5 |  |  |
| MCM3 |  |  |
| ACTG2 |  |  |
| GRM7 |  |  |
| KIF9 |  |  |
| ABCC4 |  |  |
| TRIM25 |  |  |
| MVD |  |  |
| AFAP1 |  |  |
| ABCC1 |  |  |
| RPSAP19 |  |  |
| MPP5 |  |  |
| GFPT1 |  |  |
| EEF1G |  |  |
| VPS33B |  |  |
| NUDT2 |  |  |
| MC1R |  |  |
| KIF2A |  |  |
| FAM3C |  |  |
| GSDMD |  |  |
| SLAIN1 |  |  |
| USP39 |  |  |
| ACAT1 |  |  |
| NELFB |  |  |
| HNRNPU |  |  |
| HNRNPM |  |  |
| FAM120A |  |  |
| PLAT |  |  |
| CANX |  |  |
| IQSEC1 |  |  |
| GJC1 |  |  |
| LAMA5 |  |  |
| NSDHL |  |  |
| IGKC |  |  |
| PYROXD1 |  |  |
| MTHFD1 |  |  |
| SLC39A14 |  |  |
| EXOSC9 |  |  |
| SMG8 |  |  |
| MTMR14 |  |  |
| MYO16 |  |  |
| ANKRD52 |  |  |
| MYO10 |  |  |
| SRP72 |  |  |
| ACTR2 |  |  |
| ARL6 |  |  |
| HSPB1 |  |  |
| PPWD1 |  |  |
| HIP1R |  |  |
| ADAD1 |  |  |
| THRAP3 |  |  |
| HIST1H4A |  |  |
| SF3B3 |  |  |
| CDK19 |  |  |
| PPP2R5D |  |  |
| TNFRSF11B |  |  |
| NRBP1 |  |  |
| SASS6 |  |  |
| BPIFA1 |  |  |
| CFAP20 |  |  |
| INTS5 |  |  |
| PARVA |  |  |
| NME3 |  |  |
| PSMG3 |  |  |
| EIF5A |  |  |
| HLA-B |  |  |
| ANAPC4 |  |  |
| RPS2P5 |  |  |
| ENAH |  |  |
| RTCA |  |  |
| TBCD |  |  |
| ST3GAL2 |  |  |
| GEMIN5 |  |  |
| TSC2 |  |  |
| GRB2 |  |  |
| HIST1H1E |  |  |
| VCAN |  |  |
| YBX1 |  |  |
| AP3B1 |  |  |
| AGAP3 |  |  |
| PITPNM1 |  |  |
| AOX1 |  |  |
| HSPA1B |  |  |
| ATP5I |  |  |
| CSNK2A1 |  |  |
| DYNC1H1 |  |  |
| RAB7A |  |  |
| G3BP1 |  |  |
| KPNA6 |  |  |
| MFI2 |  |  |
| RBFOX1 |  |  |
| PANK2 |  |  |
| TSPAN6 |  |  |
| PRSS12 |  |  |
| FHOD1 |  |  |
| GSTA1 |  |  |
| PRKD3 |  |  |
| NTM |  |  |
| SRP68 |  |  |
| WDR20 |  |  |
| FPGS |  |  |
| PPP1R12A |  |  |
| MAPK14 |  |  |
| FUBP3 |  |  |
| XYLT2 |  |  |
| UROD |  |  |
| PFKM |  |  |
| MAN2B1 |  |  |
| EEF1A1 |  |  |
| SLC26A11 |  |  |
| PGM2 |  |  |
| STK24 |  |  |
| COL8A2 |  |  |
| IQSEC2 |  |  |
| FDFT1 |  |  |
| THOP1 |  |  |
| AHSG |  |  |
| ABHD4 |  |  |
| IPO8 |  |  |
| FRMD6 |  |  |
| WDTC1 |  |  |
| THBS1 |  |  |
| TNS3 |  |  |
| CTNNAL1 |  |  |
| SRRT |  |  |
| MTAP |  |  |
| IGF2BP1 |  |  |
| ATP5J2 |  |  |
| PTPRM |  |  |
| MME |  |  |
| CLASP2 |  |  |
| KRT80 |  |  |
| EIF2B3 |  |  |
| HNRNPH2 |  |  |
| TM9SF3 |  |  |
| CTSD |  |  |
| ZPR1 |  |  |
| IGHM |  |  |
| STX12 |  |  |
| AP1G1 |  |  |
| TOR1B |  |  |
| SNX33 |  |  |
| HIST1H2BD |  |  |
| SELO |  |  |
| PLXNB1 |  |  |
| BDNF |  |  |
| NUSAP1 |  |  |
| RPL18 |  |  |
| SEC24D |  |  |
| CTNNA1 |  |  |
| OXNAD1 |  |  |
| DIMT1 |  |  |
| DNAJC13 |  |  |
| ABHD17A |  |  |
| CCDC17 |  |  |
| CST4 |  |  |
| MYOM2 |  |  |
| MDH1 |  |  |
| CAPZA1 |  |  |
| HINT1 |  |  |
| STAM2 |  |  |
| DOPEY2 |  |  |
| NEK7 |  |  |
| PON1 |  |  |
| AP3M1 |  |  |
| PARD3B |  |  |
| GLMP |  |  |
| CSTF3 |  |  |
| CPN1 |  |  |
| CTSK |  |  |
| DHRS7 |  |  |
| LAMA4 |  |  |
| TMEM30A |  |  |
| ECH1 |  |  |
| ALDH7A1 |  |  |
| MVP |  |  |
| SYNPO |  |  |
| SDC1 |  |  |
| SLC7A2 |  |  |
| CCM2 |  |  |
| PRPF6 |  |  |
| ATAD3A |  |  |
| RNH1 |  |  |
| GNG12 |  |  |
| ITGA1 |  |  |
| LMNA |  |  |
| PDCD6 |  |  |
| MALT1 |  |  |
| MCMBP |  |  |
| POLN |  |  |
| XYLT1 |  |  |
| NRD1 |  |  |
| DOCK1 |  |  |
| CLASP1 |  |  |
| ASL |  |  |
| RPL12P6 |  |  |
| CAD |  |  |
| HPR |  |  |
| HIST1H4L |  |  |
| CYBRD1 |  |  |
| UBP1 |  |  |
| NANS |  |  |
| L3HYPDH |  |  |
| PRPF39 |  |  |
| CTSA |  |  |
| APAF1 |  |  |
| RPS13 |  |  |
| BAX |  |  |
| STAT6 |  |  |
| NUP43 |  |  |
| B4GALT1 |  |  |
| LRRC47 |  |  |
| SSRP1 |  |  |
| UBL5 |  |  |
| FRYL |  |  |
| MINA |  |  |
| UBE2O |  |  |
| TBC1D10B |  |  |
| GPI |  |  |
| COL7A1 |  |  |
| PSMD11 |  |  |
| BPNT1 |  |  |
| CSF3 |  |  |
| PPA1 |  |  |
| DCD |  |  |
| PARVB |  |  |
| SAR1A |  |  |
| RPS6KA3 |  |  |
| NANP |  |  |
| DNPH1 |  |  |
| CAPNS1 |  |  |
| MAN2C1 |  |  |
| TNPO3 |  |  |
| SNRPD3 |  |  |
| LRCH3 |  |  |
| ECHS1 |  |  |
| DHX15 |  |  |
| ACTBL2 |  |  |
| PRPF4 |  |  |
| RBM12 |  |  |
| NAMPT |  |  |
| ETF1 |  |  |
| STRN3 |  |  |
| RPS2P8 |  |  |
| ATP1A1 |  |  |
| CALM1 |  |  |
| ERP29 |  |  |
| KCNMA1 |  |  |
| SLC16A1 |  |  |
| LIF |  |  |
| RFC5 |  |  |
| PFDN5 |  |  |
| MAPRE2 |  |  |
| CHD4 |  |  |
| RAB3GAP2 |  |  |
| COMMD2 |  |  |
| LRIG1 |  |  |
| USP47 |  |  |
| ATXN2 |  |  |
| HSP90AB2P |  |  |
| RPS5 |  |  |
| UNC13B |  |  |
| KRT84 |  |  |
| MROH1 |  |  |
| IGHG4 |  |  |
| APOBEC3C |  |  |
| MTHFR |  |  |
| KRT9 |  |  |
| PAFAH1B2 |  |  |
| TAOK3 |  |  |
| FHL2 |  |  |
| ESYT2 |  |  |
| ACTN3 |  |  |
| HIST1H1C |  |  |
| MYO6 |  |  |
| HMGCL |  |  |
| LRP12 |  |  |
| LPIN1 |  |  |
| FKRP |  |  |
| COMMD9 |  |  |
| EIF5 |  |  |
| BUD31 |  |  |
| RPL17 |  |  |
| VPS29 |  |  |
| MEP1A |  |  |
| CCAR1 |  |  |
| DLG1 |  |  |
| ITM2C |  |  |
| FAM98A |  |  |
| TM7SF3 |  |  |
| LGALS3BP |  |  |
| RAB18 |  |  |
| VAMP3 |  |  |
| ARF1 |  |  |
| PCDH18 |  |  |
| SERPINA3 |  |  |
| IGHV3-7 |  |  |
| USP11 |  |  |
| HIST1H2AE |  |  |
| SSH3 |  |  |
| CSDE1 |  |  |
| HSPA1A |  |  |
| HMOX1 |  |  |
| TNPO1 |  |  |
| ARG1 |  |  |
| FBXL20 |  |  |
| CCT8 |  |  |
| SPRY2 |  |  |
| PRDX3 |  |  |
| CRTAP |  |  |
| MAP4 |  |  |
| GOLGA8CP |  |  |
